# Italian Odonates in the Pandora’s Box: A Comprehensive DNA Barcoding Inventory Shows Taxonomic Warnings at the Holarctic Scale

**DOI:** 10.1101/2020.04.23.056911

**Authors:** Andrea Galimberti, Giacomo Assandri, Davide Maggioni, Fausto Ramazzotti, Daniele Baroni, Gaia Bazzi, Ivan Chiandetti, Andrea Corso, Vincenzo Ferri, Mirko Galuppi, Luca Ilahiane, Gianandrea La Porta, Lorenzo Laddaga, Federico Landi, Fabio Mastropasqua, Samuele Ramellini, Roberto Santinelli, Giovanni Soldato, Salvatore Surdo, Maurizio Casiraghi

**Affiliations:** ZooPlantLab, Department of Biotechnology and Biosciences, University of Milano - Bicocca, P.za Della Scienza 2, 20126-I Milan, Italy; Area per l’Avifauna Migratrice, Istituto Superiore per la Protezione e la Ricerca Ambientale (ISPRA), Via Ca’ Fornacetta 9, 40064-I Ozzano Emilia BO, Italy; Department of Environmental and Earth Sciences (DISAT), University of Milano - Bicocca, P.za Della Scienza 1, 20126-I Milan, Italy; Marine Research and High Education (MaRHE) Center, University of Milano - Bicocca, 12030 Faafu Magoodhoo, Maldives; Section of Ecology, Department of Biology, University of Turku, FI-20014 Turku, Finland; CROS Varenna, via Venini 17, 23829 Varenna (LC), Italy; Via Braide Podé 8, 33010 Colloredo di Monte Albano (UD), Italy; Via Camastra 10, 96100 Siracusa, Italy; Department of Biology, University of Rome 2 - Tor Vergata, Via della Ricerca Scientifica 1, 00133-I Rome, Italy; Via Federico Borromeo 24, 20822 Seveso (MB), Italy; Department of Sciences and Technological Innovation (DISIT), University of Eastern Piedmont, Viale Teresa Michel 11, 15121 Alessandria, Italy; Department of Chemistry, Biology and Biotechnology (DCBB), University of Perugia, Via Elce di Sotto 8, 06123 Perugia, Italy; Società di Scienze Naturali del Verbano Cusio Ossola, Natural Science Museum Collegio Mellerio Rosmini, Domodossola (VB), Italy; Via G. Mameli 14- I, 62100 Macerata, Italy; Association Centro Studi de Romita, Via G. Postiglione 9, 70126-I Bari, Italy; Department of Ecology and Environmental Policies, University of Milan, Via Celoria 26, 20133 Milan, Italy; via Alfieri 24, 20855 Lesmo (MB), Italy; Via Ormea 130, 10126 Torino, Italy; Department of Agriculture, Food and Forest Sciences, University of Palermo, Viale delle Scienze, 90128 Palermo, Italy

**Keywords:** Anisoptera, BOLD, Cryptic species, Odonata, Species delimitation, Zygoptera

## Abstract

The Odonata are considered among the most endangered freshwater faunal taxa. Their DNA-based monitoring relies on validated reference datasets that are often lacking or do not cover important biogeographical centres of diversification. This study presents the results of a DNA barcoding campaign on Odonata, based on the standard 658 bp 5’ end region of the mitochondrial COI gene, involving the collection of 812 specimens (409 of which barcoded) from peninsular Italy and its main islands (328 localities), belonging to all the 88 species (31 Zygoptera and 57 Anisoptera) known from the country. Additional BOLD and GenBank data from Holarctic samples expanded the dataset to 1294 DNA barcodes. A multi-approach species delimitation analysis involving two distance (OT and ABGD) and four tree-based (PTP, MPTP, GMYC, bGMYC) methods were used to explore these data. Of the 88 investigated morphospecies, 75 (85%) unequivocally corresponded to distinct Molecular Operational Units, whereas the remaining ones were classified as ‘warnings’ (i.e., showing a mismatch between morphospecies assignment and DNA-based species delimitation). These results are in contrast with other DNA barcoding studies on Odonata showing up to 95% of identification success. The species causing warnings were grouped in three categories depending on if they showed low, high, or mixed genetic divergence patterns. The analysis of haplotype networks revealed unexpected intraspecific complexity at the Italian, Palearctic, and Holarctic scale, possibly indicating the occurrence of cryptic species. Overall, this study provides new insights into the taxonomy of odonates and a valuable basis for future DNA and eDNA-based monitoring studies.

## Introduction

Modern conservation efforts rely on the availability of regional or local comprehensive species inventories (Morinière et al., 2019; Weigand et al., 2019; Altermatt et al., 2020). However, in most cases, morphology alone makes it hard to complete such inventories, so that alternative taxonomic methods, based on technological and analytical advances, have become fundamental to support and integrate the study of biodiversity (Padial et al., 2010; DeSalle & Goldstein, 2019; Schmid-Egger et al., 2019). Since the early 2000s, DNA barcoding has become a reliable basis for assembling the reference sequence libraries necessary to identify specimens of known species, also enhancing species discovery in neglected or poorly investigated taxonomic groups (e.g., Hebert et al., 2003; Galimberti et al., 2012; Dapporto et al., 2019). Curated and comprehensive DNA barcode reference libraries such as the Barcode of Life Data System BOLD (Ratnasingham & Hebert, 2007; 2013) often allow fast and reliable species identification with a considerable saving in terms of time and resources when personnel and taxonomic expertise are limited.

DNA barcoding has certainly revolutionized modern taxonomy, but some caveats dealing for example with pseudogenes (Berthier et al., 2011), introgression, and incomplete lineage sorting phenomena may lead to misidentification or wrong taxonomic assumptions (Eberle et al., 2020). However, even considering these possible pitfalls, the use of a variety of sequence analysis methods and parameters can be adopted to detect warnings (i.e., mismatches between morphospecies assignment and DNA-based species delimitation) of possible taxonomic relevance from DNA barcoding data (De Salle & Goldstein, 2019; Matos-Maraví et al., 2019). Such warnings could be then used to plan subsequent research aimed at assessing presumptive new species detected with these methods (Carstens et al., 2013). For example, by integrating morphology, multi-locus genetics, ecology or other information, the molecularly defined species could be either confirmed or rejected (Kajitoch, Montagna, & Wanat, 2018; Dufresnes et al., 2019). On the other hand, morphological species that are inconsistently delimited at the molecular level could be designated for further analyses, such as geometric morphometrics (Solano et al., 2018) or population genomics (Dufresnes et al., 2020).

Another possible weakness of DNA barcoding regards the geographical distribution of the samples. Some studies observed a diminished identification accuracy due to the increasing intraspecific genetic divergence when a wider geographical sampling scale was considered (Meyer & Paulay 2005; Bergsten et al. 2012) or when genetic diversity hotspots are ignored or undersampled (Gaytán et al., 2020). The availability of reliable reference databases covering taxonomic diversity and including the whole intraspecific geographic variability of the considered taxa constitutes a keystone element for large biodiversity investigations associated with DNA and environmental DNA (eDNA) metabarcoding (Porter & Hajibabaei, 2018; Piper et al., 2019; Creedy et al., 2020).

In this context, freshwater ecosystems are gaining more and more attention concerning eDNA-based studies (Pawloski et al., 2018; Bush et al., 2019; Weigand et al., 2019) since they are globally threatened and there is an increasing need to improve the monitoring of their biodiversity and changes over time (Collen et al., 2014; Darwall et al., 2018). Among freshwater bioindicators, the Odonata is a group of growing social and scientific interest. It is a small order, by insect standards, counting about 6300 species worldwide (Schorr & Paulson, 2019); 145 species have been reported in Europe, with 51 and 94 species belonging to the suborders Zygoptera (damselflies) and Anisoptera (dragonflies), respectively (Boudot & Kalkman, 2015; Viganò, Janni, & Corso, 2017; Lopez-Estrada et al., 2020). Traditionally, odonate identification has relied on morphological data that could be biased by difficulties in larval/exuviae recognition, the occurrence of cryptic species, and even high introgression rates (Marinov 2001; Bried, D’Amico, & Samways, 2012; Solano et al., 2018; Bried & Hinchliffe, 2019). The integration of traditional field protocols with alternative strategies, such as DNA and eDNA-based approaches, may support fast and accurate species identification and monitoring of this insects order.

Although the Odonata, together with Trichoptera, Hemiptera, and crustaceans are the best-covered taxa in the BOLD System database (Weigand et al., 2019), the currently available data do not take into account possible biases due to the absence of representatives from areas of biogeographical/speciation interest (Gaytán et al., 2020), and to the absence of genetic information (including DNA barcoding data) for some species of conservation concern (Sahlén et al., 2010; Boudot & Kalman, 2015; Kalkman et al., 2018). Europe can be considered the cradle of odonate systematics, but the evolutionary history and radiation of many genera, especially in Mediterranean glacial refugia, is still unresolved (e.g., *Calopteryx, Ischnura* and *Onychogomphus*, see Dijkstra & Kalkman, 2012). In the Western Mediterranean, at least three areas have been recognized as distinct glacial refugia during Pleistocene glaciations: Iberia, peninsular Italy (and its islands) and the Maghreb (Schmitt & Varga, 2012; Husemann et al 2013; Heiser, Dapporto, & Schmitt, 2014). Similarly, the Alps acted as a barrier and a refuge through climatically induced range changes, and for this reason are today recognized as a major geographical feature in shaping the phylogeography of European species (Hewitt, 2004). For this reason, the Italian peninsula, which is a natural bridge between the Mediterranean basin and the Alps, is a major biodiversity hotspot for odonates with 89 breeding species (Riservato et al., 2014a; http://www.odonata.it/libe-italiane/), of which 19 are listed as threatened or near-threatened in the Italian Red List (Riservato et al. 2014b) and 10 listed in the European “Habitat” Directive. Nevertheless, almost no genetic information is available in public databases for Italian odonates with less than 100 COI barcode sequences deposited there, mostly belonging to a few *Cordulegaster* and *Coenagrion* species. Thus, the claim that Italy has not played an important biogeographical and evolutionary role for the odonate diversification (Heiser & Schmitt, 2010) appears questionable, given the limited amount of molecular investigation occurred to date.

In order to increase the availability of genetic information on European Odonata, in this study we aimed at *i*) creating the first comprehensive and well-curated reference DNA barcoding library of Italian species, based on intense national sampling encompassing the main regions of biogeographical interest; *ii*) investigating the diversity and distribution of mitochondrial lineages of Italian Odonata using multiple species delimitation approaches, based on both character-explicit and distance-based methods (*sensu* DeSalle & Goldstein, 2019). Moreover, since almost all the species occurring in Italy are more widely distributed in Europe, Asia and North America, the same approach has also been performed at the Holarctic scale by creating a second comprehensive dataset including a mixture of Italian and non-Italian publicly available sequences.

## Materials and Methods

### Specimen collection, identification and archiving

During 2018-2019, a network of 20 professional and voluntary taxonomists (including the authors), involved in the “DNA barcoding Italian odonates” project, collected odonates in six macro-regions of Italy (see Figure 1). Such regions correspond to the main cradles of intraspecific genetic divergence previously identified for other taxa (e.g., Dapporto, 2010; Bernini et al., 2016; Wauters et al., 2017; Dufresnes et al., 2019; Scalercio et al., 2020).

**FIGURE 1:**
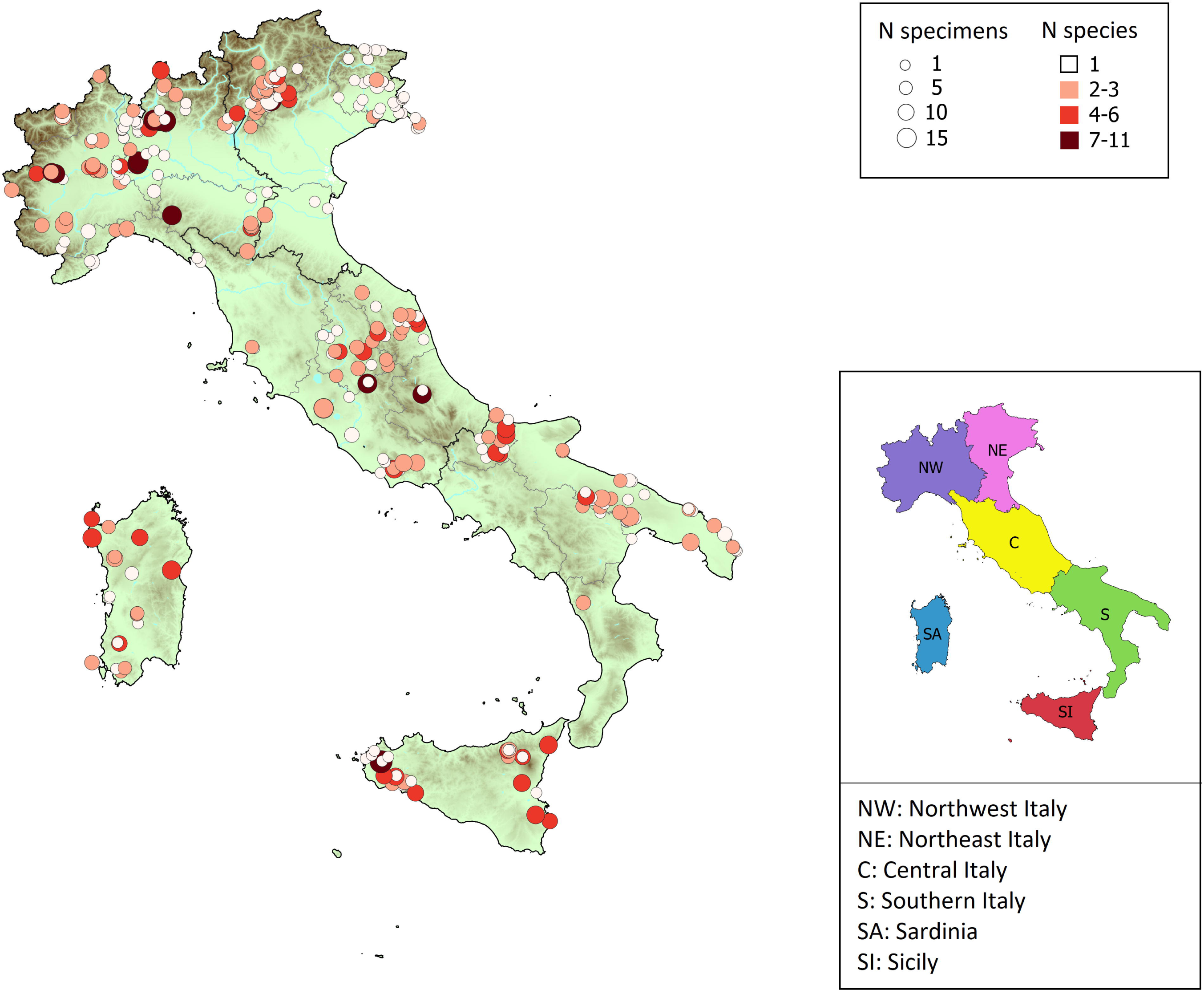
Collection sites for the sampled 88 odonate species included in the present study. The left map shows the number of specimens and the number of collected species per site indicated by circle size and colour respectively. The right map shows the borders of the macro-regions used for the analysis of genetic distances and structure of each species. Details of the 328 sites and collected and DNA barcoded specimens are provided in Appendix1. This map was created in QGIS version 3.4.13 (https://www.qgis.org/it/site/).

Sampling sites included 328 localities scattered in most of the Italian administrative regions and were mostly wet ecosystems belonging to protected or not protected areas from the Alpine (up to 2300 m a.s.l.), peninsular and insular (i.e., Sardinia and Sicily) regions. For the sampling of protected species, the Italian Ministry of the Environment issued a special permit (Prot. DPN 0031783.20-11-2019).

Only adults were sampled by hand netting and identified according to Dijkstra & Lewington (2006). In some cases, especially for damselflies belonging to *Coenagrion* and *Chalcolestes* genera, fine characters diagnostic for species identification (e.g., pronotum, cerci and ovipositor) were checked using a stereomicroscope (Olympus SZX7). The nomenclature used in this study follows the last update of the World Odonata List (Schorr and Paulson, 2019). For each individual, one or two middle legs were stored in 96% EtOH before DNA extraction. All samples and most specimens were vouchered and stored in the MIB:ZPL collection (University of Milano - Bicocca, Milan, Italy) whereas the remaining specimens are conserved in GA private collection.

A total of 812 specimens belonging to 88 morphospecies (31 Zygoptera and 57 Anisoptera) were collected. Sampling localities and details are available in BOLD under the project code ZPLOD (https://www.boldsystems.org/index.php/MAS_Management_DataConsole?codes=ZPLOD) and are also reported in Appendix S1. Sampling location coordinates were given in decimal degrees (EPSG: 4326) and approximated at the first decimal unit (i.e., ≈ 11 km). Since many species (or biotopes) are of conservation concern, we chose not to give details on exact locations (Lunghi et al., 2019).

### DNA extraction, amplification, and sequencing

For each morphospecies, samples for genetic analysis were selected from distant sites to maximize the chance of observing intraspecific geographic variation. Overall, 409 specimens were selected for DNA barcoding characterization. DNA was extracted from a middle leg (Zygoptera) or a coxa (Anisoptera) using the Qiagen DNeasy Blood and Tissue Kit (Qiagen, Milan, Italy) following the manufacturer’s instructions.

For each sample, the standard DNA barcode 5’-end region of the mitochondrial COI gene (658 bp) was amplified with the M13-tailed primers ODOF1_t1 and ODOR1_t1 (Semotok, unpublished, Source: BOLD Systems primer database), previously used by Koroiva and co-workers (2017). In case of unsuccessful amplification, the alternative COI primers ODO_LCO1490d/ ODO_HCO2198d (Dijkstra et al., 2014) were adopted to amplify the selected region. In both cases, PCR amplification conditions were as follows: 94°C for 5 min, 35 cycles at 94°C for 60 s, 50°C for 90 s and 72°C for 60 s, and a final extension at 72°C for 7 min. PCRs were conducted as in Mazzamuto et al. (2016) and sequencing was performed bi-directionally at Eurofins Genomics (Milan, Italy). Standard M13F and M13R oligos were used for sequencing the amplicons obtained with the ODOF1_tl/ODOR1_tl primer pair. Consensus sequences were obtained by editing the electropherograms with Bioedit 7.2 (Hall, 1999). After primer trimming, the presence of open reading frame was verified for the obtained consensus sequences by using the on-line tool EMBOSS Transeq (http://www.ebi.ac.uk/Tools/st/emboss_transeq/), then sequences were aligned with MAFFT 7.110 (Katoh & Standley, 2013) using the *E-INS-i* option. Consensus sequences were deposited in the BOLD Systems (project code ZPLOD) and GenBank (accession numbers reported in Appendix S1 and Appendix S2).

### Sequence mining and datasets assembling

Orthologous COI barcode sequences, belonging to the species included in the Italian odonate list (Riservato et al., 2014a) were downloaded from public repositories (i.e., BOLD and GenBank). Sequences showing insertions/deletions, > 1% of missing sites or overlapping less than 500 bp with the COI region amplified and sequenced for the Italian samples, were discarded. Overall, a total of 885 COI barcode sequences were retrieved from public repositories. These nucleotide sequences and those obtained in the present study were grouped in two datasets: (i) dataset DS1, composed of the DNA barcoding sequences produced in this study on Italian samples (i.e., 409) plus 69 sequences from Italian samples of 10 odonate species analysed in previous studies (see Appendix S1); (ii) dataset DS2, composed of the sequences mined from online databases plus dataset DS1 and other COI barcode sequences obtained from additional samples retrieved by the authors in other western Palearctic countries (See Appendix S2). The two datasets were kept separated in order to evaluate first the efficiency of the multi-approach species delimitation and to detect taxonomic warnings (e.g., reduced interspecific genetic variability or high intraspecific structure at the COI barcode locus) within Italian reference specimens only (DS1). Then, to better investigate such warnings on a wider geographic scale and find new cases of possible taxonomic relevance, the same species delimitation strategy was adopted on sequence data from across the Holarctic range of the taxa occurring in Italy (DS2).

### Species delimitation analysis

Phylogenetic trees were reconstructed separately for Zygoptera and Anisoptera. First, identical sequences were collapsed into unique haplotypes (See Appendix S1 and Appendix S2) and appropriate substitution models were determined using JModelTest 2 (Darriba et al., 2012) using the Akaike Information Criterion (AIC), the corrected Akaike Information Criterion (AICc), and the Bayesian Information Criterion (BIC). For Anisoptera, the best-fitting substitution models were GTR+I+G (AIC, BIC) and HKY+I+G (AICc), whereas for Zygoptera GTR+I+G (AIC) and HKY+I+G (BIC, AICc), and the use of different models did not influence downstream analyses. Phylogenetic inference analyses were performed using two optimality criteria: Bayesian inference (BI) and maximum likelihood (ML). BI was conducted by using BEAST 1.8.2 (Drummond et al., 2012), setting a coalescent tree prior and an uncorrelated lognormal relaxed clock. Three replicate analyses were run for 10^8^ million generations with a sampling frequency of 10,000 and were combined using LogCombiner 1.8.2 (Drummond et al., 2012) with a burn-in set to 25%, after checking the stationarity for effective sampling size (ESS) and unimodal posterior distribution using Tracer v.1.6 (Rambaut et al., 2014), making sure ESS values were higher than 200 for all parameters. Maximum clade credibility trees were computed using TreeAnnotator 1.8.2 (Drummond et al., 2012). ML analyses were performed using RAxML 8.2.9 (Stamatakis, 2014) with 1000 bootstrap replicates, applying the GTR+I+G substitution model. Each sub-order dataset was rooted using two sequences belonging to the other dataset (*Nehalennia speciosa* (Charpentier, 1840) and *Pyrrhosoma nymphula* (Sulzer, 1776) for Anisoptera, and *Libellula quadrimaculata* Linnaeus, 1758 and *Lindenia tetraphylla* (Vander Linden, 1825) for Zygoptera). The obtained trees were visualised using FigTree v.1.4.3 (Rambaut, 2012) and customized using CorelDRAW X7. All analyses were run on CIPRES server (Miller, Pfeiffer, & Schwartz, 2010).

Distance-based and tree-based species delimitation approaches (*sensu* DeSalle & Goldstein, 2019) were employed to explore species boundaries in DS1, after removing the outgroups.

Distance-based methods rely on genetic distances to explore the presence of barcoding gaps and allow for species non-monophyly (Fontaneto, Flot, & Tang, 2015). In this work, we used the Automatic Barcoding Gap Discovery (ABGD; Puillandre et al., 2012) and the Optimal Threshold (OT) (Brown et al., 2012). ABGD delimitations were run on the website https://bioinfo.mnhn.fr/abi/public/abgd/. Parameters were set keeping Pmin = 0.001, Steps = 100, X = 1, Nb bins = 20, distance = Kimura 2-parameter, and varying Pmax between 0.01 to 0.1, in order to test the consistency of delimitations by changing the *a priori* maximum level of intraspecific distance. Only results obtained with Pmax = 0.1 have been reported. OT was calculated from the K2P genetic distance matrices with the ‘LocalMinima’ and ‘treshOpt’ functions of the SPIDER package (Brown et al., 2012) in R (R Core Team, 2019). In particular, OT is the genetic distance value that minimizes the cumulative identification error, that is the sum of false positive (no conspecific matches within the threshold of the query) and false negative (sequences from multiple species within the threshold) cases (Galimberti et al., 2012). Tree-based methods require species monophyly and are based on the analysis of branching rates (Fontaneto, Flot, & Tang, 2015). Here, the Poisson Tree Process (PTP; Zhang et al., 2013) and Generalized Mixed Yule-Coalescent (GMYC; Pons et al., 2006; Fujisawa & Barraclough, 2013) models were used. Single-threshold Bayesian PTP analyses (Zhang et al., 2013) were performed on the website http://species.h-its.org/ptp, running the analyses for 400,000 MCMC generations, with thinning value = 100 and burn-in = 0.25, and checking the trace file for convergence of the MCMC, whereas multiple-threshold PTP (MPTP) analyses (Kapli et al., 2017) were run on the website https://mptp.h-its.org. Single-threshold GMYC (Pons et al., 2006) and bGMYC (Reid & Carstens, 2012) analyses were run using BI ultrametric trees and were performed in R (R Core Team, 2019) using the packages SPLITS (Ezard, Tomochika, & Barraclough, 2009), APE (Paradis, Claude & Strimmer, 2004), and bGMYC (Reid & Carstens, 2012). bGMYC analyses were performed on a subset of 100 trees retrieved from the 10,000 trees obtained with each BI analysis, after assessing the convergence of the MCMC. Support values for the species delimitation hypotheses obtained with the Bayesian implementation of PTP and GMYC are shown in Appendix S3.

Apart from OT, all the other species delimitation approaches were independent, requiring no *a priori* information on the existing morphospecies. Overall, we considered a delimitation to be reliable when the different methods were mostly in agreement. The same pipeline was applied to the Zygoptera and Anisoptera COI sequences alignments of DS2. Conversely, OT values remained the same inferred from DS1 because reliable *a priori* information on morphospecies was available only for the reference samples collected in this study.

When mismatches occurred between delimitation based on DNA barcoding data and morphospecies assignment based on samples collected in this study (DS1) and/or inferred from public data (DS2), we identified a ‘warning’. Such mismatches have been grouped into three distinct categories (i.e., no interspecific-, intraspecific- and mixed-delimitation), depending on if they occur in spite of low, high or mixed genetic divergence patterns.

Considering the possible pitfalls in using a single mitochondrial marker to investigate species boundaries (Lohse, 2009; Petit & Excoffier, 2009), additional information from molecular repositories, literature and unpublished data from the entomologists participating in this study were considered to support the reliability of delimited species and legitimate the necessity of better investigating the detected warnings.

### Warnings investigation

To better support the main categories of warnings resulting from species delimitation analyses and to explore patterns of intraspecific geographic genetic variation, multi-locus genetic distance comparisons and/or haplotype networks were generated for selected species groups. In particular, public nucleotide sequences of the mitochondrial 16s rDNA and the nuclear AgT, PRMT and MLC genes were retrieved from GenBank for *Coenagrion mercuriale* (Charpentier, 1840), *C. puella* (Linnaeus, 1758), *C. pulchellum* (Vander Linden, 1825) and *C. ornatum* (Sélys, 1850). These nuclear markers were selected among the others as they were found to show a higher phylogeographic resolution in damselflies (Ferreira et al., 2014a). Genetic distances among species or lineages were computed using MEGA X (Kumar et al., 2018), and intra- and inter-clade uncorrected *p*-distances were calculated with 1000 bootstrap replicates.

In case of intraspecific delimitation, and when at least 20 COI barcode sequences were available from specimens collected in Italy and across the species Holarctic range, unrooted minimum spanning COI networks were obtained using the median-joining algorithm implemented in PopART 1.7 (Leigh & Bryant 2015).

## Results

### A reference DNA barcoding database of Italian odonates

We obtained high-quality full-length COI DNA barcode sequences (658 bp) for all the 409 selected specimens (see Appendix S1 for BOLD Process IDs and GenBank accession numbers), representing 88 morphospecies belonging to 36 genera and 10 families. No sequences contained insertions/deletions (indels), stop codons, or were biased by NUMT interference, with the only exception of one *Platycnemis pennipes* (Pallas, 1771) specimen (ZPLOD687-20) and one *Orthetrum cancellatum* (Linnaeus, 1758) specimen (ZPLOD613-20) with the primer pair ODOF1_t1/ODOR1_t1. Re-amplification and sequencing with the other primer set (i.e., ODO_LCO1490d/ODO_HCO2198d) allowed obtaining the correct DNA barcode sequence for these samples.

The number of newly barcoded specimens per species ranged from 1 to 34 (mean ± SD = 4.6 ± 4.6); 8 morphospecies were represented by a single accession (Appendix S1). The obtained DNA barcode sequences allowed to add two Italian (and Palearctic) species (i.e., *Orthetrum nitidinerve* (Sélys, 1841) and *Somatochlora meridionalis* Nielsen, 1935) not represented at all in the BOLD and GenBank databases, and novel haplotypes for almost all the investigated taxa. The DNA barcodes of DS1 were assigned to 83 distinct Barcode Index Numbers (BINs, Ratnasingham & Hebert, 2013) by BOLD and at least three of these (i.e., BOLD:AEC5518, BOLD:AEC4388, and BOLD:AEC4264 corresponding to *E. lindenii, L. dryas* and *O. nitidinerve*, respectively) were unique to the system at the moment of dataset submission (February 2020).

Considering the 69 Italian DNA barcode records mined from GenBank, a DNA barcoding reference library of 478 COI sequences from 88 species was established (i.e, DS1, see Appendix S1). The analysis of this dataset showed that genetic K2P-distance variation within morphospecies ranged from 0% to 9.17% (mean ± SD= 0.48 ± 0.62 %) in Zygoptera and from 0% to 2.64% (mean ± SD = 0.33 ± 0.29 %) in Anisoptera. Interspecific K2P-distance values ranged from 0% to 27.29% (mean ± SD= 19.41 ± 3.71%) and from 0% to 25.28% (mean ± SD= 18.25 ± 3.18 %) in Zygoptera and Anisoptera, respectively.

The distributions of intraspecific and interspecific K2P distances overlap, thus resulting in the absence of a complete barcode gap in DS1 (Appendix S4), similarly to what observed by Koroiva and Kvist (2018). Both ABGD and OT approaches identified a number of groups quite lower than the collected morphospecies (Table 1). In particular, the optimal threshold that minimises the number of false positive and false negative identifications resulted in 3.16% and 1.96% of K2P distance for Zygoptera and Anisoptera respectively, with associated cumulative errors 24.1% (57 sequences out of 237) and 9.96% (24 sequences out of 241) and mainly due to false negative cases. Overall, the Maximum intraspecific vs. Nearest Neighbour and Mean intraspecific vs. Nearest Neighbour analyses conducted in BOLD, confirmed the occurrence of a barcode gap for most investigated species groups (Appendix S5).

**TABLE 1:**
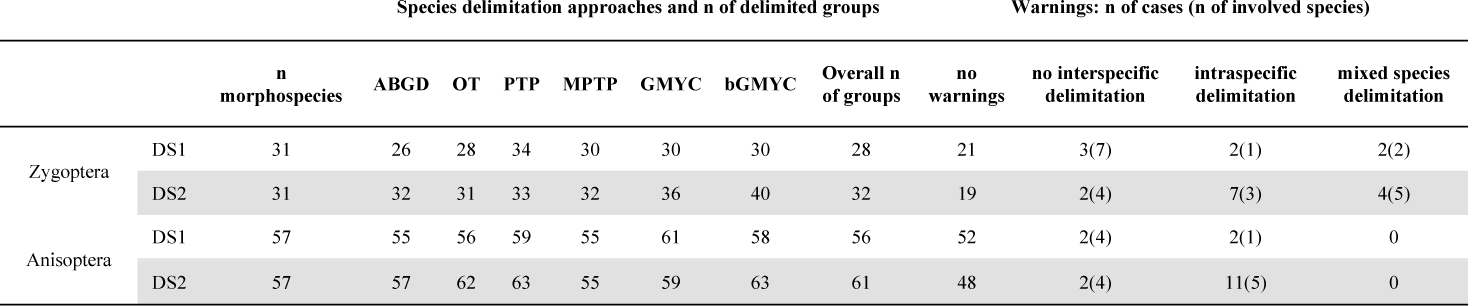
Species delimitation results. The table provides the number of groups delimited by each distance-based (ABGD and OT) and tree-based (PTP, MPTP, GMYC and bGMYC) approach conducted on the DNA barcode sequences of Italian odonate morphospecies grouped by suborder (Zygoptera and Anisoptera) and by DNA barcoding dataset (DS1 and DS2). The number of overall delimited groups, groups for each warning category and involved species (within brackets) are also reported.

We obtained further 816 DNA barcode records of the same species included in DS1 from BOLD and GenBank referred to Holarctic specimens. These were assembled in a comprehensive DNA barcoding inventory of 1294 COI sequences (i.e, DS2, see Appendix S2). In DS2, Genetic K2P-distance variation within morphospecies ranged from 0% to 8.81% (mean ± SD = 0.69% ± 0.78%) in Zygoptera and from 0% to 6.72% (mean ± SD = 0.59% ± 0.63%) in Anisoptera. Interspecific K2P-distance values ranged from 0% to 27.29% (mean ± SD = 19.51 ± 3.67%) and from 0% to 26.69% (mean ± SD = 18.09% ± 3.17%) in Zygoptera and Anisoptera, respectively. In contrast to DS1, the two genetic distance-based approaches (i.e., ABGD and OT) identified in DS2 a number of groups equal or quite higher than the collected morphospecies (Table 1), nevertheless not showing a complete barcode gap (Appendix S4). To better investigate this mismatch, multiple species delimitation approaches were conducted on both datasets.

### Species delimitation

Species delimitation analyses were conducted first on the reference DNA barcoding DS1 dataset and were based on separate phylogenetic trees encompassing, respectively, the damselfly and dragonfly morphospecies occurring in Italy (see Figure 2 and Figure 3). As highlighted in the trees and reported in Table 1, the overall number of groups delimited upon the criteria explained in Material and Methods, was quite lower than the number of recognized morphospecies (i.e., 28/31 in Zygoptera and 56/57 in Anisoptera). In general, the congruence between the delimited groups and the morphology-based identifications (‘no warnings’ in Table 1) was higher in Anisoptera, with 52 out of 57 (91.2%) cases, than in Zygoptera, with 21 out of 31 (67.7%) cases. Most of the warnings were due to reduced or no interspecific genetic diversity with seven and four morphospecies involved in Zygoptera and Anisoptera, respectively. Conversely, one case in both suborders was related to high genetic divergence at the intraspecific level (i.e., *Erythromma lindenii* (Sélys, 1840) and *Onychogomphus forcipatus* (Linnaeus, 1758)). Finally, in two damselfly species (i.e., *Chalcolestes viridis* (Vander Linden, 1825) and *C. parvidens* Artobolevskii, 1929), the pattern of genetic divergence was a mix of the other two warning categories with one sample (ZPLOD168-20), morphologically identified as *C. viridis*, included in the group of *C. parvidens* (see Figure 2, Figure 3 and Table 1).

**FIGURE 2:**
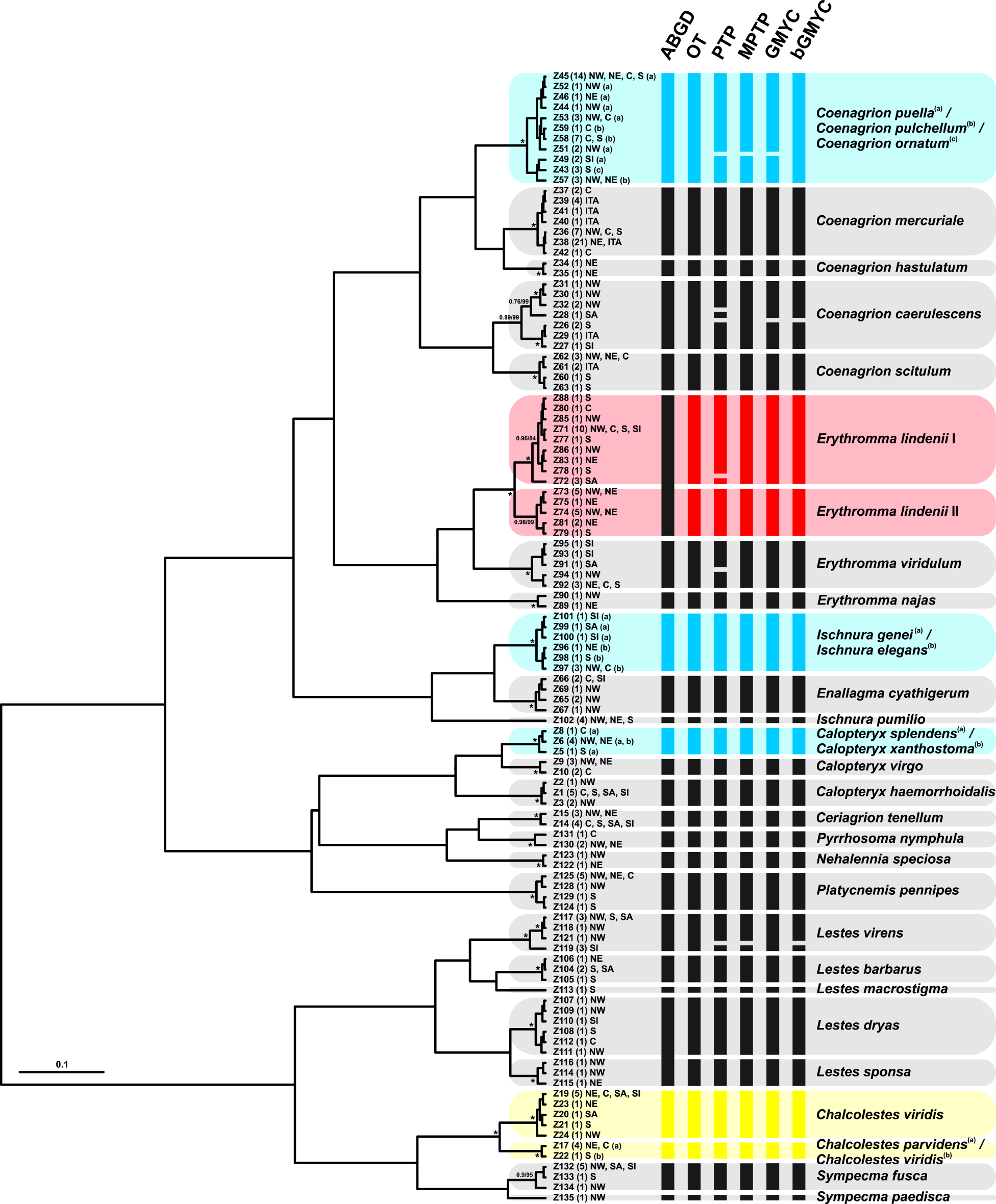
Multi-approach species delimitation of the 31 Italian Zygoptera morphospecies analyzed in this study, based on COI DNA barcode sequences from DS1. A Bayesian tree is used as a base to summarize the two threshold-based (OT and ABGD) and four character explicit-based (PTP, MPTP, GMYC, bGMYC) approaches. Specimen haplotype identifiers are reported on tips (see Appendix S1 for further details), the number of specimens sharing each haplotype is reported within brackets and abbreviations NW, NE, C, S, SI and SA correspond to the geographic haplotype origin as schematized in Figure 1 (ITA corresponds to unknown locality in Italy). Numbers above nodes represent BPP and BS, respectively, whereas asterisks correspond to maximal node support (BPP ≥ 0.99 and BS ≥ 95). Vertical colored solid boxes delimit putative species identified by the different approaches. Black: delimitation congruent with the identified morphospecies; Red: intraspecific delimitation; Light Blue: no interspecific delimitation; Yellow: Mixed species delimitation. In the case of intraspecific delimitation, each lineage has been numbered with Roman numerals. This tree was created with BEAST 1.8.2.

**FIGURE 3:**
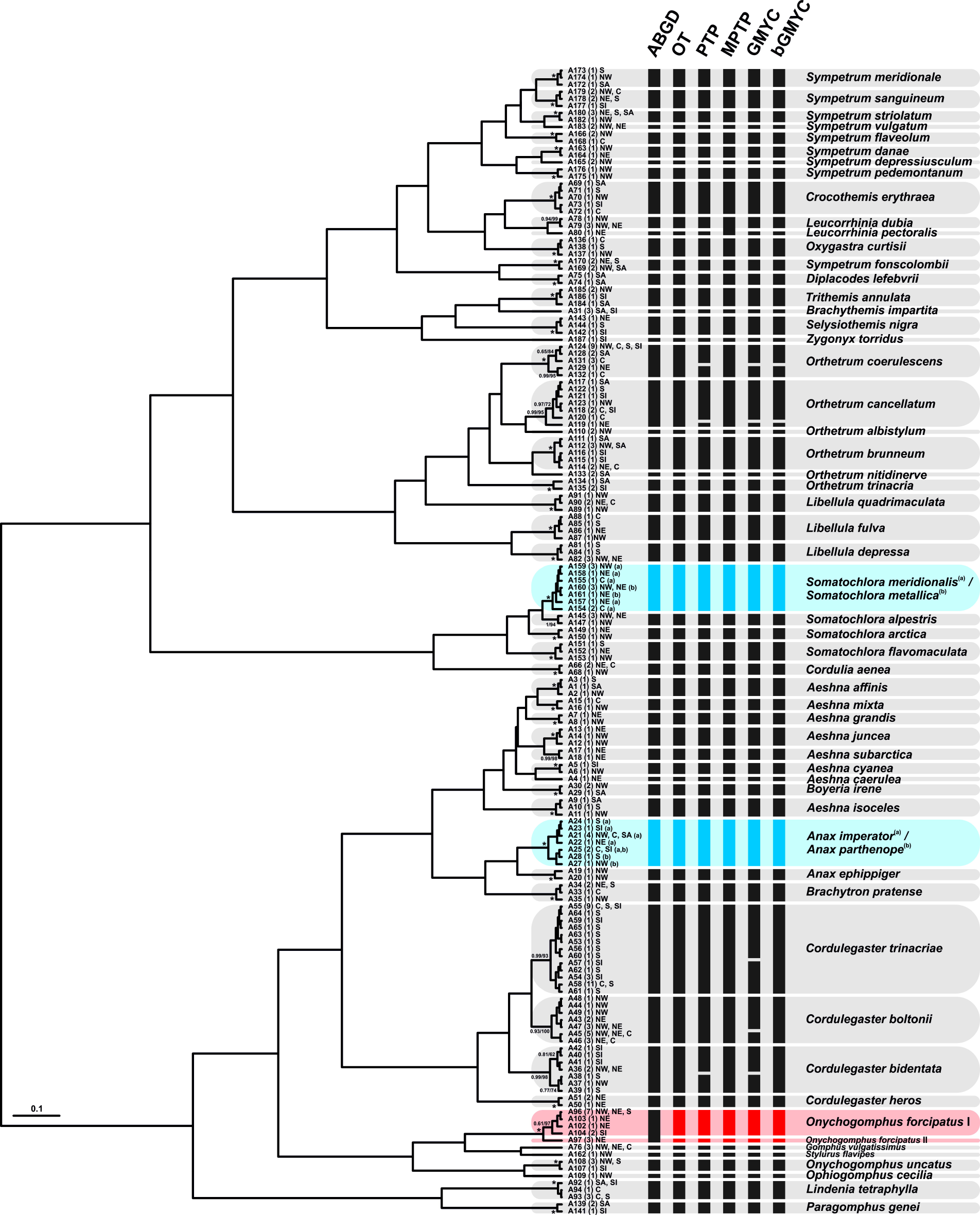
Multi-approach species delimitation of the 57 Italian Anisoptera morphospecies analyzed in this study, based on COI DNA barcode sequences from DS1. A Bayesian tree is used as a base to summarize the two threshold-based (OT and ABGD) and four character explicit-based (PTP, MPTP, GMYC, bGMYC) approaches. Specimen haplotype identifiers are reported on tips (see Appendix S1 for further details), the number of specimens sharing each haplotype is reported within brackets and abbreviations NW, NE, C, S, SI and SA correspond to the geographic haplotype origin as schematized in Figure 1. Numbers above nodes represent Bayesian posterior probabilities (BPP) and maximum likelihood bootstrap values (BS), respectively, whereas asterisks correspond to maximal node support (BPP ≥ 0.99 and BS ≥ 95). Vertical colored solid boxes delimit putative species identified by the different approaches. Black: delimitation congruent with the identified morphospecies; Red: intraspecific delimitation; Light Blue: no interspecific delimitation. In the case of intraspecific delimitation, each lineage has been numbered with Roman numerals. This tree was created with BEAST 1.8.2.

In contrast to DS1, the overall number of delimited groups in DS2 was higher than the number of investigated species (i.e., 32/31 in Zygoptera and 61/57 in Anisoptera). Overall, the warnings found in DS1 were almost all confirmed in DS2 (Table 1 and Appendix S6) and new 17 cases, belonging to 7 species (3 damselflies and 4 dragonflies) were found due to the new BOLD and GenBank DNA barcode sequences included in this dataset. Table 2 provides a list of the warning cases outlined in DS2.

**TABLE 2:** List of odonate species showing warnings as a result of species delimitation on DS2. The warning category (as in Table 1), the number of delimited groups and the geographic overall provenance of DNA barcode sequences included in DS2 for each species group are also reported.

| Suborder | Involved morphospecies | warning category | n of delimited groups | Geographic scale | Also occurring in DS1? |
| --- | --- | --- | --- | --- | --- |
| Zygoptera | *Coenagrion mercuriale* | intraspecific delimitation | 3 | Paleartic | no |
| Zygoptera | *Coenagrion puella* / *Coenagrion pulchellum* / *Coenagrion ornatum* | mixed species delimitation | 2 | Paleartic | no |
| Zygoptera | *Erythromma lindenii* | intraspecific delimitation | 2 | Paleartic | yes |
| Zygoptera | *Ischnura genei* / *Ischnura elegans* | no interspecific delimitation | 1 | Paleartic | yes |
| Zygoptera | *Calopteryx splendens* / *Calopteryx xanthostoma* | no interspecific delimitation | 1 | Paleartic | yes |
| Zygoptera | *Lestes dryas* | intraspecific delimitation | 2 | Holarctic | no |
| Zygoptera | *Chalcolestes viridis* / *Chalcolestes parvidens* | mixed species delimitation | 2 | Paleartic | yes |
| Anisoptera | *Aeshna juncea* | intraspecific delimitation | 2 | Holarctic | no |
| Anisoptera | *Anax imperator* / *Anax parthenope* | no interspecific delimitation | 1 | Paleartic | yes |
| Anisoptera | *Onychogomphus forcipatus* | intraspecific delimitation | 2 | Paleartic | yes |
| Anisoptera | *Stylurus flavipes* | intraspecific delimitation | 2 | Paleartic | no |
| Anisoptera | *Orthetrum albistylum* | intraspecific delimitation | 3 | Paleartic | no |
| Anisoptera | *Sympetrum danae* | intraspecific delimitation | 2 | Holarctic | no |
| Anisoptera | *Somatochlora metallica* / *Somatochlora meridionalis* | no interspecific delimitation | 1 | Paleartic | yes |

In Zygoptera, several changes occurred in species delimitation between DS1 and DS2 involving all the three categories of warnings (Table 1, Table 2 and Appendix S6). Warnings related to a reduced interspecific genetic divergence slightly decreased because *Coenagrion puella*, which in DS1 was delimited in a single group with *C. pulchellum* and *C. ornatum*, in DS2 showed a mixed delimitation pattern due to the inclusion of some North African highly divergent DNA barcode sequences, already reported by a previous study (Ferreira et al., 2016). The remaining two cases belonging to this category involved the species pairs *Ischnura elegans* (Vander Linden, 1820) */ I. genei* (Rambur, 1842) and *Calopteryx splendens* (Harris, 1780) */ C. xanthostoma* (Charpentier, 1825). Conversely, the warning cases (and species) of intraspecific delimitation increased with *Coenagrion mercuriale* and *Lestes dryas* Kirby, 1890 showing three and two lineages respectively, whereas the same two lineages of *Erythromma lindenii* found in DS1 were confirmed. Finally, apart from the already cited case of *C. puella* and close congenerics, *Chalcolestes viridis* and *C. parvidens* showed a mixed delimitation pattern, similarly to DS1. In Anisoptera, the groups of species not delimited remained the same of DS1 (i.e., *Anax imperator* Leach, 1815 / *A. parthenope* (Sélys, 1839) and *Somatochlora metallica* (Vander Linden, 1825) / *S. meridionalis*), whereas the number of warnings due to high intraspecific delimitation increased from two (one species) to 11 (five species). Concerning these latter, two lineages were delimited in *Aeshna juncea* (Linnaeus, 1758), *Onychogomphus forcipatus, Stylurus flavipes* (Charpentier, 1825) and *Sympetrum danae* (Sulzer, 1776), while three groups were delimited from *Orthetrum albistylum* (Sélys, 1848) DNA barcode sequences.

Finally, the “BIN Discordance” analysis conducted on BOLD was used to verify the warning cases mentioned above. Based on a similarity clustering strategy, COI barcode sequences are grouped into clusters which are assigned to a unique identifier called BIN (Ratnasingham & Hebert, 2013). This tool confirmed almost all the warning cases revealed by our multi-approach species delimitation. In particular, of the 88 barcoded Italian odonate morphospecies, 68 were assigned to unique BINs, 13 shared the same BINs and 7 were assigned to two or more BINs. The only case that did not match with our species delimitation results was *O. forcipatus*, for which all the sequenced specimens clustered with the public deposited DNA barcodes into a single BIN (see Appendix S1 for BINs assignment).

### Warnings investigation

Some warning cases, resulting from the species delimitation of DS2, involved odonate species represented by a high number of publicly available DNA barcode sequences from different geographic regions or by the availability of multi-locus data deposited in GenBank. These data allowed to better investigate the geographic distribution of genetic diversity at the COI barcode region for at least four Italian species, also distributed at the Palearctic or Holarctic scale (two Zygoptera, *Coenagrion mercuriale* and *Lestes dryas* and two Anisoptera, *Aeshna juncea* and *Sympetrum danae*, see Figure 4). The three groups of *C. mercuriale* DNA barcodes delimited in DS2 correspond precisely to three distinct haplogroups that are exclusive to Europe (except for Italy), North Africa and Italy, showing no shared haplotypes and high values of K2P and uncorrected *p*-distance among them (Figure 4, Appendix S7). Such isolation and genetic distinctiveness are also supported by uncorrected genetic *p*-distance values at another mitochondrial (16s rRNA) and three nuclear (AgT, PRMT, MLC) loci sequences retrieved in GenBank (see Appendix S7).

**FIGURE 4:**
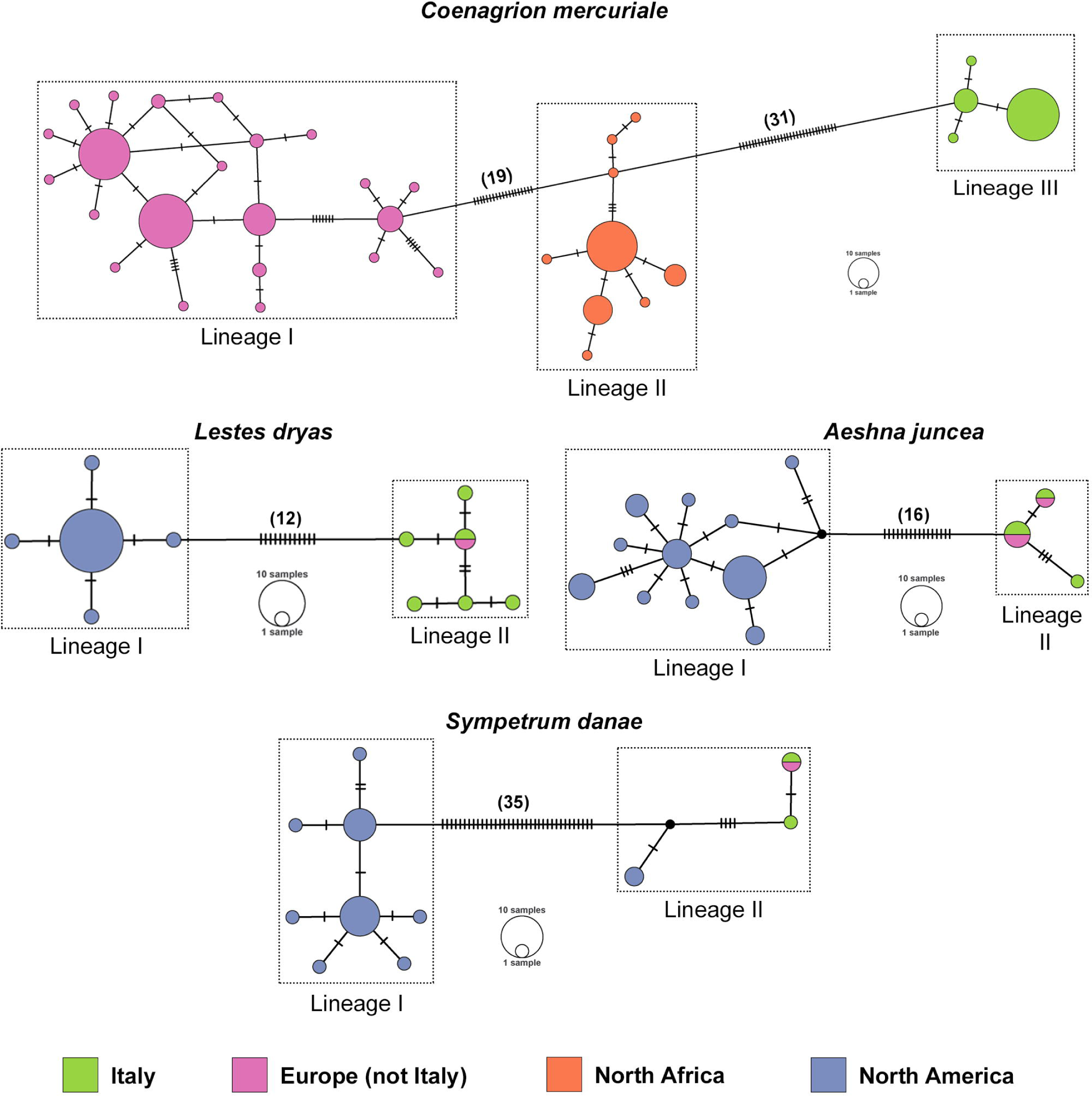
Median-joining network of COI DNA barcode haplotypes of four warning cases found by species delimitation of DS2 (*Coenagrion mercuriale, Lestes dryas, Aeshna juncea* and *Sympetrum danae*; see Appendix S2 and S6 for haplotypes and lineages subdivision). Each circle represents a haplotype and circle size is proportional to haplotype frequency. Colours indicate different sampling regions. Small black traits represent possible median vectors, while dashes represent substitutions (also indicated within brackets when > 10). Networks have been created with PopArt 1.7.

In the other three species depicted in Figure 4, the European (and Italian) haplotypes were clearly separated from the North American ones, supporting the delimitation inferred from DS2. The number of substitutions characterizing this Holarctic disjunction was more than the double in *S. danae* with respect to *A. juncea* and *L. dryas* delimited groups, with the only exception of one *S. danae* haplotype from Canada (A388, see Appendix S2) that is much closer to the Western Palearctic haplogroup.

Additional nuclear DNA sequences retrieved in GenBank allowed to better investigate the case of absent interspecific delimitation among *Coenagrion puella, C. pulchellum* and *C. ornatum*. However, these species were found to be clearly distinct at the three nuclear loci with no shared haplotypes, thus indicating a mito-nuclear discordance pattern of genetic divergence (Appendix S7).

## Discussion

### DNA barcoding performance and reference dataset

To the best of our knowledge, this study is one of the most comprehensive DNA barcoding-based surveys on Odonata conducted on a national scale in the Northern Hemisphere. Eighty-eight of the 89 Italian odonate species were successfully characterized by DNA barcoding, thus increasing the available reference molecular data for many taxa distributed at the European, Palearctic, or Holarctic scales (Figure 4, Table 2). *Ischnura fountaineae* Morton, 1905, was the only species which has not been collected, as just one isolated population is known to occur on Pantelleria island (Corso et al., 2012).

The multi-approach species delimitation showed a moderately high congruence between molecular groups and the Linnaean taxa with COI DNA barcodes, delimiting 85% of species in the reference Italian database as separate MOTUs (71% and 93% in Zygoptera and Anisoptera, respectively). Although most of the morphospecies are diagnosable, the identification performance of this study is lower compared to other odonate DNA barcoding surveys, which have reported up to a 95% success rate (Bergmann et al., 2013). However, all the previous DNA barcoding studies regarding odonates were based on a lower number of specimens and morphospecies (e.g., Bergmann et al., 2013; Kim et al., 2014; Koroiva et al., 2017). Concerning sequence variation within and among morphospecies, the average values obtained in this study are in line with those obtained by Koroiva and Kvist (2018) in their global overview of DNA barcoding of odonates, with the only exception of the mean intraspecific genetic variation in Italian Anisoptera (i.e., 0.33 DS1, 0.59 DS2), which is 4.8 (DS1) - 2.7 (DS2) times lower than the global average (i.e., 1.60). No more than a slight variation was observed in the average genetic distance values between DS1 and DS2. This was likely due to the higher completeness of DS2, where intraspecific variation was the result of the inclusion of samples from the whole Palearctic or Holarctic regions (Bergsten et al., 2012).

In the remaining cases (Table 1 and Table 2), the multi-approach species delimitation lumped the morphospecies or split them into additional molecular groups.

### Species delimitation and warnings

In this study, the multi-approach species delimitation conducted on DS1 showed that 32.3% of Zygoptera and 8.8% of Anisoptera species were highlighted by warnings (Table 1), and these percentages increased in DS2 (38.7% and 15.8%). While the discrepancy between the two datasets was due to the wider sampling area considered by the DS2 (see also Bergsten et al., 2012), the higher warning rate in Zygoptera is more challenging to explain. In both datasets, damselflies were involved in all the three possible groups of warnings, that even occur simultaneously within the genus *Coenagrion* (Table 1, Figure 2, Appendix S6). The situation of Italian Zygoptera could confirm previous findings that recorded a wide variety of global biogeographical patterns among damselflies (e.g., Sánchez-Guillén et al., 2014a; Swaegers et al., 2015; Ferreira et al., 2016; Wellenreuther & Sánchez-Guillén, 2016). Such a situation could be partly due to the weaker dispersal ability of damselflies as compared to dragonflies (Heiser & Schmitt 2010; Pinkert et al., 2018), which could have been resulted in more frequent isolation and diversification events. However, other studies (reviewed in Wellenreuther & Sánchez-Guillén 2016) point to sexual selection, and not ecological constraints, in playing the most important role in damselflies diversification over time and space.

Warnings identified by species delimitation of DS1 and DS2 provide evidence for different scenarios regarding Italian odonates taxonomy and conservation, with possible influence also at the Holarctic scale.

#### No interspecific delimitation

This category of warning encompasses four cases in DS2 that were already confirmed in the reference Italian sampling of DS1. The missing mitochondrial diversification in the damselflies species pairs *Ischnura elegans/I. genei* and *Calopteryx splendens/C. xanthostoma*, even though poorly investigated so far for Italian populations, was expected. The reproductive isolation of both species pairs in Mediterranean areas is indeed highly debated by entomologists with morphological and genetic evidence of introgressive hybridization, largely documented in *Ischnura* (Sánchez-Guillén et al., 2014a; 2014b) and less clear in *Calopteryx* (Dumont, Mertens, & De Coster, 1993; Weekers, De Jonckheere, & Dumont, 2001). Conversely, the missing delimitation in *Somatochlora metallica/S. meridionalis* and *Anax imperator*/*A. parthenope* was unprecedented, also because no Italian samples and no COI barcode sequences at all for *S. meridionalis* were available before this study. This taxon was originally described as a subspecies of *S. metallica* (Nielsen, 1935) and elevated at species rank more recently (Schmidt, 1957); however, this status is still debated as intermediate specimens have been recorded in the contact areas (Fleck, Grand, & Boudot, 2007). *S. meridionalis* replaces *S. metallica* in southern Europe; in Italy, both the morphospecies occur with the former distributed in central and southern portions of the peninsula (and in two disjunct and isolated populations in the North) and the latter mainly spread in northern regions. Both taxa are sometimes found syntopically (Boudot & Kalkman, 2015; Dijkstra & Kalkman, 2012) and lack clear structural differences apart from subtle, though stable, differences in colouration in the adults (Boudot & Kalkman, 2015) and minor differences at the larval stage (Seidenbusch, 1996). Species delimitation based on COI barcode sequences from specimens coming from all the three disjunct Italian ranges of *S. meridionalis* suggests that the two morphospecies in Italy are not distinguishable according to mitochondrial DNA. More challenging is the case of the two *Anax* species, which, contrary to *Somatochlora*, are well separated morphologically (but also according to their ecology and life-history traits), although they may share the same mitochondrial haplotypes. This scenario is not exclusive to Italian populations, as the same pattern was confirmed by DS2 European and Palearctic populations (see also Rewicz et al., 2020).

#### Intraspecific delimitation

The first group of morphospecies showing marked intraspecific genetic divergence encompasses the Italian samples (DS1) of *Erythromma lindenii* and *Onychogomphus forcipatus*. Concerning the former taxon, it was initially described as *Agrion lindenii*, placed in the new genus *Cercion* by Navas (1907) and recently assigned to its current genus by Weekers & Dumont (2004). Across its vast range, it shows remarkably little morphological variation, but the eastern populations were described as *E. l. zernyi* (Schmidt, 1938). This taxon occurs from Turkey eastward in isolated populations, surrounded by, and/or overlapping with the nominal subspecies; thus, a longitudinal introgression generating hybrid populations was hypothesized, although no genetic data has ever confirmed this (Dumont et al., 1995). Unexpectedly, our Italian sampling, and the comparison with the few available European COI barcode sequences, provide evidence for the occurrence of two supported mitochondrial lineages within western Palearctic (Figure 2, Appendix S6). One of these lineages is completely new, and, based on these preliminary data, it is mainly occurring in Montenegro and north-eastern Italy sampling localities.

*Onychogomphus forcipatus* has three recognized subspecies showing structural differentiation (both in adults and nymphs) and occurring in Europe, with discrete and mostly non-overlapping ranges. Of these, two subspecies are recorded in Italy, *O. f. forcipatus* being distributed only in the northeast and allegedly in Sicily, and *O. f. unguiculatus* (Vander Linden, 1823) being distributed in north-western and peninsular Italy (Boudot, Jacquemin, & Dumont, 1990, Boudot & Kalkman, 2015). Our species delimitation approach identifies two distinct lineages as well (also supported in DS2); however, these do not show clear geographic separation, with a group including Sicilian, central, southern, and north-western Italy samples, but also several samples from north-eastern Italy and the other one encompassing a few samples from north-eastern Italy where the other lineage also occurs. In this sense, our study, as that by Ferreira et al. (2014b), is still not resolutive, and further integrated studies are needed to possibly solve this long-lasting taxonomic enigma.

Other warning cases of intraspecific delimitation emerged from the analysis of DS2 and encompass lineages diverging at the Palearctic scale. The most interesting case is *Coenagrion mercuriale* that forms three delimited haplogroups, of which one exclusive of the Italian peninsula. In 1948, *C. castellani* was described from central Italy (Roberts, 1948), and in 1949 it was reported that *C. mercuriale* male specimens from Italy invariably showed clear morphological differences (i.e., male appendages) with respect to specimens from central Europe, suggesting to treat the Italian ones as *C. m. castellani* (Conci, 1949). Notably, the Italian populations show a geographic disjunction from the range of *C. mercuriale*, as also supported by genetic distance values at other mitochondrial and nuclear loci (Figure 4, Appendix S7). Similarly, the populations from the Maghreb were referred to as *C. m. hermeticum* (Selys 1872), since they appeared slightly morphologically distinct from the nominal subspecies (Ben Azzouz, Guemmouh, & Aguesse, 1989). The validity of *hermeticum* was questioned by several authors (e.g. Lieftinck, 1966), although the necessity of further comparative investigations was advocated (Jacquemin & Boudot 1990; 1999). This long-awaited study did not come to light until a recent multi-locus characterization of North African populations (Ferreira et al., 2014a) and the data here provided for *castellani*. Although Boudot & Kalkman (2015) stated that at present no subspecies of *C. mercuriale* are recognized, our results suggest that three clearly distinct lineages occur within its range, one of which possibly endemic to Italy. Further research including morphometric investigations is definitely needed to clarify the matter, also considering the high conservation measures involving *C. mercuriale* in Europe, which is listed in the Annex II of Council Directive 92/43/EEC.

Other two intraspecific delimitation warnings emerging from DS2 concerned *Orthetrum albistylum* and *Stylurus flavipes*. These showed, respectively, two and one delimited Asian lineages, in addition to the Italian/European one. Both taxa have a huge Eurasian range and while some (maybe isolated) subspecies have been described for *O. albistylum* (e.g., *O. a. speciosum* (Uhler, 1858), see Schorr & Paulson, 2019), *S. flavipes* has no currently accepted subspecies (Steinmann, 2013; Schorr & Paulson, 2019). Unfortunately, only a few DNA barcode sequences are available for these taxa and this does not permit further clarification of the observed warnings.

The last group of intraspecific delimitation warnings includes taxa having a Holarctic distribution, namely *Lestes dryas* (Zygoptera), *Aeshna juncea*, and *Sympetrum danae* (Anisoptera). The COI DS2 haplotypes of these three species showed a marked geographic structure with Italian/European haplotypes diverging from the North American ones (Figure 4, Appendix S6). Regarding *A. juncea*, Bartenev in 1929 described the subspecies *A. j. americana* based on material from Canada (Steinmann, 2013). Subsequently, Boudot & Kalkman (2015) stated that none of the subspecies described should be considered valid while admitting that the specimens from North America are clearly morphologically different from those found in Europe. *Lestes dryas* and *S. danae* were described based on European specimens and no intraspecific taxa are currently recognized (Bridges 1994; Steinmann 2013), making the warnings here presented and their possible taxonomic significance an element of primary importance to be clarified.

#### Mixed species delimitation

This warning category involves two morphospecies groups of damselflies which DNA barcoded individuals showed either marked genetic divergence or complete sequence identity with congenerics. The first case, also occurring in DS1, concerns *Chalcolestes viridis* and *C. parvidens*. The latter was originally described as a subspecies of *C. viridis* and was subsequently elevated to the species rank (Dell’Anna, Utzeri, & De Matthaeis, 1996; Gyulavari et al., 2011). Both species can occur syntopically in the Italian peninsula and in other parts of Europe, with hybridization events reported (Olias et al., 2007). Our DNA barcoding data confirm the genetic differentiation between *C. viridis* and *C. parvidens*, even if one specimen from Apulia, morphologically identified as *C. viridis*, was genetically assigned to *C. parvidens*. Several *Chalcolestes* populations from southern Italy are often difficult to be morphologically identified (authors pers. obs.), and intermediate morphological traits could be due to the close relationship and relatively recent divergence between the two taxa (Gyulavari et al., 2011). Hybridization and/or incomplete lineage sorting events could have played a role in causing this warning. A multi-locus phyogeographic survey, based on an extensive sampling, especially in the contact zones between the two taxa, is needed to clarify the matter. Surprisingly, a mixed species delimitation warning was found among *Coenagrion puella, C. pulchellum* and *C. ornatum* in DS2, with *C. puella* showing a delimited diverging lineage corresponding to North African populations and the other Italian/western Palearctic representatives of the three species delimited in a single group and even sharing the same haplotypes (as already found in DS1). These taxa can be found in syntopic conditions but can be clearly distinguished through morphology, ecological requirements, life-history traits, though sporadic hybridization is known to occur among them. The marked divergence at the investigated nuclear markers (Appendix S7) indicates that a mito-nuclear discordance pattern occurs throughout their overall distribution with the only exception of northern African populations, as already described by Ferreira et al. (2016). Also in this case, further investigation is needed to better address such discordance, also considering the high conservation value of *C. ornatum*, which is included in the Annex II of Council Directive 92/43/EEC.

On the whole, our comprehensive DNA barcoding characterization of Italian Odonata sheds light on several novel and other long-time debated taxonomic aspects of this taxon, with an unexpected geographical resonance well beyond Italy. The newly detected lineages may lead to the discovery of separate cryptic taxonomic entities within the Linnaean species and may provide fertile ground for future studies. Conversely, the reduced or missing delimitation between some known species challenges the current taxonomic knowledge on this presumed “well-explored” group of insects (at least in Europe). The warnings resulting from our species delimitation approach could indicate Italy as an important biogeographical and evolutionary centre for odonates diversification. A wide array of possible scenarios could explain the warnings found by our DNA barcoding-based species delimitation, such as the occurrence of still undiscovered cryptic species (e.g., the unexpected and most surprising recent description of a novel species endemic to Spain, *Onychogomphus cazuma*; López-Estrada et al., 2020) and different forms of mito-nuclear discordance due to more or less ancient introgression phenomena (e.g., Solano et al., 2018; Ottenburghs, 2020).

Further sampling campaigns, in Italy and throughout the whole range of the warning species groups (as recommended by Gaytán et al., 2020), the integration of multiple genetic markers (e.g., the universal single-copy orthologs USCOs recently described by Eberle et al., 2020), and accurate morphometrics analyses will be necessary to correctly address each possible case of taxonomic (and conservation) interest.

Finally, this thorough DNA barcoding investigation of Italian Odonata and related warnings will allow to better support and interpret HTS eDNA metabarcoding data from European and even Holarctic freshwater environments and, at the same time, to design species-specific (or lineage-specific) probes to reliably address conservation studies on the whole range of the investigated species.

## Acknowledgments

Stefano Aguzzi, Gianluca Ancarani, Paolo Biella, Giovanni Boano, Marco Bonifacino, Martina Cadin, Gianmaria Carchini, Giovanni Colombo, Andrea Cusmano, Davide De Rosa, Federica Ferrario, Tiziano Fiorenza, Ilaria Fozzi, Carlo Galliani, Roberto Garavaglia, Gabriele Gheza, Luca Giussani, Marco Guglielmi, Cristiano Liuzzi, Davide Magnani, Alexandro Minicò, Valerio Orioli, Francesco Ornaghi, Silvia Pelti, Prisca, Verena Penna, Pietro Ramellini, Gloria Ramello, Francesca Ricci, Laura Ricci, Giuseppe Rossi, Leonardo Siddi, Elena Ternelli, Nicola Tommasi, Parco Adda Nord, Parco Lombardo della valle del Ticino, Parco della Valle Lambro, Parco Regionale Marturanum. We thank the association Odonata.it for warmly welcoming this project. The “Servizio Sviluppo Sostenibile e Aree Protette of Provincia Autonoma di Trento” allowed specimen collection in full-protected areas in Trentino (Prot. S175/2018/382452/17.11.2/LS/57EI); and the Italian Ministry of the Environment, Land and Sea released a national permit for the collection of species included in European and Italian conservation directives (Prot. 0031783.20-11-2019).

## Data Accessibility

All specimen data are accessible on BOLD (www.boldsystems.org) under the project name ‘ZPLOD - DNA barcoding Italian Odonata’. The data include collection locality, geographic coordinates, collector, identifier, voucher depository, BINs and other sampling details. Sequence data are available on BOLD and have also been deposited in GenBank (Accession numbers MT298232 - MT298679).

## Author contributions

AG and GA conceived the idea and designed the sampling protocol, which was then developed by all authors. AG, GA, DB, GB, IC, AC, VF, MG, LI, GLP, LL, FL, FM, SR, RS, GS, SS conducted the field sampling. FR with the help of AG stored the biological tissues, performed the lab analyses, and mined data from BOLD and GenBank. AG, GA, and DM ran the statistical analyses and all authors contributed to analyses checking/development and check the final model outcomes. AG, GA and DM lead the writing of a first draft of the manuscript and all authors critically contributed revising the final version.

## Supporting information

**APPENDIX S1:** dataset DS1, composed by the DNA barcoding sequences produced in this study on Italian samples (409) plus 69 GenBank sequences from other Italian specimens. Species taxonomy, sex, specimen IDs, depositories accessions, sampling geographic details and COI haplotype are reported for each DNA barcoded sample.

**APPENDIX S2:** dataset DS2, composed by the sequences mined from BOLD and GenBank plus dataset DS1 and other COI barcode sequences obtained from additional samples retrieved by authors in other western Palearctic countries. Species taxonomy, DS1 specimen IDs, depositories accessions, are reported for each COI haplotype.

**APPENDIX S3:** Support values of the species delimitation hypotheses obtained with the Bayesian implementation of the PTP and GMYC methods, for both DS1 and DS2 (n.d.: not detected in DS1).

**APPENDIX S4:** Distributions of pairwise genetic distances (K2P) obtained with ABGD for a) Zygoptera DS1, b) Anisoptera DS1, c) Zygoptera DS2, and d) Anisoptera DS2.

**APPENDIX S5:** Barcode Gap Analysis of DS1 for Anisoptera and Zygoptera respectively, generated by BOLD. Three scatterplots are provided to confirm the existence and magnitude of the Barcode Gap. For each suborder, the first two scatterplots show the overlap of the max and mean intra-specific distances vs the inter-specific (nearest neighbour) distances. The third scatterplot plots the number of individuals in each species against their max intra-specific distances, as a test for sampling bias. Distance Model: Kimura 2 Parameter, Deletion Method: Pairwise Deletion, Alignment: BOLD Aligner (Amino Acid based HMM), Filters Applied: ≥ 500 bp only.

**APPENDIX S6:** Multi-approach species delimitation of the 31 Zygoptera and 57 Anisoptera species investigated in this study based on Holarctic COI DNA barcode sequences from DS2. A Bayesian tree is used as a base to summarize the two threshold-based (OT and ABGD) and four character explicit-based (PTP, MPTP, GMYC, bGMYC) approaches. Specimen haplotype identifiers are reported on tips (see Appendix S1 and S2 for further details) and the number of specimens sharing each haplotype is reported within brackets. Numbers above nodes represent BPP and BS, respectively, whereas asterisks correspond to maximal node support (BPP ≥ 0.99 and BS ≥ 95). Vertical colored solid boxes delimit putative species identified by the different approaches. Black: delimitation congruent with the identified morphospecies; Red: intraspecific delimitation; Light Blue: no interspecific delimitation; Yellow: Mixed species delimitation. In case of intraspecific delimitation, each lineage has been numbered with Roman numerals. This tree was created with BEAST 1.8.2.

**APPENDIX S7:** average genetic p-distance divergence and Standard Deviation values among the geographic population of *Coenagrion mercuriale* (ITA: Italy, EUR: Europe excluding Italy, NAF: North Africa) and among three species belonging to *Coenagrion* (ORN: *ornatum*, PUL: *pulchellum*, PUE: *puella*) calculated at two mitochondrial (16s rDNA and COI) and three nuclear (AgT, PRMT and MLC) markers. Sequence data were retrieved from GenBank. Concerning COI, also the sequences produced in this study were used to calculate the genetic distances. The exclusiveness of haplotypes belonging to the Italian populations of *C. mercurial* and to the other three *Coenagrion* species is indicated for each genetic marker. Genetic distance values have been calculated using MEGA X (Kumar et al., 2018).

