## Appendix S1-S5,S7 for "Italian Odonates in the Pandora’s Box: A Comprehensive DNA Barcoding Inventory Shows Taxonomic Warnings at the Holarctic Scale"

MOLECULAR ECOLOGY RESOURCES

#### Table of Contents:



| | |
|---|---|
| **Appendix S1** | Page 2 |
| **Appendix S2** | Page 11 |
| **Appendix S3** | Page 21 |
| **Appendix S4** | Page 24 |
| **Appendix S5** | Page 25 |
| **Appendix S7** | Page 27 |

MOLECULAR ECOLOGY
RESOURCES

**APPENDIX S1:** dataset DS1, composed by the DNA barcoding sequences produced in this study on Italian samples (409) plus 69 NCBI Genbank sequences from other Italian specimens. Species taxonomy, sex, specimen IDs, Barcode Index Number (BIN), depositories accessions, sampling geographic details and COI haplotype are reported for each DNA barcoded sample.

| BOLD PROCESS ID | BIN | VOUCHER ID | GENBANK a.n. | Suborder | Family | SPECIES | SEX | ADMINISTRATIVE REGION | ADMINISTRATIVE PROVINCE | Coordinate N, WGS84 | Coordinate E, WGS84 | MACRO-REGION | HAPLOTYPE | SOURCE |
|---|---|---|---|---|---|---|---|---|---|---|---|---|---|---|
| ZPLOD001-20 | BOLD:AAJ5779 | MIB:ZPL:07801 | MT298234 | Anisoptera | Aeshnidae | *Aeshna affinis* | M | Sardinia | Sassari | 40.9 | 8.2 | SA | A1 | this study |
| ZPLOD004-20 | BOLD:AAJ5779 | MIB:ZPL:07804 | MT298233 | Anisoptera | Aeshnidae | *Aeshna affinis* | M | Piedmont | Torino | 45.1 | 7.5 | NW | A2 | this study |
| ZPLOD007-20 | BOLD:AAJ5779 | MIB:ZPL:07807 | MT298232 | Anisoptera | Aeshnidae | *Aeshna affinis* | M | Apulia | Taranto | 40.6 | 16.8 | S | A3 | this study |
| ZPLOD010-20 | BOLD:AAA6531 | MIB:ZPL:07810 | MT298235 | Anisoptera | Aeshnidae | *Aeshna caerulea* | M | Trentino-Alto Adige | Trento | 46.4 | 11.8 | NE | A4 | this study |
| _ | _ | _ | KU180305 | Anisoptera | Aeshnidae | *Aeshna cyanea* | _ | Sicily | _ | _ | _ | SI | A5 | NCBI GenBank |
| ZPLOD016-20 | BOLD:ACI1053 | MIB:ZPL:07816 | MT298236 | Anisoptera | Aeshnidae | *Aeshna cyanea* | M | Lombardy | Bergamo | 45.9 | 9.8 | NW | A6 | this study |
| ZPLOD017-20 | BOLD:AAJ5811 | MIB:ZPL:07817 | MT298238 | Anisoptera | Aeshnidae | *Aeshna grandis* | M | Trentino-Alto Adige | Trento | 46.1 | 11.2 | NE | A7 | this study |
| ZPLOD018-20 | BOLD:AAJ5811 | MIB:ZPL:07818 | MT298237 | Anisoptera | Aeshnidae | *Aeshna grandis* | M | Aosta Valley | Aosta | 45.8 | 7.6 | NW | A8 | this study |
| ZPLOD019-20 | BOLD:ADC2941 | MIB:ZPL:07819 | MT298241 | Anisoptera | Aeshnidae | *Aeshna isoceles* | M | Sardinia | Sassari | 40.9 | 8.2 | SA | A9 | this study |
| ZPLOD023-20 | BOLD:ADC2941 | MIB:ZPL:07823 | MT298240 | Anisoptera | Aeshnidae | *Aeshna isoceles* | M | Apulia | Brindisi | 40.7 | 17.8 | S | A10 | this study |
| ZPLOD024-20 | BOLD:ADC2941 | MIB:ZPL:07824 | MT298239 | Anisoptera | Aeshnidae | *Aeshna isoceles* | F | Lombardy | Como | 45.8 | 9.3 | NW | A11 | this study |
| ZPLOD028-20 | BOLD:AAJ1281 | MIB:ZPL:07828 | MT298243 | Anisoptera | Aeshnidae | *Aeshna juncea* | F | Aosta Valley | Aosta | 45.8 | 7.6 | NW | A12 | this study |
| ZPLOD030-20 | BOLD:AAJ1281 | MIB:ZPL:07830 | MT298244 | Anisoptera | Aeshnidae | *Aeshna juncea* | M | Trentino-Alto Adige | Trento | 46.3 | 11.3 | NE | A13 | this study |
| ZPLOD033-20 | BOLD:AAJ1281 | MIB:ZPL:07833 | MT298245 | Anisoptera | Aeshnidae | *Aeshna juncea* | M | Lombardy | Sondrio | 46.4 | 9.4 | NW | A14 | this study |
| ZPLOD034-20 | BOLD:AAJ5810 | MIB:ZPL:07834 | MT298247 | Anisoptera | Aeshnidae | *Aeshna mixta* | M | Marches | Macerata | 43.3 | 13.4 | C | A15 | this study |
| ZPLOD038-20 | BOLD:AAJ5810 | MIB:ZPL:07838 | MT298248 | Anisoptera | Aeshnidae | *Aeshna mixta* | F | Lombardy | Como | 45.8 | 9.3 | NW | A16 | this study |
| ZPLOD040-20 | BOLD:ABZ5296 | MIB:ZPL:07840 | MT298249 | Anisoptera | Aeshnidae | *Aeshna subarctica* | M | Trentino-Alto Adige | Trento | 46.3 | 11.3 | NE | A17 | this study |
| ZPLOD041-20 | BOLD:ABZ5296 | MIB:ZPL:07841 | MT298250 | Anisoptera | Aeshnidae | *Aeshna subarctica* | M | Trentino-Alto Adige | Trento | 46.2 | 11.4 | NE | A18 | this study |
| ZPLOD042-20 | BOLD:ACH7840 | MIB:ZPL:07842 | MT298251 | Anisoptera | Aeshnidae | *Anax ephippiger* | M | Liguria | Genova | 44.4 | 8.8 | NW | A19 | this study |
| ZPLOD044-20 | BOLD:ACH7840 | MIB:ZPL:07844 | MT298252 | Anisoptera | Aeshnidae | *Anax ephippiger* | F | Lombardy | Pavia | 45.2 | 8.7 | NW | A20 | this study |
| ZPLOD045-20 | BOLD:ABX6596 | MIB:ZPL:07845 | MT298260 | Anisoptera | Aeshnidae | *Anax imperator* | M | Sardinia | Sassari | 40.7 | 8.2 | SA | A21 | this study |
| ZPLOD046-20 | BOLD:ABX6596 | MIB:ZPL:07846 | MT298257 | Anisoptera | Aeshnidae | *Anax imperator* | M | Marches | Fermo | 43.2 | 13.6 | C | A21 | this study |
| ZPLOD048-20 | BOLD:ABX6596 | MIB:ZPL:07848 | MT298258 | Anisoptera | Aeshnidae | *Anax imperator* | F | Trentino-Alto Adige | Trento | 46.2 | 11.3 | NE | A22 | this study |
| ZPLOD049-20 | BOLD:ABX6596 | MIB:ZPL:07849 | MT298259 | Anisoptera | Aeshnidae | *Anax imperator* | F | Piedmont | Torino | 45.1 | 7.2 | NW | A21 | this study |
| ZPLOD051-20 | BOLD:ABX6596 | MIB:ZPL:07851 | MT298253 | Anisoptera | Aeshnidae | *Anax imperator* | M | Sicily | Messina | 38 | 15.3 | SI | A23 | this study |
| ZPLOD053-20 | BOLD:ABX6596 | MIB:ZPL:07853 | MT298256 | Anisoptera | Aeshnidae | *Anax imperator* | M | Lombardy | Milano | 45.3 | 9 | NW | A21 | this study |
| ZPLOD056-20 | BOLD:ABX6596 | MIB:ZPL:07856 | MT298255 | Anisoptera | Aeshnidae | *Anax imperator* | M | Molise | Campobasso | 41.6 | 14.8 | S | A24 | this study |
| ZPLOD057-20 | BOLD:ABX6596 | MIB:ZPL:07857 | MT298254 | Anisoptera | Aeshnidae | *Anax imperator* | M | Sicily | Trapani | 37.9 | 12.7 | SI | A25 | this study |
| ZPLOD058-20 | BOLD:ABX6596 | MIB:ZPL:07858 | MT298261 | Anisoptera | Aeshnidae | *Anax parthenope* | M | Marches | Macerata | 43.3 | 13.5 | C | A25 | this study |
| ZPLOD060-20 | BOLD:ABX6596 | MIB:ZPL:07860 | MT298263 | Anisoptera | Aeshnidae | *Anax parthenope* | _ | Sicily | Catania | 37.5 | 14.9 | SI | A27 | this study |
| ZPLOD062-20 | BOLD:ABX6596 | MIB:ZPL:07862 | MT298264 | Anisoptera | Aeshnidae | *Anax parthenope* | M | Lombardy | Varese | 45.7 | 8.7 | NW | A27 | this study |
| ZPLOD063-20 | BOLD:ABX6596 | MIB:ZPL:07863 | MT298262 | Anisoptera | Aeshnidae | *Anax parthenope* | M | Molise | Campobasso | 42 | 14.8 | S | A28 | this study |
| ZPLOD066-20 | BOLD:ADN2163 | MIB:ZPL:07866 | MT298266 | Anisoptera | Aeshnidae | *Boyeria irene* | M | Sardinia | Sud Sardegna | 39.4 | 8.7 | SA | A29 | this study |
| ZPLOD068-20 | BOLD:ADN2163 | MIB:ZPL:07868 | MT298267 | Anisoptera | Aeshnidae | *Boyeria irene* | M | Piedmont | Torino | 45.1 | 7.5 | NW | A30 | this study |
| ZPLOD070-20 | BOLD:ADN2163 | MIB:ZPL:07870 | MT298265 | Anisoptera | Aeshnidae | *Boyeria irene* | M | Lombardy | Pavia | 45.3 | 9 | NW | A30 | this study |
| ZPLOD072-20 | BOLD:ABA9406 | MIB:ZPL:07872 | MT298269 | Anisoptera | Libellulidae | *Brachythemis impartita* | F | Sardinia | Sassari | 40.7 | 9 | SA | A31 | this study |
| ZPLOD073-20 | BOLD:ABA9406 | MIB:ZPL:07873 | MT298271 | Anisoptera | Libellulidae | *Brachythemis impartita* | M | Sardinia | Oristano | 39.9 | 8.5 | SA | A31 | this study |
| ZPLOD075-20 | BOLD:ABA9406 | MIB:ZPL:07875 | MT298270 | Anisoptera | Libellulidae | *Brachythemis impartita* | F | Sicily | Catania | 37.5 | 14.9 | SI | A31 | this study |
| ZPLOD078-20 | BOLD:ACI1765 | MIB:ZPL:07878 | MT298275 | Anisoptera | Aeshnidae | *Brachytron pratense* | M | Umbria | Terni | 42.5 | 12.8 | C | A33 | this study |
| ZPLOD079-20 | BOLD:ACI1765 | MIB:ZPL:07879 | MT298274 | Anisoptera | Aeshnidae | *Brachytron pratense* | M | Friuli-Venezia Giulia | Udine | 45.8 | 13.1 | NE | A34 | this study |
| ZPLOD081-20 | BOLD:ACI1765 | MIB:ZPL:07881 | MT298273 | Anisoptera | Aeshnidae | *Brachytron pratense* | M | Apulia | Lecce | 40.2 | 18.4 | S | A34 | this study |
| ZPLOD082-20 | BOLD:ACI1765 | MIB:ZPL:07882 | MT298272 | Anisoptera | Aeshnidae | *Brachytron pratense* | M | Piedmont | Novara | 45.4 | 8.8 | NW | A35 | this study |
| ZPLOD083-20 | BOLD:ADV2208 | MIB:ZPL:07883 | MT298277 | Zygoptera | Calopterygidae | *Calopteryx haemorrhoidalis* | M | Sardinia | Sassari | 40.5 | 8.6 | SA | Z1 | this study |
| ZPLOD084-20 | BOLD:ADV2208 | MIB:ZPL:07884 | MT298278 | Zygoptera | Calopterygidae | *Calopteryx haemorrhoidalis* | M | Liguria | Genova | 44.5 | 8.8 | NW | Z2 | this study |
| ZPLOD085-20 | BOLD:ADV2208 | MIB:ZPL:07885 | MT298279 | Zygoptera | Calopterygidae | *Calopteryx haemorrhoidalis* | F | Liguria | Genova | 44.5 | 8.8 | NW | Z3 | this study |
| ZPLOD086-20 | BOLD:ADV2208 | MIB:ZPL:07886 | MT298280 | Zygoptera | Calopterygidae | *Calopteryx haemorrhoidalis* | M | Liguria | Genova | 44.5 | 8.8 | NW | Z3 | this study |
| ZPLOD091-20 | BOLD:ADV2208 | MIB:ZPL:07891 | MT298281 | Zygoptera | Calopterygidae | *Calopteryx haemorrhoidalis* | M | Apulia | Barletta-Andria-Trani | 41 | 16.2 | S | Z1 | this study |
| ZPLOD095-20 | BOLD:ADV2208 | MIB:ZPL:07895 | MT298282 | Zygoptera | Calopterygidae | *Calopteryx haemorrhoidalis* | F | Sicily | Messina | 38 | 15.3 | SI | Z1 | this study |

MOLECULAR ECOLOGY
RESOURCES

| BOLD PROCESS ID | BIN | VOUCHER ID | GENBANK a.n. | Suborder | Family | SPECIES | SEX | ADMINISTRATIVE REGION | ADMINISTRATIVE PROVINCE | Coordinate N, WGS84 | Coordinate E, WGS84 | MACRO-REGION | HAPLOTYPE | SOURCE |
|---|---|---|---|---|---|---|---|---|---|---|---|---|---|---|
| ZPLOD096-20 | BOLD:ADV2208 | MIB:ZPL:07896 | MT298283 | Zygoptera | Calopterygidae | *Calopteryx haemorrhoidalis* | M | Tuscany | Grosseto | 43 | 10.9 | C | Z1 | this study |
| ZPLOD098-20 | BOLD:ADV2208 | MIB:ZPL:07898 | MT298284 | Zygoptera | Calopterygidae | *Calopteryx haemorrhoidalis* | M | Lazio | Latina | 41.4 | 13.2 | C | Z1 | this study |
| ZPLOD102-20 | BOLD:ADC4648 | MIB:ZPL:07902 | MT298288 | Zygoptera | Calopterygidae | *Calopteryx splendens* | M | Apulia | Bari | 40.8 | 16.4 | S | Z5 | this study |
| ZPLOD103-20 | BOLD:ADC4648 | MIB:ZPL:07903 | MT298289 | Zygoptera | Calopterygidae | *Calopteryx splendens* | M | Trentino-Alto Adige | Trento | 46 | 11.3 | NE | Z6 | this study |
| ZPLOD110-20 | BOLD:ADC4648 | MIB:ZPL:07910 | MT298287 | Zygoptera | Calopterygidae | *Calopteryx splendens* | M | Lombardy | Como | 45.8 | 9.2 | NW | Z6 | this study |
| ZPLOD113-20 | BOLD:ADC4648 | MIB:ZPL:07913 | MT298286 | Zygoptera | Calopterygidae | *Calopteryx splendens* | M | Lazio | Latina | 41.5 | 13.1 | C | Z8 | this study |
| ZPLOD114-20 | BOLD:AAE7398 | MIB:ZPL:07914 | MT298290 | Zygoptera | Calopterygidae | *Calopteryx virgo* | M | Piedmont | Cuneo | 44.3 | 7.7 | NW | Z9 | this study |
| ZPLOD115-20 | BOLD:AAE7398 | MIB:ZPL:07915 | MT298291 | Zygoptera | Calopterygidae | *Calopteryx virgo* | M | Marches | Macerata | 43.1 | 13.3 | C | Z10 | this study |
| ZPLOD117-20 | BOLD:AAE7398 | MIB:ZPL:07917 | MT298293 | Zygoptera | Calopterygidae | *Calopteryx virgo* | M | Trentino-Alto Adige | Trento | 46.1 | 11 | NE | Z9 | this study |
| ZPLOD119-20 | BOLD:AAE7398 | MIB:ZPL:07919 | MT298294 | Zygoptera | Calopterygidae | *Calopteryx virgo* | M | Umbria | Perugia | 42.8 | 13.1 | C | Z10 | this study |
| ZPLOD123-20 | BOLD:AAE7398 | MIB:ZPL:07923 | MT298295 | Zygoptera | Calopterygidae | *Calopteryx virgo* | M | Lombardy | Pavia | 45.3 | 9 | NW | Z9 | this study |
| ZPLOD124-20 | BOLD:ADC4648 | MIB:ZPL:07924 | MT298297 | Zygoptera | Calopterygidae | *Calopteryx xanthostoma* | M | Liguria | Savona | 44.1 | 8.2 | NW | Z6 | this study |
| ZPLOD125-20 | BOLD:ADC4648 | MIB:ZPL:07925 | MT298296 | Zygoptera | Calopterygidae | *Calopteryx xanthostoma* | F | Liguria | Savona | 44.1 | 8.2 | NW | Z6 | this study |
| ZPLOD127-20 | BOLD:ACH6070 | MIB:ZPL:07927 | MT298301 | Zygoptera | Coenagrionidae | *Ceriagrion tenellum* | M | Sardinia | Sassari | 40.4 | 8.6 | SA | Z14 | this study |
| ZPLOD129-20 | BOLD:ACH6070 | MIB:ZPL:07929 | MT298300 | Zygoptera | Coenagrionidae | *Ceriagrion tenellum* | M | Friuli-Venezia Giulia | Udine | 45.9 | 13.2 | NE | Z15 | this study |
| ZPLOD130-20 | BOLD:ACH6070 | MIB:ZPL:07930 | MT298299 | Zygoptera | Coenagrionidae | *Ceriagrion tenellum* | M | Sicily | Siracusa | 37.1 | 15.3 | SI | Z14 | this study |
| ZPLOD134-20 | BOLD:ACH6070 | MIB:ZPL:07934 | MT298303 | Zygoptera | Coenagrionidae | *Ceriagrion tenellum* | M | Apulia | Lecce | 40.2 | 18.4 | S | Z14 | this study |
| ZPLOD135-20 | BOLD:ACH6070 | MIB:ZPL:07935 | MT298302 | Zygoptera | Coenagrionidae | *Ceriagrion tenellum* | M | Lombardy | Como | 45.8 | 9.2 | NW | Z15 | this study |
| ZPLOD138-20 | BOLD:ACH6070 | MIB:ZPL:07938 | MT298298 | Zygoptera | Coenagrionidae | *Ceriagrion tenellum* | M | Piedmont | Cuneo | 44.5 | 7.7 | NW | Z15 | this study |
| ZPLOD140-20 | BOLD:ACH6070 | MIB:ZPL:07940 | MT298304 | Zygoptera | Coenagrionidae | *Ceriagrion tenellum* | M | Lazio | Latina | 41.5 | 13.1 | C | Z14 | this study |
| ZPLOD141-20 | BOLD:ADR7794 | MIB:ZPL:07941 | MT298308 | Zygoptera | Lestidae | *Chalcolestes parvidens* | M | Umbria | Terni | 42.6 | 12.2 | C | Z17 | this study |
| ZPLOD142-20 | BOLD:ADR7794 | MIB:ZPL:07942 | MT298310 | Zygoptera | Lestidae | *Chalcolestes parvidens* | F | Friuli-Venezia Giulia | Trieste | 45.6 | 13.8 | NE | Z17 | this study |
| ZPLOD143-20 | BOLD:ADR7794 | MIB:ZPL:07943 | MT298311 | Zygoptera | Lestidae | *Chalcolestes parvidens* | M | Friuli-Venezia Giulia | Gorizia | 45.8 | 13.6 | NE | Z17 | this study |
| ZPLOD147-20 | BOLD:ADR7794 | MIB:ZPL:07947 | MT298309 | Zygoptera | Lestidae | *Chalcolestes parvidens* | M | Emilia-Romagna | Bologna | 44.6 | 11.1 | NE | Z17 | this study |
| ZPLOD164-20 | BOLD:AAI7225 | MIB:ZPL:07964 | MT298313 | Zygoptera | Lestidae | *Chalcolestes viridis* | F | Sardinia | Sud Sardegna | 39.4 | 8.7 | SA | Z19 | this study |
| ZPLOD165-20 | BOLD:AAI7225 | MIB:ZPL:07965 | MT298314 | Zygoptera | Lestidae | *Chalcolestes viridis* | M | Sardinia | Nuoro | 40.3 | 9.5 | SA | Z20 | this study |
| ZPLOD167-20 | BOLD:AAI7225 | MIB:ZPL:07967 | MT298315 | Zygoptera | Lestidae | *Chalcolestes viridis* | F | Apulia | Bari | 40.9 | 16.4 | S | Z21 | this study |
| ZPLOD168-20 | BOLD:ADR7794 | MIB:ZPL:07968 | MT298312 | Zygoptera | Lestidae | *Chalcolestes viridis* | M | Apulia | Bari | 40.9 | 16.4 | S | Z22 | this study |
| ZPLOD173-20 | BOLD:AAI7225 | MIB:ZPL:07973 | MT298316 | Zygoptera | Lestidae | *Chalcolestes viridis* | F | Trentino-Alto Adige | Trento | 45.9 | 11.1 | NE | Z23 | this study |
| ZPLOD174-20 | BOLD:AAI7225 | MIB:ZPL:07974 | MT298317 | Zygoptera | Lestidae | *Chalcolestes viridis* | M | Umbria | Perugia | 42.9 | 12.3 | C | Z19 | this study |
| ZPLOD177-20 | BOLD:AAI7225 | MIB:ZPL:07977 | MT298318 | Zygoptera | Lestidae | *Chalcolestes viridis* | F | Friuli-Venezia Giulia | Udine | 45.8 | 13.1 | NE | Z19 | this study |
| ZPLOD179-20 | BOLD:AAI7225 | MIB:ZPL:07979 | MT298319 | Zygoptera | Lestidae | *Chalcolestes viridis* | M | Sicily | Catania | 37.9 | 14.9 | SI | Z19 | this study |
| ZPLOD181-20 | BOLD:AAI7225 | MIB:ZPL:07981 | MT298320 | Zygoptera | Lestidae | *Chalcolestes viridis* | M | Lombardy | Como | 45.7 | 8.9 | NW | Z24 | this study |
| ZPLOD186-20 | BOLD:AAI7225 | MIB:ZPL:07986 | MT298321 | Zygoptera | Lestidae | *Chalcolestes viridis* | M | Sicily | Trapani | 37.9 | 12.8 | SI | Z19 | this study |
| _ | _ | _ | KP272422 | Zygoptera | Coenagrionidae | *Coenagrion caerulescens* | _ | _ | _ | _ | _ | _ | Z29 | NCBI GenBank |
| ZPLOD189-20 | BOLD:ADK6267 | MIB:ZPL:07989 | MT298328 | Zygoptera | Coenagrionidae | *Coenagrion caerulescens* | M | Sicily | Agrigento | 37.5 | 13.2 | SI | Z27 | this study |
| ZPLOD190-20 | BOLD:ADK6267 | MIB:ZPL:07990 | MT298329 | Zygoptera | Coenagrionidae | *Coenagrion caerulescens* | M | Sardinia | Sassari | 40.4 | 8.6 | SA | Z28 | this study |
| ZPLOD192-20 | BOLD:ADK6267 | MIB:ZPL:07992 | MT298325 | Zygoptera | Coenagrionidae | *Coenagrion caerulescens* | M | Apulia | Bari | 40.9 | 16.6 | S | Z26 | this study |
| ZPLOD194-20 | BOLD:ADK6267 | MIB:ZPL:07994 | MT298326 | Zygoptera | Coenagrionidae | *Coenagrion caerulescens* | M | Piedmont | Cuneo | 44.5 | 7.7 | NW | Z30 | this study |
| ZPLOD195-20 | BOLD:ADK6267 | MIB:ZPL:07995 | MT298327 | Zygoptera | Coenagrionidae | *Coenagrion caerulescens* | M | Piedmont | Cuneo | 44.5 | 7.7 | NW | Z31 | this study |
| ZPLOD196-20 | BOLD:ADK6267 | MIB:ZPL:07996 | MT298322 | Zygoptera | Coenagrionidae | *Coenagrion caerulescens* | M | Piedmont | Cuneo | 44.5 | 7.7 | NW | Z32 | this study |
| ZPLOD197-20 | BOLD:ADK6267 | MIB:ZPL:07997 | MT298323 | Zygoptera | Coenagrionidae | *Coenagrion caerulescens* | M | Piedmont | Cuneo | 44.5 | 7.7 | NW | Z32 | this study |
| ZPLOD198-20 | BOLD:ADK6267 | MIB:ZPL:07998 | MT298324 | Zygoptera | Coenagrionidae | *Coenagrion caerulescens* | M | Molise | Campobasso | 41.6 | 14.8 | S | Z26 | this study |
| ZPLOD201-20 | BOLD:ACH0316 | MIB:ZPL:08001 | MT298331 | Zygoptera | Coenagrionidae | *Coenagrion hastulatum* | M | Trentino-Alto Adige | Trento | 46.3 | 11.4 | NE | Z34 | this study |
| ZPLOD202-20 | BOLD:ACH0316 | MIB:ZPL:08002 | MT298330 | Zygoptera | Coenagrionidae | *Coenagrion hastulatum* | M | Friuli-Venezia Giulia | Udine | 46.6 | 13.3 | NE | Z35 | this study |
| _ | _ | _ | KP272414 | Zygoptera | Coenagrionidae | *Coenagrion mercuriale* | _ | _ | _ | _ | _ | _ | Z38 | NCBI GenBank |
| _ | _ | _ | KP272415 | Zygoptera | Coenagrionidae | *Coenagrion mercuriale* | _ | _ | _ | _ | _ | _ | Z38 | NCBI GenBank |
| _ | _ | _ | KP272416 | Zygoptera | Coenagrionidae | *Coenagrion mercuriale* | _ | _ | _ | _ | _ | _ | Z38 | NCBI GenBank |
| _ | _ | _ | KP272417 | Zygoptera | Coenagrionidae | *Coenagrion mercuriale* | _ | _ | _ | _ | _ | _ | Z38 | NCBI GenBank |
| _ | _ | _ | KP272418 | Zygoptera | Coenagrionidae | *Coenagrion mercuriale* | _ | _ | _ | _ | _ | _ | Z38 | NCBI GenBank |
| _ | _ | _ | KP272419 | Zygoptera | Coenagrionidae | *Coenagrion mercuriale* | _ | _ | _ | _ | _ | _ | Z38 | NCBI GenBank |
| _ | _ | _ | KP272420 | Zygoptera | Coenagrionidae | *Coenagrion mercuriale* | _ | _ | _ | _ | _ | _ | Z38 | NCBI GenBank |
| _ | _ | _ | KP272424 | Zygoptera | Coenagrionidae | *Coenagrion mercuriale* | _ | _ | _ | _ | _ | _ | Z38 | NCBI GenBank |
| _ | _ | _ | KP272425 | Zygoptera | Coenagrionidae | *Coenagrion mercuriale* | _ | _ | _ | _ | _ | _ | Z38 | NCBI GenBank |
| _ | _ | _ | KP272448 | Zygoptera | Coenagrionidae | *Coenagrion mercuriale* | _ | _ | _ | _ | _ | _ | Z39 | NCBI GenBank |

MOLECULAR ECOLOGY
RESOURCES

| BOLD PROCESS ID | BIN | VOUCHER ID | GENBANK a.n. | Suborder | Family | SPECIES | SEX | ADMINISTRATIVE REGION | ADMINISTRATIVE PROVINCE | Coordinate N, WGS84 | Coordinate E, WGS84 | MACRO-REGION | HAPLOTYPE | SOURCE |
|---|---|---|---|---|---|---|---|---|---|---|---|---|---|---|
| _ | _ | _ | KP272449 | Zygoptera | Coenagrionidae | *Coenagrion mercuriale* | _ | _ | _ | _ | _ | _ | Z39 | NCBI GenBank |
| _ | _ | _ | KP272450 | Zygoptera | Coenagrionidae | *Coenagrion mercuriale* | _ | _ | _ | _ | _ | _ | Z40 | NCBI GenBank |
| _ | _ | _ | KP272456 | Zygoptera | Coenagrionidae | *Coenagrion mercuriale* | _ | _ | _ | _ | _ | _ | Z39 | NCBI GenBank |
| _ | _ | _ | KP272457 | Zygoptera | Coenagrionidae | *Coenagrion mercuriale* | _ | _ | _ | _ | _ | _ | Z39 | NCBI GenBank |
| _ | _ | _ | KP272505 | Zygoptera | Coenagrionidae | *Coenagrion mercuriale* | _ | _ | _ | _ | _ | _ | Z38 | NCBI GenBank |
| _ | _ | _ | KP272506 | Zygoptera | Coenagrionidae | *Coenagrion mercuriale* | _ | _ | _ | _ | _ | _ | Z38 | NCBI GenBank |
| _ | _ | _ | KP272507 | Zygoptera | Coenagrionidae | *Coenagrion mercuriale* | _ | _ | _ | _ | _ | _ | Z38 | NCBI GenBank |
| _ | _ | _ | KP272551 | Zygoptera | Coenagrionidae | *Coenagrion mercuriale* | _ | _ | _ | _ | _ | _ | Z38 | NCBI GenBank |
| _ | _ | _ | KP272552 | Zygoptera | Coenagrionidae | *Coenagrion mercuriale* | _ | _ | _ | _ | _ | _ | Z38 | NCBI GenBank |
| _ | _ | _ | KP272553 | Zygoptera | Coenagrionidae | *Coenagrion mercuriale* | _ | _ | _ | _ | _ | _ | Z38 | NCBI GenBank |
| _ | _ | _ | KP272554 | Zygoptera | Coenagrionidae | *Coenagrion mercuriale* | _ | _ | _ | _ | _ | _ | Z41 | NCBI GenBank |
| _ | _ | _ | KP272555 | Zygoptera | Coenagrionidae | *Coenagrion mercuriale* | _ | _ | _ | _ | _ | _ | Z38 | NCBI GenBank |
| _ | _ | _ | KP272556 | Zygoptera | Coenagrionidae | *Coenagrion mercuriale* | _ | _ | _ | _ | _ | _ | Z38 | NCBI GenBank |
| _ | _ | _ | KP272557 | Zygoptera | Coenagrionidae | *Coenagrion mercuriale* | _ | _ | _ | _ | _ | _ | Z38 | NCBI GenBank |
| _ | _ | _ | KP272558 | Zygoptera | Coenagrionidae | *Coenagrion mercuriale* | _ | _ | _ | _ | _ | _ | Z38 | NCBI GenBank |
| _ | _ | _ | KP272559 | Zygoptera | Coenagrionidae | *Coenagrion mercuriale* | _ | Emilia-Romagna | Ravenna | _ | _ | NE | Z38 | NCBI GenBank |
| _ | _ | _ | KP272560 | Zygoptera | Coenagrionidae | *Coenagrion mercuriale* | _ | _ | _ | _ | _ | _ | Z38 | NCBI GenBank |
| _ | _ | _ | KX241512 | Zygoptera | Coenagrionidae | *Coenagrion mercuriale* | _ | Lazio | Roma | _ | _ | C | Z36 | NCBI GenBank |
| _ | _ | _ | KX241513 | Zygoptera | Coenagrionidae | *Coenagrion mercuriale* | _ | Lazio | Roma | _ | _ | C | Z36 | NCBI GenBank |
| _ | _ | _ | KX241514 | Zygoptera | Coenagrionidae | *Coenagrion mercuriale* | _ | Lazio | Roma | _ | _ | C | Z42 | NCBI GenBank |
| ZPLOD203-20 | BOLD:ADK3511 | MIB:ZPL:08003 | MT298338 | Zygoptera | Coenagrionidae | *Coenagrion mercuriale* | M | Marches | Macerata | 43.3 | 13.6 | C | Z36 | this study |
| ZPLOD204-20 | BOLD:ADK3511 | MIB:ZPL:08004 | MT298339 | Zygoptera | Coenagrionidae | *Coenagrion mercuriale* | M | Apulia | Lecce | 40.1 | 18.5 | S | Z36 | this study |
| ZPLOD207-20 | BOLD:ADK3511 | MIB:ZPL:08007 | MT298336 | Zygoptera | Coenagrionidae | *Coenagrion mercuriale* | F | Apulia | Barletta-Andria-Trani | 41 | 16.2 | S | Z36 | this study |
| ZPLOD208-20 | BOLD:ADK3511 | MIB:ZPL:08008 | MT298333 | Zygoptera | Coenagrionidae | *Coenagrion mercuriale* | M | Umbria | Perugia | 42.9 | 12.7 | C | Z37 | this study |
| ZPLOD209-20 | BOLD:ADK3511 | MIB:ZPL:08009 | MT298337 | Zygoptera | Coenagrionidae | *Coenagrion mercuriale* | M | Piedmont | Cuneo | 44.6 | 7.7 | NW | Z36 | this study |
| ZPLOD210-20 | BOLD:ADK3511 | MIB:ZPL:08010 | MT298334 | Zygoptera | Coenagrionidae | *Coenagrion mercuriale* | M | Piedmont | Cuneo | 44.5 | 7.7 | NW | Z36 | this study |
| ZPLOD211-20 | BOLD:ADK3511 | MIB:ZPL:08011 | MT298335 | Zygoptera | Coenagrionidae | *Coenagrion mercuriale* | M | Lazio | Latina | 41.5 | 13 | C | Z37 | this study |
| ZPLOD212-20 | BOLD:AAJ0782 | MIB:ZPL:08012 | MT298340 | Zygoptera | Coenagrionidae | *Coenagrion ornatum* | M | Apulia | Bari | 40.8 | 16.4 | S | Z43 | this study |
| ZPLOD213-20 | BOLD:AAJ0782 | MIB:ZPL:08013 | MT298341 | Zygoptera | Coenagrionidae | *Coenagrion ornatum* | M | Basilicata | Potenza | 40.9 | 16.1 | S | Z43 | this study |
| ZPLOD214-20 | BOLD:AAJ0782 | MIB:ZPL:08014 | MT298342 | Zygoptera | Coenagrionidae | *Coenagrion ornatum* | M | Apulia | Barletta-Andria-Trani | 41 | 16.2 | S | Z43 | this study |
| ZPLOD218-20 | BOLD:AAJ0782 | MIB:ZPL:08018 | MT298353 | Zygoptera | Coenagrionidae | *Coenagrion puella* | M | Lombardy | Como | 46.2 | 9.4 | NW | Z44 | this study |
| ZPLOD220-20 | BOLD:AAJ0782 | MIB:ZPL:08020 | MT298354 | Zygoptera | Coenagrionidae | *Coenagrion puella* | M | Calabria | Cosenza | 39.7 | 16 | S | Z45 | this study |
| ZPLOD221-20 | BOLD:AAJ0782 | MIB:ZPL:08021 | MT298355 | Zygoptera | Coenagrionidae | *Coenagrion puella* | M | Trentino-Alto Adige | Trento | 45.7 | 10.9 | NE | Z46 | this study |
| ZPLOD222-20 | BOLD:AAJ0782 | MIB:ZPL:08022 | MT298356 | Zygoptera | Coenagrionidae | *Coenagrion puella* | M | Trentino-Alto Adige | Trento | 45.9 | 11.3 | NE | Z45 | this study |
| ZPLOD223-20 | BOLD:AAJ0782 | MIB:ZPL:08023 | MT298357 | Zygoptera | Coenagrionidae | *Coenagrion puella* | M | Umbria | Perugia | 43 | 12.6 | C | Z45 | this study |
| ZPLOD227-20 | BOLD:AAJ0782 | MIB:ZPL:08027 | MT298358 | Zygoptera | Coenagrionidae | *Coenagrion puella* | M | Sicily | Messina | 38 | 14.7 | SI | Z49 | this study |
| ZPLOD228-20 | BOLD:AAJ0782 | MIB:ZPL:08028 | MT298359 | Zygoptera | Coenagrionidae | *Coenagrion puella* | M | Sicily | Messina | 38 | 14.7 | SI | Z49 | this study |
| ZPLOD230-20 | BOLD:AAJ0782 | MIB:ZPL:08030 | MT298360 | Zygoptera | Coenagrionidae | *Coenagrion puella* | M | Apulia | Bari | 40.9 | 16.4 | S | Z45 | this study |
| ZPLOD231-20 | BOLD:AAJ0782 | MIB:ZPL:08031 | MT298361 | Zygoptera | Coenagrionidae | *Coenagrion puella* | M | Lombardy | Lecco | 45.8 | 9.3 | NW | Z45 | this study |
| ZPLOD232-20 | BOLD:AAJ0782 | MIB:ZPL:08032 | MT298362 | Zygoptera | Coenagrionidae | *Coenagrion puella* | M | Lombardy | Como | 45.8 | 9.2 | NW | Z51 | this study |
| ZPLOD233-20 | BOLD:AAJ0782 | MIB:ZPL:08033 | MT298363 | Zygoptera | Coenagrionidae | *Coenagrion puella* | M | Emilia-Romagna | Piacenza | 44.6 | 9.5 | NW | Z52 | this study |
| ZPLOD234-20 | BOLD:AAJ0782 | MIB:ZPL:08034 | MT298364 | Zygoptera | Coenagrionidae | *Coenagrion puella* | F | Lombardy | Pavia | 44.9 | 9.2 | NW | Z53 | this study |
| ZPLOD237-20 | BOLD:AAJ0782 | MIB:ZPL:08037 | MT298365 | Zygoptera | Coenagrionidae | *Coenagrion puella* | M | Friuli-Venezia Giulia | Trieste | 45.64 | 13.86 | NE | Z45 | this study |
| ZPLOD238-20 | BOLD:AAJ0782 | MIB:ZPL:08038 | MT298345 | Zygoptera | Coenagrionidae | *Coenagrion puella* | F | Piedmont | Vercelli | 45.2 | 8.2 | NW | Z45 | this study |
| ZPLOD239-20 | BOLD:AAJ0782 | MIB:ZPL:08039 | MT298346 | Zygoptera | Coenagrionidae | *Coenagrion puella* | M | Emilia-Romagna | Modena | 44.4 | 10.9 | NW | Z45 | this study |
| ZPLOD240-20 | BOLD:AAJ0782 | MIB:ZPL:08040 | MT298343 | Zygoptera | Coenagrionidae | *Coenagrion puella* | M | Piedmont | Cuneo | 44.6 | 7.7 | NW | Z45 | this study |
| ZPLOD241-20 | BOLD:AAJ0782 | MIB:ZPL:08041 | MT298347 | Zygoptera | Coenagrionidae | *Coenagrion puella* | M | Molise | Campobasso | 41.6 | 14.8 | S | Z45 | this study |
| ZPLOD243-20 | BOLD:AAJ0782 | MIB:ZPL:08043 | MT298348 | Zygoptera | Coenagrionidae | *Coenagrion puella* | M | Abruzzi | L`Aquila | 42.3 | 13.6 | C | Z45 | this study |
| ZPLOD245-20 | BOLD:AAJ0782 | MIB:ZPL:08045 | MT298349 | Zygoptera | Coenagrionidae | *Coenagrion puella* | M | Lombardy | Como | 45.7 | 9.2 | NW | Z51 | this study |
| ZPLOD246-20 | BOLD:AAJ0782 | MIB:ZPL:08046 | MT298350 | Zygoptera | Coenagrionidae | *Coenagrion puella* | M | Emilia-Romagna | Modena | 44.2 | 10.8 | NW | Z53 | this study |
| ZPLOD249-20 | BOLD:AAJ0782 | MIB:ZPL:08049 | MT298351 | Zygoptera | Coenagrionidae | *Coenagrion puella* | M | Lazio | Viterbo | 42.2 | 12 | C | Z45 | this study |
| ZPLOD251-20 | BOLD:AAJ0782 | MIB:ZPL:08051 | MT298352 | Zygoptera | Coenagrionidae | *Coenagrion puella* | M | Lazio | Viterbo | 42.2 | 12 | C | Z53 | this study |
| ZPLOD252-20 | BOLD:AAJ0782 | MIB:ZPL:08052 | MT298376 | Zygoptera | Coenagrionidae | *Coenagrion pulchellum* | M | Lombardy | Como | 46.2 | 9.4 | NW | Z57 | this study |
| ZPLOD253-20 | BOLD:AAJ0782 | MIB:ZPL:08053 | MT298377 | Zygoptera | Coenagrionidae | *Coenagrion pulchellum* | M | Lombardy | Como | 46.2 | 9.4 | NW | Z45 | this study |
| ZPLOD254-20 | BOLD:AAJ0782 | MIB:ZPL:08054 | MT298378 | Zygoptera | Coenagrionidae | *Coenagrion pulchellum* | F | Trentino-Alto Adige | Trento | 46.1 | 11.2 | NE | Z45 | this study |

MOLECULAR ECOLOGY RESOURCES

| BOLD PROCESS ID | BIN | VOUCHER ID | GENBANK a.n. | Suborder | Family | SPECIES | SEX | ADMINISTRATIVE REGION | ADMINISTRATIVE PROVINCE | Coordinate N, WGS84 | Coordinate E, WGS84 | MACRO-REGION | HAPLOTYPE | SOURCE |
|---|---|---|---|---|---|---|---|---|---|---|---|---|---|---|
| ZPLOD255-20 | BOLD:AAJ0782 | MIB:ZPL:08055 | MT298372 | Zygoptera | Coenagrionidae | *Coenagrion pulchellum* | M | Trentino-Alto Adige | Trento | 46.1 | 11.2 | NE | Z57 | this study |
| ZPLOD256-20 | BOLD:AAJ0782 | MIB:ZPL:08056 | MT298373 | Zygoptera | Coenagrionidae | *Coenagrion pulchellum* | M | Umbria | Terni | 42.5 | 12.8 | C | Z58 | this study |
| ZPLOD257-20 | BOLD:AAJ0782 | MIB:ZPL:08057 | MT298374 | Zygoptera | Coenagrionidae | *Coenagrion pulchellum* | M | Apulia | Foggia | 41.6 | 15.9 | S | Z58 | this study |
| ZPLOD258-20 | BOLD:AAJ0782 | MIB:ZPL:08058 | MT298375 | Zygoptera | Coenagrionidae | *Coenagrion pulchellum* | F | Apulia | Lecce | 40.2 | 18.4 | S | Z58 | this study |
| ZPLOD259-20 | BOLD:AAJ0782 | MIB:ZPL:08059 | MT298371 | Zygoptera | Coenagrionidae | *Coenagrion pulchellum* | M | Lombardy | Como | 45.8 | 9.2 | NW | Z57 | this study |
| ZPLOD260-20 | BOLD:AAJ0782 | MIB:ZPL:08060 | MT298366 | Zygoptera | Coenagrionidae | *Coenagrion pulchellum* | M | Lazio | Latina | 41.5 | 13.1 | C | Z58 | this study |
| ZPLOD261-20 | BOLD:AAJ0782 | MIB:ZPL:08061 | MT298367 | Zygoptera | Coenagrionidae | *Coenagrion pulchellum* | F | Lazio | Latina | 41.5 | 13.1 | C | Z58 | this study |
| ZPLOD262-20 | BOLD:AAJ0782 | MIB:ZPL:08062 | MT298368 | Zygoptera | Coenagrionidae | *Coenagrion pulchellum* | M | Lazio | Latina | 41.5 | 13.1 | C | Z59 | this study |
| ZPLOD263-20 | BOLD:AAJ0782 | MIB:ZPL:08063 | MT298369 | Zygoptera | Coenagrionidae | *Coenagrion pulchellum* | M | Lazio | Latina | 41.2 | 13.1 | C | Z58 | this study |
| ZPLOD264-20 | BOLD:AAJ0782 | MIB:ZPL:08064 | MT298370 | Zygoptera | Coenagrionidae | *Coenagrion pulchellum* | M | Lazio | Latina | 41.5 | 13.1 | C | Z58 | this study |
| _ | _ | _ | KP272421 | Zygoptera | Coenagrionidae | *Coenagrion scitulum* | _ | _ | _ | _ | _ | _ | Z61 | NCBI GenBank |
| _ | _ | _ | KP272423 | Zygoptera | Coenagrionidae | *Coenagrion scitulum* | _ | _ | _ | _ | _ | _ | Z61 | NCBI GenBank |
| ZPLOD266-20 | BOLD:ACP4983 | MIB:ZPL:08066 | MT298379 | Zygoptera | Coenagrionidae | *Coenagrion scitulum* | M | Apulia | Bari | 40.9 | 16.4 | S | Z60 | this study |
| ZPLOD272-20 | BOLD:ACP4983 | MIB:ZPL:08072 | MT298380 | Zygoptera | Coenagrionidae | *Coenagrion scitulum* | M | Lombardy | Varese | 45.7 | 9 | NW | Z62 | this study |
| ZPLOD273-20 | BOLD:ACP4983 | MIB:ZPL:08073 | MT298381 | Zygoptera | Coenagrionidae | *Coenagrion scitulum* | M | Friuli-Venezia Giulia | Trieste | 45.64 | 13.86 | NE | Z62 | this study |
| ZPLOD275-20 | BOLD:ACP4983 | MIB:ZPL:08075 | MT298382 | Zygoptera | Coenagrionidae | *Coenagrion scitulum* | M | Molise | Campobasso | 41.8 | 15 | S | Z63 | this study |
| ZPLOD278-20 | BOLD:ACP4983 | MIB:ZPL:08078 | MT298383 | Zygoptera | Coenagrionidae | *Coenagrion scitulum* | M | Lazio | Viterbo | 42.2 | 12 | C | Z62 | this study |
| _ | _ | _ | KF584929 | Anisoptera | Cordulegastridae | *Cordulegaster bidentata* | _ | Basilicata | Potenza | _ | _ | S | A38 | NCBI GenBank |
| _ | _ | _ | KF584930 | Anisoptera | Cordulegastridae | *Cordulegaster bidentata* | _ | Basilicata | Potenza | _ | _ | S | A39 | NCBI GenBank |
| _ | _ | _ | KF584947 | Anisoptera | Cordulegastridae | *Cordulegaster bidentata* | _ | Sicily | Messina | _ | _ | SI | A40 | NCBI GenBank |
| _ | _ | _ | KF584971 | Anisoptera | Cordulegastridae | *Cordulegaster bidentata* | _ | Sicily | Messina | _ | _ | SI | A41 | NCBI GenBank |
| _ | _ | _ | KF584972 | Anisoptera | Cordulegastridae | *Cordulegaster bidentata* | _ | Sicily | Siracusa | _ | _ | SI | A42 | NCBI GenBank |
| ZPLOD287-20 | BOLD:AAJ5749 | MIB:ZPL:08087 | MT298384 | Anisoptera | Cordulegastridae | *Cordulegaster bidentata* | M | Trentino-Alto Adige | Trento | 46.2 | 11.3 | NE | A36 | this study |
| ZPLOD288-20 | BOLD:AAJ5749 | MIB:ZPL:08088 | MT298386 | Anisoptera | Cordulegastridae | *Cordulegaster bidentata* | M | Piedmont | Cuneo | 44.2 | 7.7 | NW | A37 | this study |
| ZPLOD289-20 | BOLD:AAJ5749 | MIB:ZPL:08089 | MT298385 | Anisoptera | Cordulegastridae | *Cordulegaster bidentata* | M | Lombardy | Lecco | 45.9 | 9.3 | NW | A36 | this study |
| _ | _ | _ | KF584933 | Anisoptera | Cordulegastridae | *Cordulegaster boltonii* | _ | Tuscany | Firenze | _ | _ | C | A45 | NCBI GenBank |
| _ | _ | _ | KF584934 | Anisoptera | Cordulegastridae | *Cordulegaster boltonii* | _ | Tuscany | Firenze | _ | _ | C | A46 | NCBI GenBank |
| _ | _ | _ | MH304646 | Anisoptera | Cordulegastridae | *Cordulegaster boltonii* | _ | Piedmont | Alessandria | _ | _ | NW | A45 | NCBI GenBank |
| _ | _ | _ | MH304646 | Anisoptera | Cordulegastridae | *Cordulegaster boltonii* | _ | Piedmont | Asti | _ | _ | NW | A45 | NCBI GenBank |
| _ | _ | _ | MH304646 | Anisoptera | Cordulegastridae | *Cordulegaster boltonii* | _ | Marches | Macerata | _ | _ | C | A45 | NCBI GenBank |
| _ | _ | _ | MH304646 | Anisoptera | Cordulegastridae | *Cordulegaster boltonii* | _ | Emilia-Romagna | Ravenna | _ | _ | NE | A45 | NCBI GenBank |
| _ | _ | _ | MH304647 | Anisoptera | Cordulegastridae | *Cordulegaster boltonii* | _ | Piedmont | Asti | _ | _ | NW | A47 | NCBI GenBank |
| _ | _ | _ | MH304647 | Anisoptera | Cordulegastridae | *Cordulegaster boltonii* | _ | Lombardy | Brescia | _ | _ | NW | A47 | NCBI GenBank |
| _ | _ | _ | MH304647 | Anisoptera | Cordulegastridae | *Cordulegaster boltonii* | _ | Trentino-Alto Adige | Trento | _ | _ | NE | A47 | NCBI GenBank |
| _ | _ | _ | MH304648 | Anisoptera | Cordulegastridae | *Cordulegaster boltonii* | _ | _ | _ | _ | _ | _ | A48 | NCBI GenBank |
| _ | _ | _ | MH304649 | Anisoptera | Cordulegastridae | *Cordulegaster boltonii* | _ | Piedmont | Alessandria | _ | _ | NW | A49 | NCBI GenBank |
| _ | _ | _ | MH304650 | Anisoptera | Cordulegastridae | *Cordulegaster boltonii* | _ | Tuscany | Firenze | _ | _ | C | A46 | NCBI GenBank |
| _ | _ | _ | MH304650 | Anisoptera | Cordulegastridae | *Cordulegaster boltonii* | _ | Emilia-Romagna | Ravenna | _ | _ | NE | A46 | NCBI GenBank |
| ZPLOD295-20 | BOLD:AAJ5773 | MIB:ZPL:08095 | MT298389 | Anisoptera | Cordulegastridae | *Cordulegaster boltonii* | M | Trentino-Alto Adige | Trento | 46.1 | 11 | NE | A43 | this study |
| ZPLOD297-20 | BOLD:AAJ5773 | MIB:ZPL:08097 | MT298387 | Anisoptera | Cordulegastridae | *Cordulegaster boltonii* | M | Piedmont | Torino | 45.1 | 7.5 | NW | A44 | this study |
| ZPLOD299-20 | BOLD:AAJ5773 | MIB:ZPL:08099 | MT298388 | Anisoptera | Cordulegastridae | *Cordulegaster boltonii* | M | Friuli-Venezia Giulia | Udine | 46.2 | 13 | NE | A43 | this study |
| ZPLOD302-20 | BOLD:ACQ4796 | MIB:ZPL:08102 | MT298393 | Anisoptera | Cordulegastridae | *Cordulegaster heros* | M | Friuli-Venezia Giulia | Gorizia | 45.9 | 13.6 | NE | A50 | this study |
| ZPLOD303-20 | BOLD:ACQ4796 | MIB:ZPL:08103 | MT298392 | Anisoptera | Cordulegastridae | *Cordulegaster heros* | M | Friuli-Venezia Giulia | Trieste | 45.64 | 13.86 | NE | A51 | this study |
| ZPLOD304-20 | BOLD:ACQ4796 | MIB:ZPL:08104 | MT298391 | Anisoptera | Cordulegastridae | *Cordulegaster heros* | M | Friuli-Venezia Giulia | Udine | 46.2 | 13.3 | NE | A51 | this study |
| _ | _ | _ | KF584945 | Anisoptera | Cordulegastridae | *Cordulegaster trinacriae* | _ | Sicily | _ | _ | _ | SI | A54 | NCBI GenBank |
| _ | _ | _ | KF584946 | Anisoptera | Cordulegastridae | *Cordulegaster trinacriae* | _ | _ | _ | _ | _ | _ | A55 | NCBI GenBank |
| _ | _ | _ | MH304651 | Anisoptera | Cordulegastridae | *Cordulegaster trinacriae* | _ | Basilicata | Potenza | _ | _ | S | A56 | NCBI GenBank |
| _ | _ | _ | MH304653 | Anisoptera | Cordulegastridae | *Cordulegaster trinacriae* | _ | Sicily | _ | _ | _ | SI | A55 | NCBI GenBank |
| _ | _ | _ | MH304654 | Anisoptera | Cordulegastridae | *Cordulegaster trinacriae* | _ | Sicily | Messina | _ | _ | SI | A57 | NCBI GenBank |
| _ | _ | _ | MH304656 | Anisoptera | Cordulegastridae | *Cordulegaster trinacriae* | _ | _ | _ | _ | _ | _ | A58 | NCBI GenBank |
| _ | _ | _ | MH304658 | Anisoptera | Cordulegastridae | *Cordulegaster trinacriae* | _ | Sicily | Siracusa | _ | _ | SI | A59 | NCBI GenBank |
| _ | _ | _ | MH304661 | Anisoptera | Cordulegastridae | *Cordulegaster trinacriae* | _ | Calabria | Vibo Valentia | _ | _ | S | A60 | NCBI GenBank |
| _ | _ | _ | MH304662 | Anisoptera | Cordulegastridae | *Cordulegaster trinacriae* | _ | Calabria | Vibo Valentia | _ | _ | S | A61 | NCBI GenBank |
| _ | _ | _ | MH304663 | Anisoptera | Cordulegastridae | *Cordulegaster trinacriae* | _ | Basilicata | Potenza | _ | _ | S | A62 | NCBI GenBank |
| _ | _ | _ | MH304664 | Anisoptera | Cordulegastridae | *Cordulegaster trinacriae* | _ | Apulia | Barletta-Andria-Trani | _ | _ | S | A63 | NCBI GenBank |

MOLECULAR ECOLOGY RESOURCES

| BOLD PROCESS ID | BIN | VOUCHER ID | GENBANK a.n. | Suborder | Family | SPECIES | SEX | ADMINISTRATIVE REGION | ADMINISTRATIVE PROVINCE | Coordinate N, WGS84 | Coordinate E, WGS84 | MACRO-REGION | HAPLOTYPE | SOURCE |
|---|---|---|---|---|---|---|---|---|---|---|---|---|---|---|
| _ | _ | _ | MH304665 | Anisoptera | Cordulegastridae | *Cordulegaster trinacriae* | _ | Basilicata | Potenza | _ | _ | S | A64 | NCBI GenBank |
| ZPLOD314-20 | BOLD:ACQ2278 | MIB:ZPL:08114 | MT298395 | Anisoptera | Cordulegastridae | *Cordulegaster trinacriae* | M | Apulia | Bari | 40.8 | 16.4 | S | A53 | this study |
| ZPLOD315-20 | BOLD:ACQ2278 | MIB:ZPL:08115 | MT298394 | Anisoptera | Cordulegastridae | *Cordulegaster trinacriae* | M | Molise | Campobasso | 41.7 | 14.9 | S | A65 | this study |
| ZPLOD316-20 | BOLD:AAJ5771 | MIB:ZPL:08116 | MT298396 | Anisoptera | Anisoptera | *Cordulia aenea* | M | Trentino-Alto Adige | Trento | 46.1 | 11.2 | NE | A66 | this study |
| ZPLOD317-20 | BOLD:AAJ5771 | MIB:ZPL:08117 | MT298397 | Anisoptera | Anisoptera | *Cordulia aenea* | M | Umbria | Terni | 42.5 | 12.8 | C | A66 | this study |
| ZPLOD319-20 | BOLD:AAJ5771 | MIB:ZPL:08119 | MT298398 | Anisoptera | Anisoptera | *Cordulia aenea* | M | Lombardy | Varese | 45.9 | 8.9 | NW | A68 | this study |
| ZPLOD320-20 | BOLD:AAI9726 | MIB:ZPL:08120 | MT298405 | Anisoptera | Libellulidae | *Crocothemis erythraea* | M | Sardinia | Sassari | 40.7 | 9 | SA | A69 | this study |
| ZPLOD328-20 | BOLD:AAI9726 | MIB:ZPL:08128 | MT298400 | Anisoptera | Libellulidae | *Crocothemis erythraea* | M | Lombardy | Monza Brianza | 45.7 | 9.1 | NW | A70 | this study |
| ZPLOD331-20 | BOLD:AAI9726 | MIB:ZPL:08131 | MT298401 | Anisoptera | Libellulidae | *Crocothemis erythraea* | M | Molise | Campobasso | 42 | 15 | S | A71 | this study |
| ZPLOD332-20 | BOLD:AAI9726 | MIB:ZPL:08132 | MT298402 | Anisoptera | Libellulidae | *Crocothemis erythraea* | M | Lazio | Latina | 41.5 | 13.1 | C | A72 | this study |
| ZPLOD333-20 | BOLD:AAI9726 | MIB:ZPL:08133 | MT298403 | Anisoptera | Libellulidae | *Crocothemis erythraea* | M | Sicily | Trapani | 37.9 | 12.7 | SI | A73 | this study |
| ZPLOD334-20 | BOLD:ABU6643 | MIB:ZPL:08134 | MT298407 | Anisoptera | Libellulidae | *Diplacodes lefebvrii* | M | Sardinia | Sud Sardegna | 39.2 | 8.2 | SA | A74 | this study |
| ZPLOD335-20 | BOLD:ABU6643 | MIB:ZPL:08135 | MT298406 | Anisoptera | Libellulidae | *Diplacodes lefebvrii* | M | Sardinia | Sud Sardegna | 39.2 | 8.2 | SA | A75 | this study |
| ZPLOD336-20 | BOLD:AAA2218 | MIB:ZPL:08136 | MT298408 | Zygoptera | Coenagrionidae | *Enallagma cyathigerum* | M | Aosta Valley | Aosta | 45.8 | 7.6 | NW | Z65 | this study |
| ZPLOD343-20 | BOLD:AAA2218 | MIB:ZPL:08143 | MT298409 | Zygoptera | Coenagrionidae | *Enallagma cyathigerum* | M | Sicily | Messina | 37.9 | 14.7 | SI | Z66 | this study |
| ZPLOD345-20 | BOLD:AAA2218 | MIB:ZPL:08145 | MT298413 | Zygoptera | Coenagrionidae | *Enallagma cyathigerum* | M | Lombardy | Lecco | 45.8 | 9.4 | NW | Z67 | this study |
| ZPLOD346-20 | BOLD:AAA2218 | MIB:ZPL:08146 | MT298412 | Zygoptera | Coenagrionidae | *Enallagma cyathigerum* | F | Emilia-Romagna | Piacenza | 44.6 | 9.5 | NW | Z65 | this study |
| ZPLOD348-20 | BOLD:AAA2218 | MIB:ZPL:08148 | MT298411 | Zygoptera | Coenagrionidae | *Enallagma cyathigerum* | F | Emilia-Romagna | Modena | 44.5 | 10.9 | NW | Z69 | this study |
| ZPLOD351-20 | BOLD:AAA2218 | MIB:ZPL:08151 | MT298410 | Zygoptera | Coenagrionidae | *Enallagma cyathigerum* | M | Abruzzi | L`Aquila | 42.3 | 13.6 | C | Z66 | this study |
| ZPLOD352-20 | BOLD:AAL4439 | MIB:ZPL:08152 | MT298414 | Zygoptera | Coenagrionidae | *Erythromma lindenii* | M | Sicily | Agrigento | 37.5 | 13.2 | SI | Z71 | this study |
| ZPLOD354-20 | BOLD:AAL4439 | MIB:ZPL:08154 | MT298449 | Zygoptera | Coenagrionidae | *Erythromma lindenii* | M | Sicily | Trapani | 37.7 | 12.8 | SI | Z71 | this study |
| ZPLOD355-20 | BOLD:AAL4439 | MIB:ZPL:08155 | MT298448 | Zygoptera | Coenagrionidae | *Erythromma lindenii* | M | Sicily | Siracusa | 37.1 | 15 | SI | Z71 | this study |
| ZPLOD358-20 | BOLD:AAL4439 | MIB:ZPL:08158 | MT298447 | Zygoptera | Coenagrionidae | *Erythromma lindenii* | M | Sardinia | Sassari | 40.4 | 8.6 | SA | Z72 | this study |
| ZPLOD364-20 | BOLD:AAL4439 | MIB:ZPL:08164 | MT298446 | Zygoptera | Coenagrionidae | *Erythromma lindenii* | M | Sardinia | Nuoro | 40.3 | 9.5 | SA | Z72 | this study |
| ZPLOD365-20 | BOLD:AEC5518 | MIB:ZPL:08165 | MT298445 | Zygoptera | Coenagrionidae | *Erythromma lindenii* | M | Trentino-Alto Adige | Trento | 46 | 11.3 | NE | Z73 | this study |
| ZPLOD366-20 | BOLD:AEC5518 | MIB:ZPL:08166 | MT298444 | Zygoptera | Coenagrionidae | *Erythromma lindenii* | M | Trentino-Alto Adige | Trento | 46 | 11.3 | NE | Z74 | this study |
| ZPLOD367-20 | BOLD:AEC5518 | MIB:ZPL:08167 | MT298443 | Zygoptera | Coenagrionidae | *Erythromma lindenii* | M | Trentino-Alto Adige | Trento | 46 | 11.3 | NE | Z75 | this study |
| ZPLOD373-20 | BOLD:AEC5518 | MIB:ZPL:08173 | MT298442 | Zygoptera | Coenagrionidae | *Erythromma lindenii* | M | Trentino-Alto Adige | Trento | 46.1 | 11.1 | NE | Z74 | this study |
| ZPLOD374-20 | BOLD:AEC5518 | MIB:ZPL:08174 | MT298441 | Zygoptera | Coenagrionidae | *Erythromma lindenii* | M | Trentino-Alto Adige | Trento | 46.1 | 11.1 | NE | Z73 | this study |
| ZPLOD375-20 | BOLD:AEC5518 | MIB:ZPL:08175 | MT298440 | Zygoptera | Coenagrionidae | *Erythromma lindenii* | M | Trentino-Alto Adige | Trento | 46.1 | 11.1 | NE | Z74 | this study |
| ZPLOD376-20 | BOLD:AAL4439 | MIB:ZPL:08176 | MT298439 | Zygoptera | Coenagrionidae | *Erythromma lindenii* | M | Sardinia | Medio Campidano | 39.6 | 9 | SA | Z72 | this study |
| ZPLOD377-20 | BOLD:AAL4439 | MIB:ZPL:08177 | MT298438 | Zygoptera | Coenagrionidae | *Erythromma lindenii* | M | Apulia | Bari | 40.8 | 16.4 | S | Z71 | this study |
| ZPLOD378-20 | BOLD:AAL4439 | MIB:ZPL:08178 | MT298437 | Zygoptera | Coenagrionidae | *Erythromma lindenii* | M | Apulia | Barletta-Andria-Trani | 41 | 16.2 | S | Z77 | this study |
| ZPLOD379-20 | BOLD:AAL4439 | MIB:ZPL:08179 | MT298436 | Zygoptera | Coenagrionidae | *Erythromma lindenii* | M | Apulia | Lecce | 40.3 | 17.8 | S | Z78 | this study |
| ZPLOD380-20 | BOLD:AEC5518 | MIB:ZPL:08180 | MT298435 | Zygoptera | Coenagrionidae | *Erythromma lindenii* | M | Apulia | Lecce | 40.5 | 18.2 | S | Z79 | this study |
| ZPLOD381-20 | BOLD:AEC5518 | MIB:ZPL:08181 | MT298434 | Zygoptera | Coenagrionidae | *Erythromma lindenii* | M | Trentino-Alto Adige | Trento | 46 | 11.3 | NE | Z73 | this study |
| ZPLOD383-20 | BOLD:AAL4439 | MIB:ZPL:08183 | MT298433 | Zygoptera | Coenagrionidae | *Erythromma lindenii* | M | Umbria | Perugia | 42.7 | 12.6 | C | Z80 | this study |
| ZPLOD384-20 | BOLD:AEC5518 | MIB:ZPL:08184 | MT298432 | Zygoptera | Coenagrionidae | *Erythromma lindenii* | _ | Piedmont | Vercelli | 45.2 | 8.2 | NW | Z74 | this study |
| ZPLOD385-20 | BOLD:AEC5518 | MIB:ZPL:08185 | MT298431 | Zygoptera | Coenagrionidae | *Erythromma lindenii* | M | Friuli-Venezia Giulia | Udine | 45.8 | 13.2 | NE | Z81 | this study |
| ZPLOD386-20 | BOLD:AEC5518 | MIB:ZPL:08186 | MT298430 | Zygoptera | Coenagrionidae | *Erythromma lindenii* | M | Friuli-Venezia Giulia | Udine | 45.8 | 13.1 | NE | Z74 | this study |
| ZPLOD387-20 | BOLD:AAL4439 | MIB:ZPL:08187 | MT298429 | Zygoptera | Coenagrionidae | *Erythromma lindenii* | M | Friuli-Venezia Giulia | Trieste | 45.64 | 13.86 | NE | Z83 | this study |
| ZPLOD388-20 | BOLD:AEC5518 | MIB:ZPL:08188 | MT298428 | Zygoptera | Coenagrionidae | *Erythromma lindenii* | M | Friuli-Venezia Giulia | Pordenone | 46 | 12.9 | NE | Z81 | this study |
| ZPLOD389-20 | BOLD:AEC5518 | MIB:ZPL:08189 | MT298427 | Zygoptera | Coenagrionidae | *Erythromma lindenii* | M | Lombardy | Brescia | 45.7 | 10.5 | NW | Z73 | this study |
| ZPLOD390-20 | BOLD:AEC5518 | MIB:ZPL:08190 | MT298426 | Zygoptera | Coenagrionidae | *Erythromma lindenii* | M | Lombardy | Brescia | 45.7 | 10.5 | NW | Z73 | this study |
| ZPLOD391-20 | BOLD:AAL4439 | MIB:ZPL:08191 | MT298425 | Zygoptera | Coenagrionidae | *Erythromma lindenii* | M | Sicily | Catania | 37.5 | 14.9 | SI | Z71 | this study |
| ZPLOD392-20 | BOLD:AAL4439 | MIB:ZPL:08192 | MT298424 | Zygoptera | Coenagrionidae | *Erythromma lindenii* | M | Lombardy | Lecco | 45.8 | 9.4 | NW | Z85 | this study |
| ZPLOD393-20 | BOLD:AAL4439 | MIB:ZPL:08193 | MT298423 | Zygoptera | Coenagrionidae | *Erythromma lindenii* | M | Lombardy | Lecco | 45.8 | 9.4 | NW | Z71 | this study |
| ZPLOD394-20 | BOLD:AAL4439 | MIB:ZPL:08194 | MT298422 | Zygoptera | Coenagrionidae | *Erythromma lindenii* | M | Lombardy | Milano | 45.3 | 9 | NW | Z86 | this study |
| ZPLOD395-20 | BOLD:AAL4439 | MIB:ZPL:08195 | MT298421 | Zygoptera | Coenagrionidae | *Erythromma lindenii* | M | Emilia-Romagna | Modena | 44.5 | 10.9 | NW | Z71 | this study |
| ZPLOD396-20 | BOLD:AAL4439 | MIB:ZPL:08196 | MT298420 | Zygoptera | Coenagrionidae | *Erythromma lindenii* | M | Lombardy | Varese | 45.7 | 8.7 | NW | Z71 | this study |
| ZPLOD397-20 | BOLD:AAL4439 | MIB:ZPL:08197 | MT298419 | Zygoptera | Coenagrionidae | *Erythromma lindenii* | M | Molise | Campobasso | 41.9 | 15 | S | Z71 | this study |
| ZPLOD399-20 | BOLD:AAL4439 | MIB:ZPL:08199 | MT298418 | Zygoptera | Coenagrionidae | *Erythromma lindenii* | M | Molise | Campobasso | 41.6 | 14.8 | S | Z88 | this study |
| ZPLOD401-20 | BOLD:AAL4439 | MIB:ZPL:08201 | MT298417 | Zygoptera | Coenagrionidae | *Erythromma lindenii* | M | Abruzzi | L`Aquila | 42.3 | 13.6 | C | Z71 | this study |
| ZPLOD406-20 | BOLD:AAA4234 | MIB:ZPL:08206 | MT298450 | Zygoptera | Coenagrionidae | *Erythromma najas* | M | Trentino-Alto Adige | Trento | 46 | 11 | NE | Z89 | this study |
| ZPLOD407-20 | BOLD:AAA4234 | MIB:ZPL:08207 | MT298451 | Zygoptera | Coenagrionidae | *Erythromma najas* | M | Lombardy | Varese | 45.8 | 8.8 | NW | Z90 | this study |

MOLECULAR ECOLOGY RESOURCES

| BOLD PROCESS ID | BIN | VOUCHER ID | GENBANK a.n. | Suborder | Family | SPECIES | SEX | ADMINISTRATIVE REGION | ADMINISTRATIVE PROVINCE | Coordinate N, WGS84 | Coordinate E, WGS84 | MACRO-REGION | HAPLOTYPE | SOURCE |
|---|---|---|---|---|---|---|---|---|---|---|---|---|---|---|
| ZPLOD409-20 | BOLD:AAL4437 | MIB:ZPL:08209 | MT298458 | Zygoptera | Coenagrionidae | *Erythromma viridulum* | M | Sardinia | Sassari | 40.7 | 8.2 | SA | Z91 | this study |
| ZPLOD412-20 | BOLD:AAL4437 | MIB:ZPL:08212 | MT298457 | Zygoptera | Coenagrionidae | *Erythromma viridulum* | F | Apulia | Lecce | 40.3 | 17.8 | S | Z92 | this study |
| ZPLOD414-20 | BOLD:AAL4437 | MIB:ZPL:08214 | MT298454 | Zygoptera | Coenagrionidae | *Erythromma viridulum* | M | Trentino-Alto Adige | Trento | 46 | 11.3 | NE | Z92 | this study |
| ZPLOD417-20 | BOLD:AAL4437 | MIB:ZPL:08217 | MT298455 | Zygoptera | Coenagrionidae | *Erythromma viridulum* | M | Sicily | Siracusa | 37.1 | 15.3 | SI | Z93 | this study |
| ZPLOD419-20 | BOLD:AAL4437 | MIB:ZPL:08219 | MT298456 | Zygoptera | Coenagrionidae | *Erythromma viridulum* | M | Lombardy | Lecco | 45.8 | 9.4 | NW | Z94 | this study |
| ZPLOD421-20 | BOLD:AAL4437 | MIB:ZPL:08221 | MT298453 | Zygoptera | Coenagrionidae | *Erythromma viridulum* | M | Abruzzi | L`Aquila | 42.3 | 13.6 | C | Z92 | this study |
| ZPLOD422-20 | BOLD:AAL4437 | MIB:ZPL:08222 | MT298452 | Zygoptera | Coenagrionidae | *Erythromma viridulum* | M | Sicily | Trapani | 37.9 | 12.7 | SI | Z95 | this study |
| ZPLOD424-20 | BOLD:AAN0925 | MIB:ZPL:08224 | MT298461 | Anisoptera | Gomphidae | *Gomphus vulgatissimus* | M | Marches | Ancona | 43.4 | 13 | C | A76 | this study |
| ZPLOD425-20 | BOLD:AAN0925 | MIB:ZPL:08225 | MT298462 | Anisoptera | Gomphidae | *Gomphus vulgatissimus* | F | Friuli-Venezia Giulia | Udine | 45.8 | 13.1 | NE | A76 | this study |
| ZPLOD426-20 | BOLD:AAN0925 | MIB:ZPL:08226 | MT298463 | Anisoptera | Gomphidae | *Gomphus vulgatissimus* | M | Lombardy | Como | 45.8 | 9.2 | NW | A76 | this study |
| ZPLOD430-20 | BOLD:AAE5570 | MIB:ZPL:08230 | MT298465 | Zygoptera | Coenagrionidae | *Ischnura elegans* | M | Trentino-Alto Adige | Trento | 46.2 | 11.1 | NE | Z96 | this study |
| ZPLOD431-20 | BOLD:AAE5570 | MIB:ZPL:08231 | MT298466 | Zygoptera | Coenagrionidae | *Ischnura elegans* | M | Piedmont | Torino | 45.1 | 7.5 | NW | Z97 | this study |
| ZPLOD438-20 | BOLD:AAE5570 | MIB:ZPL:08238 | MT298467 | Zygoptera | Coenagrionidae | *Ischnura elegans* | M | Apulia | Lecce | 40.2 | 18.4 | S | Z98 | this study |
| ZPLOD444-20 | BOLD:AAE5570 | MIB:ZPL:08244 | MT298468 | Zygoptera | Coenagrionidae | *Ischnura elegans* | M | Tuscany | Grosseto | 43 | 10.9 | C | Z97 | this study |
| ZPLOD447-20 | BOLD:AAE5570 | MIB:ZPL:08247 | MT298469 | Zygoptera | Coenagrionidae | *Ischnura elegans* | M | Lazio | Viterbo | 42.2 | 12 | C | Z97 | this study |
| ZPLOD452-20 | BOLD:AAE5570 | MIB:ZPL:08252 | MT298470 | Zygoptera | Coenagrionidae | *Ischnura genei* | M | Sardinia | Sassari | 40.7 | 9 | SA | Z99 | this study |
| ZPLOD454-20 | BOLD:AAE5570 | MIB:ZPL:08254 | MT298471 | Zygoptera | Coenagrionidae | *Ischnura genei* | M | Sicily | Siracusa | 37.1 | 15.3 | SI | Z100 | this study |
| ZPLOD457-20 | BOLD:AAE5570 | MIB:ZPL:08257 | MT298472 | Zygoptera | Coenagrionidae | *Ischnura genei* | M | Sicily | Trapani | 37.9 | 12.7 | SI | Z101 | this study |
| ZPLOD460-20 | BOLD:AAE5571 | MIB:ZPL:08260 | MT298473 | Zygoptera | Coenagrionidae | *Ischnura pumilio* | M | Apulia | Bari | 40.7 | 16.7 | S | Z102 | this study |
| ZPLOD461-20 | BOLD:AAE5571 | MIB:ZPL:08261 | MT298476 | Zygoptera | Coenagrionidae | *Ischnura pumilio* | F | Trentino-Alto Adige | Trento | 46.1 | 11 | NE | Z102 | this study |
| ZPLOD462-20 | BOLD:AAE5571 | MIB:ZPL:08262 | MT298475 | Zygoptera | Coenagrionidae | *Ischnura pumilio* | M | Piedmont | Torino | 45.1 | 7.5 | NW | Z102 | this study |
| ZPLOD466-20 | BOLD:AAE5571 | MIB:ZPL:08266 | MT298474 | Zygoptera | Coenagrionidae | *Ischnura pumilio* | M | Emilia-Romagna | Modena | 44.9 | 11.2 | NW | Z102 | this study |
| ZPLOD468-20 | BOLD:ADC3442 | MIB:ZPL:08268 | MT298481 | Zygoptera | Lestidae | *Lestes barbarus* | M | Sardinia | Sassari | 40.9 | 8.2 | SA | Z104 | this study |
| ZPLOD474-20 | BOLD:ADC3442 | MIB:ZPL:08274 | MT298482 | Zygoptera | Lestidae | *Lestes barbarus* | M | Apulia | Taranto | 40.6 | 16.8 | S | Z105 | this study |
| ZPLOD475-20 | BOLD:ADC3442 | MIB:ZPL:08275 | MT298479 | Zygoptera | Lestidae | *Lestes barbarus* | F | Emilia-Romagna | Ferrara | 44.8 | 12 | NE | Z106 | this study |
| ZPLOD476-20 | BOLD:ADC3442 | MIB:ZPL:08276 | MT298480 | Zygoptera | Lestidae | *Lestes barbarus* | M | Molise | Campobasso | 41.6 | 14.8 | S | Z104 | this study |
| ZPLOD478-20 | BOLD:AEC4388 | MIB:ZPL:08278 | MT298486 | Zygoptera | Lestidae | *Lestes dryas* | F | Liguria | Savona | 44.4 | 8.6 | NW | Z107 | this study |
| ZPLOD481-20 | BOLD:AEC4388 | MIB:ZPL:08281 | MT298487 | Zygoptera | Lestidae | *Lestes dryas* | M | Apulia | Bari | 40.9 | 16.5 | S | Z108 | this study |
| ZPLOD482-20 | BOLD:AEC4388 | MIB:ZPL:08282 | MT298488 | Zygoptera | Lestidae | *Lestes dryas* | M | Piedmont | Torino | 44.9 | 6.8 | NW | Z109 | this study |
| ZPLOD484-20 | BOLD:AEC4388 | MIB:ZPL:08284 | MT298489 | Zygoptera | Lestidae | *Lestes dryas* | F | Sicily | Messina | 37.9 | 14.7 | SI | Z110 | this study |
| ZPLOD485-20 | BOLD:AEC4388 | MIB:ZPL:08285 | MT298484 | Zygoptera | Lestidae | *Lestes dryas* | M | Emilia-Romagna | Piacenza | 44.6 | 9.5 | NW | Z111 | this study |
| ZPLOD486-20 | BOLD:AEC4388 | MIB:ZPL:08286 | MT298485 | Zygoptera | Lestidae | *Lestes dryas* | M | Abruzzi | L`Aquila | 42.3 | 13.6 | C | Z112 | this study |
| ZPLOD488-20 | BOLD:ADC3318 | MIB:ZPL:08288 | MT298490 | Zygoptera | Lestidae | *Lestes macrostigma* | F | Apulia | Lecce | 40.3 | 17.8 | S | Z113 | this study |
| ZPLOD491-20 | BOLD:ACP4984 | MIB:ZPL:08291 | MT298493 | Zygoptera | Lestidae | *Lestes sponsa* | M | Aosta Valley | Aosta | 45.9 | 7.6 | NW | Z114 | this study |
| ZPLOD492-20 | BOLD:ACP4984 | MIB:ZPL:08292 | MT298492 | Zygoptera | Lestidae | *Lestes sponsa* | M | Trentino-Alto Adige | Trento | 46.3 | 11.4 | NE | Z115 | this study |
| ZPLOD496-20 | BOLD:ACP4984 | MIB:ZPL:08296 | MT298491 | Zygoptera | Lestidae | *Lestes sponsa* | M | Lombardy | Lecco | 45.8 | 9.4 | NW | Z116 | this study |
| ZPLOD499-20 | BOLD:ACG0123 | MIB:ZPL:08299 | MT298497 | Zygoptera | Lestidae | *Lestes virens* | M | Sicily | Catania | 37.9 | 14.9 | SI | Z119 | this study |
| ZPLOD500-20 | BOLD:ACG0123 | MIB:ZPL:08300 | MT298496 | Zygoptera | Lestidae | *Lestes virens* | M | Sicily | Messina | 38 | 14.7 | SI | Z119 | this study |
| ZPLOD501-20 | BOLD:ACG0123 | MIB:ZPL:08301 | MT298495 | Zygoptera | Lestidae | *Lestes virens* | M | Sicily | Trapani | 37.9 | 12.7 | SI | Z119 | this study |
| ZPLOD503-20 | BOLD:ACG0123 | MIB:ZPL:08303 | MT298501 | Zygoptera | Lestidae | *Lestes virens* | M | Piedmont | Torino | 45.1 | 7.5 | NW | Z118 | this study |
| ZPLOD504-20 | BOLD:ACG0123 | MIB:ZPL:08304 | MT298494 | Zygoptera | Lestidae | *Lestes virens* | M | Lombardy | Como | 45.7 | 9.2 | NW | Z121 | this study |
| ZPLOD505-20 | BOLD:ACG0123 | MIB:ZPL:08305 | MT298500 | Zygoptera | Lestidae | *Lestes virens* | F | Emilia-Romagna | Piacenza | 44.6 | 9.5 | NW | Z117 | this study |
| ZPLOD509-20 | BOLD:ACG0123 | MIB:ZPL:08309 | MT298499 | Zygoptera | Lestidae | *Lestes virens* | M | Molise | Campobasso | 41.6 | 14.8 | S | Z117 | this study |
| ZPLOD512-20 | BOLD:ACG0123 | MIB:ZPL:08312 | MT298498 | Zygoptera | Lestidae | *Lestes virens* | M | Sardinia | Sud Sardegna | 39.7 | 8.9 | SA | Z117 | this study |
| ZPLOD515-20 | BOLD:AAJ2437 | MIB:ZPL:08315 | MT298502 | Anisoptera | Libellulidae | *Leucorrhinia dubia* | M | Aosta Valley | Aosta | 45.8 | 7.6 | NW | A78 | this study |
| ZPLOD520-20 | BOLD:AAJ2437 | MIB:ZPL:08320 | MT298506 | Anisoptera | Libellulidae | *Leucorrhinia dubia* | _ | Trentino-Alto Adige | Trento | 45.1 | 7.2 | NE | A79 | this study |
| ZPLOD522-20 | BOLD:AAJ2437 | MIB:ZPL:08322 | MT298504 | Anisoptera | Libellulidae | *Leucorrhinia dubia* | M | Piedmont | Verbano-Cusio-Ossola | 46.3 | 8.3 | NW | A79 | this study |
| ZPLOD523-20 | BOLD:AAJ2437 | MIB:ZPL:08323 | MT298503 | Anisoptera | Libellulidae | *Leucorrhinia dubia* | M | Lombardy | Sondrio | 46.4 | 9.4 | NW | A79 | this study |
| ZPLOD524-20 | BOLD:ADC3719 | MIB:ZPL:08324 | MT298507 | Anisoptera | Libellulidae | *Leucorrhinia pectoralis* | M | Trentino-Alto Adige | Trento | 46.1 | 11.2 | NE | A80 | this study |
| ZPLOD526-20 | BOLD:AAJ2758 | MIB:ZPL:08326 | MT298510 | Anisoptera | Libellulidae | *Libellula depressa* | M | Apulia | Barletta-Andria-Trani | 41 | 16.2 | S | A81 | this study |
| ZPLOD527-20 | BOLD:AAJ2758 | MIB:ZPL:08327 | MT298509 | Anisoptera | Libellulidae | *Libellula depressa* | M | Trentino-Alto Adige | Trento | 45.7 | 10.9 | NE | A82 | this study |
| ZPLOD528-20 | BOLD:AAJ2758 | MIB:ZPL:08328 | MT298512 | Anisoptera | Libellulidae | *Libellula depressa* | M | Piedmont | Torino | 45.1 | 7.5 | NW | A82 | this study |
| ZPLOD531-20 | BOLD:AAJ2758 | MIB:ZPL:08331 | MT298508 | Anisoptera | Libellulidae | *Libellula depressa* | M | Emilia-Romagna | Piacenza | 44.6 | 9.5 | NW | A82 | this study |
| ZPLOD534-20 | BOLD:AAJ2758 | MIB:ZPL:08334 | MT298511 | Anisoptera | Libellulidae | *Libellula depressa* | F | Molise | Campobasso | 41.8 | 14.5 | S | A84 | this study |
| ZPLOD535-20 | BOLD:ACP3530 | MIB:ZPL:08335 | MT298514 | Anisoptera | Libellulidae | *Libellula fulva* | M | Apulia | Lecce | 40.5 | 18.2 | S | A85 | this study |

MOLECULAR ECOLOGY RESOURCES

| BOLD PROCESS ID | BIN | VOUCHER ID | GENBANK a.n. | Suborder | Family | SPECIES | SEX | ADMINISTRATIVE REGION | ADMINISTRATIVE PROVINCE | Coordinate N, WGS84 | Coordinate E, WGS84 | MACRO-REGION | HAPLOTYPE | SOURCE |
|---|---|---|---|---|---|---|---|---|---|---|---|---|---|---|
| ZPLOD536-20 | BOLD:ACP3530 | MIB:ZPL:08336 | MT298515 | Anisoptera | Libellulidae | *Libellula fulva* | M | Trentino-Alto Adige | Trento | 46.2 | 11.1 | NE | A86 | this study |
| ZPLOD541-20 | BOLD:ACP3530 | MIB:ZPL:08341 | MT298513 | Anisoptera | Libellulidae | *Libellula fulva* | F | Lombardy | Lecco | 45.8 | 9.4 | NW | A87 | this study |
| ZPLOD543-20 | BOLD:ACP3530 | MIB:ZPL:08343 | MT298516 | Anisoptera | Libellulidae | *Libellula fulva* | M | Lazio | Latina | 41.5 | 13.1 | C | A88 | this study |
| ZPLOD544-20 | BOLD:AAB5337 | MIB:ZPL:08344 | MT298521 | Anisoptera | Libellulidae | *Libellula quadrimaculata* | M | Aosta Valley | Aosta | 45.9 | 7.6 | NW | A89 | this study |
| ZPLOD545-20 | BOLD:AAB5337 | MIB:ZPL:08345 | MT298518 | Anisoptera | Libellulidae | *Libellula quadrimaculata* | M | Trentino-Alto Adige | Trento | 46.3 | 11.4 | NE | A90 | this study |
| ZPLOD547-20 | BOLD:AAB5337 | MIB:ZPL:08347 | MT298520 | Anisoptera | Libellulidae | *Libellula quadrimaculata* | M | Umbria | Terni | 42.5 | 12.8 | C | A90 | this study |
| ZPLOD549-20 | BOLD:AAB5337 | MIB:ZPL:08349 | MT298519 | Anisoptera | Libellulidae | *Libellula quadrimaculata* | M | Lombardy | Monza Brianza | 45.7 | 9.1 | NW | A91 | this study |
| _ | _ | _ | KX241516 | Anisoptera | Gomphidae | *Lindenia tetraphylla* | _ | Tuscany | Siena | _ | _ | C | A94 | NCBI GenBank |
| ZPLOD550-20 | BOLD:ACW0675 | MIB:ZPL:08350 | MT298525 | Anisoptera | Gomphidae | *Lindenia tetraphylla* | F | Sardinia | Oristano | 40 | 8.5 | SA | A92 | this study |
| ZPLOD552-20 | BOLD:ACW0675 | MIB:ZPL:08352 | MT298524 | Anisoptera | Gomphidae | *Lindenia tetraphylla* | F | Umbria | Perugia | 43.1 | 12.2 | C | A93 | this study |
| ZPLOD553-20 | BOLD:ACW0675 | MIB:ZPL:08353 | MT298526 | Anisoptera | Gomphidae | *Lindenia tetraphylla* | M | Molise | Campobasso | 42 | 14.8 | S | A93 | this study |
| ZPLOD555-20 | BOLD:ACW0675 | MIB:ZPL:08355 | MT298527 | Anisoptera | Gomphidae | *Lindenia tetraphylla* | F | Molise | Campobasso | 41.6 | 14.9 | S | A93 | this study |
| ZPLOD556-20 | BOLD:ACW0675 | MIB:ZPL:08356 | MT298523 | Anisoptera | Gomphidae | *Lindenia tetraphylla* | _ | Sicily | Trapani | 37.7 | 12.8 | SI | A92 | this study |
| ZPLOD557-20 | BOLD:AAC3125 | MIB:ZPL:08357 | MT298529 | Zygoptera | Coenagrionidae | *Nehalennia speciosa* | F | Friuli-Venezia Giulia | Udine | 46.1 | 13.2 | NE | Z122 | this study |
| ZPLOD560-20 | BOLD:AAC3125 | MIB:ZPL:08360 | MT298528 | Zygoptera | Coenagrionidae | *Nehalennia speciosa* | M | Lombardy | Varese | 45.9 | 8.9 | NW | Z123 | this study |
| ZPLOD562-20 | BOLD:AAE5061 | MIB:ZPL:08362 | MT298541 | Anisoptera | Gomphidae | *Onychogomphus forcipatus* | M | Friuli-Venezia Giulia | Pordenone | 46.2 | 12.9 | NE | A102 | this study |
| ZPLOD563-20 | BOLD:AAE5061 | MIB:ZPL:08363 | MT298540 | Anisoptera | Gomphidae | *Onychogomphus forcipatus* | M | Friuli-Venezia Giulia | Gorizia | 46 | 13.6 | NE | A103 | this study |
| ZPLOD564-20 | BOLD:AAE5061 | MIB:ZPL:08364 | MT298542 | Anisoptera | Gomphidae | *Onychogomphus forcipatus* | M | Sicily | Messina | 38 | 15.3 | SI | A104 | this study |
| ZPLOD565-20 | BOLD:AAE5061 | MIB:ZPL:08365 | MT298543 | Anisoptera | Gomphidae | *Onychogomphus forcipatus* | M | Sicily | Messina | 38 | 15.3 | SI | A104 | this study |
| ZPLOD566-20 | BOLD:AAE5061 | MIB:ZPL:08366 | MT298544 | Anisoptera | Gomphidae | *Onychogomphus forcipatus* | M | Trentino-Alto Adige | Trento | 46.1 | 11.2 | NE | A96 | this study |
| ZPLOD567-20 | BOLD:AAE5061 | MIB:ZPL:08367 | MT298539 | Anisoptera | Gomphidae | *Onychogomphus forcipatus* | M | Trentino-Alto Adige | Trento | 46.1 | 11.2 | NE | A97 | this study |
| ZPLOD568-20 | BOLD:AAE5061 | MIB:ZPL:08368 | MT298531 | Anisoptera | Gomphidae | *Onychogomphus forcipatus* | M | Trentino-Alto Adige | Trento | 46 | 11.3 | NE | A96 | this study |
| ZPLOD569-20 | BOLD:AAE5061 | MIB:ZPL:08369 | MT298532 | Anisoptera | Gomphidae | *Onychogomphus forcipatus* | M | Trentino-Alto Adige | Trento | 45.9 | 11 | NE | A97 | this study |
| ZPLOD570-20 | BOLD:AAE5061 | MIB:ZPL:08370 | MT298533 | Anisoptera | Gomphidae | *Onychogomphus forcipatus* | M | Trentino-Alto Adige | Trento | 46.1 | 11.1 | NE | A96 | this study |
| ZPLOD571-20 | BOLD:AAE5061 | MIB:ZPL:08371 | MT298534 | Anisoptera | Gomphidae | *Onychogomphus forcipatus* | M | Trentino-Alto Adige | Trento | 46.1 | 11.1 | NE | A97 | this study |
| ZPLOD576-20 | BOLD:AAE5061 | MIB:ZPL:08376 | MT298535 | Anisoptera | Gomphidae | *Onychogomphus forcipatus* | M | Lombardy | Lecco | 45.8 | 9.4 | NW | A96 | this study |
| ZPLOD578-20 | BOLD:AAE5061 | MIB:ZPL:08378 | MT298536 | Anisoptera | Gomphidae | *Onychogomphus forcipatus* | M | Apulia | Taranto | 40.6 | 16.8 | S | A96 | this study |
| ZPLOD579-20 | BOLD:AAE5061 | MIB:ZPL:08379 | MT298537 | Anisoptera | Gomphidae | *Onychogomphus forcipatus* | M | Emilia-Romagna | Modena | 44.5 | 10.9 | NW | A96 | this study |
| ZPLOD580-20 | BOLD:AAE5061 | MIB:ZPL:08380 | MT298538 | Anisoptera | Gomphidae | *Onychogomphus forcipatus* | M | Molise | Campobasso | 41.9 | 15 | S | A96 | this study |
| ZPLOD581-20 | BOLD:ADC3114 | MIB:ZPL:08381 | MT298548 | Anisoptera | Gomphidae | *Onychogomphus uncatus* | M | Sicily | Siracusa | 37.1 | 15 | SI | A107 | this study |
| ZPLOD582-20 | BOLD:ADC3114 | MIB:ZPL:08382 | MT298547 | Anisoptera | Gomphidae | *Onychogomphus uncatus* | M | Liguria | Genova | 44.5 | 8.8 | NW | A108 | this study |
| ZPLOD585-20 | BOLD:ADC3114 | MIB:ZPL:08385 | MT298546 | Anisoptera | Gomphidae | *Onychogomphus uncatus* | M | Lombardy | Milano | 45.3 | 9 | NW | A108 | this study |
| ZPLOD587-20 | BOLD:ADC3114 | MIB:ZPL:08387 | MT298545 | Anisoptera | Gomphidae | *Onychogomphus uncatus* | M | Molise | Campobasso | 41.6 | 14.6 | S | A108 | this study |
| ZPLOD588-20 | BOLD:ACP4340 | MIB:ZPL:08388 | MT298549 | Anisoptera | Gomphidae | *Ophiogomphus cecilia* | M | Lombardy | Pavia | 45.2 | 8.9 | NW | A109 | this study |
| ZPLOD589-20 | BOLD:ACQ8102 | MIB:ZPL:08389 | MT298550 | Anisoptera | Libellulidae | *Orthetrum albistylum* | M | Piedmont | Torino | 45.1 | 7.5 | NW | A110 | this study |
| ZPLOD594-20 | BOLD:ACQ8102 | MIB:ZPL:08394 | MT298551 | Anisoptera | Libellulidae | *Orthetrum albistylum* | F | Lombardy | Milano | 45.3 | 9 | NW | A110 | this study |
| ZPLOD598-20 | BOLD:AAK5997 | MIB:ZPL:08398 | MT298552 | Anisoptera | Libellulidae | *Orthetrum brunneum* | F | Liguria | Spezia | 44.1 | 10 | NW | A112 | this study |
| ZPLOD600-20 | BOLD:AAK5997 | MIB:ZPL:08400 | MT298553 | Anisoptera | Libellulidae | *Orthetrum brunneum* | M | Trentino-Alto Adige | Trento | 46 | 11.3 | NE | A114 | this study |
| ZPLOD603-20 | BOLD:AAK5997 | MIB:ZPL:08403 | MT298558 | Anisoptera | Libellulidae | *Orthetrum brunneum* | M | Umbria | Perugia | 42.8 | 13.1 | C | A114 | this study |
| ZPLOD605-20 | BOLD:AAK5997 | MIB:ZPL:08405 | MT298559 | Anisoptera | Libellulidae | *Orthetrum brunneum* | _ | Sicily | Messina | 38 | 15.3 | SI | A115 | this study |
| ZPLOD609-20 | BOLD:AAK5997 | MIB:ZPL:08409 | MT298554 | Anisoptera | Libellulidae | *Orthetrum brunneum* | M | Sicily | Trapani | 37.9 | 12.7 | SI | A116 | this study |
| ZPLOD610-20 | BOLD:AAK5997 | MIB:ZPL:08410 | MT298555 | Anisoptera | Libellulidae | *Orthetrum brunneum* | M | Sardinia | Sassari | 40.7 | 9 | SA | A111 | this study |
| ZPLOD611-20 | BOLD:AAK5997 | MIB:ZPL:08411 | MT298556 | Anisoptera | Libellulidae | *Orthetrum brunneum* | M | Sardinia | Sassari | 40.4 | 8.6 | SA | A112 | this study |
| ZPLOD612-20 | BOLD:AAK5997 | MIB:ZPL:08412 | MT298557 | Anisoptera | Libellulidae | *Orthetrum brunneum* | M | Sardinia | Sud Sardegna | 39.4 | 8.7 | SA | A112 | this study |
| ZPLOD613-20 | BOLD:AAK5996 | MIB:ZPL:08413 | MT298561 | Anisoptera | Libellulidae | *Orthetrum cancellatum* | M | Sardinia | Sassari | 40.7 | 8.2 | SA | A117 | this study |
| ZPLOD614-20 | BOLD:AAK5996 | MIB:ZPL:08414 | MT298562 | Anisoptera | Libellulidae | *Orthetrum cancellatum* | M | Marches | Macerata | 43.3 | 13.3 | C | A118 | this study |
| ZPLOD616-20 | BOLD:AAK5996 | MIB:ZPL:08416 | MT298563 | Anisoptera | Libellulidae | *Orthetrum cancellatum* | F | Trentino-Alto Adige | Trento | 46 | 11.3 | NE | A119 | this study |
| ZPLOD618-20 | BOLD:AAK5996 | MIB:ZPL:08418 | MT298564 | Anisoptera | Libellulidae | *Orthetrum cancellatum* | M | Umbria | Perugia | 42.7 | 12.6 | C | A120 | this study |
| ZPLOD619-20 | BOLD:AAK5996 | MIB:ZPL:08419 | MT298560 | Anisoptera | Libellulidae | *Orthetrum cancellatum* | _ | Sicily | Catania | 37.9 | 14.9 | SI | A121 | this study |
| ZPLOD620-20 | BOLD:AAK5996 | MIB:ZPL:08420 | MT298568 | Anisoptera | Libellulidae | *Orthetrum cancellatum* | M | Apulia | Brindisi | 40.7 | 17.8 | S | A122 | this study |
| ZPLOD623-20 | BOLD:AAK5996 | MIB:ZPL:08423 | MT298567 | Anisoptera | Libellulidae | *Orthetrum cancellatum* | F | Lombardy | Milano | 45.3 | 9 | NW | A123 | this study |
| ZPLOD625-20 | BOLD:AAK5996 | MIB:ZPL:08425 | MT298566 | Anisoptera | Libellulidae | *Orthetrum cancellatum* | _ | Sicily | Trapani | 37.9 | 12.7 | SI | A118 | this study |
| _ | _ | _ | KC912263 | Anisoptera | Libellulidae | *Orthetrum coerulescens* | _ | Lazio | Frosinone | _ | _ | C | A131 | NCBI GenBank |
| _ | _ | _ | KC912264 | Anisoptera | Libellulidae | *Orthetrum coerulescens* | _ | Lazio | Frosinone | _ | _ | C | A131 | NCBI GenBank |
| _ | _ | _ | KC912265 | Anisoptera | Libellulidae | *Orthetrum coerulescens* | _ | Lazio | Frosinone | _ | _ | C | A131 | NCBI GenBank |

MOLECULAR ECOLOGY RESOURCES

| BOLD PROCESS ID | BIN | VOUCHER ID | GENBANK a.n. | Suborder | Family | SPECIES | SEX | ADMINISTRATIVE REGION | ADMINISTRATIVE PROVINCE | Coordinate N, WGS84 | Coordinate E, WGS84 | MACRO-REGION | HAPLOTYPE | SOURCE |
|---|---|---|---|---|---|---|---|---|---|---|---|---|---|---|
| ZPLOD626-20 | BOLD:AAI2353 | MIB:ZPL:08426 | MT298572 | Anisoptera | Libellulidae | *Orthetrum coerulescens* | M | Sicily | Messina | 37.9 | 14.7 | SI | A124 | this study |
| ZPLOD627-20 | BOLD:AAI2353 | MIB:ZPL:08427 | MT298571 | Anisoptera | Libellulidae | *Orthetrum coerulescens* | M | Sicily | Catania | 37.4 | 15.1 | SI | A124 | this study |
| ZPLOD628-20 | BOLD:AAI2353 | MIB:ZPL:08428 | MT298570 | Anisoptera | Libellulidae | *Orthetrum coerulescens* | M | Sicily | Agrigento | 37.5 | 13.2 | SI | A124 | this study |
| ZPLOD632-20 | BOLD:AAI2353 | MIB:ZPL:08432 | MT298582 | Anisoptera | Libellulidae | *Orthetrum coerulescens* | M | Sicily | Trapani | 37.6 | 12.9 | SI | A124 | this study |
| ZPLOD639-20 | BOLD:AAI2353 | MIB:ZPL:08439 | MT298573 | Anisoptera | Libellulidae | *Orthetrum coerulescens* | M | Trentino-Alto Adige | Trento | 46 | 11.3 | NE | A129 | this study |
| ZPLOD640-20 | BOLD:AAI2353 | MIB:ZPL:08440 | MT298574 | Anisoptera | Libellulidae | *Orthetrum coerulescens* | M | Piedmont | Torino | 45.1 | 7.5 | NW | A124 | this study |
| ZPLOD642-20 | BOLD:AAI2353 | MIB:ZPL:08442 | MT298575 | Anisoptera | Libellulidae | *Orthetrum coerulescens* | M | Umbria | Perugia | 42.9 | 12.7 | C | A124 | this study |
| ZPLOD644-20 | BOLD:AAI2353 | MIB:ZPL:08444 | MT298576 | Anisoptera | Libellulidae | *Orthetrum coerulescens* | _ | Sicily | Siracusa | 37.1 | 15.3 | SI | A124 | this study |
| ZPLOD648-20 | BOLD:AAI2353 | MIB:ZPL:08448 | MT298577 | Anisoptera | Libellulidae | *Orthetrum coerulescens* | M | Molise | Campobasso | 42 | 14.8 | S | A124 | this study |
| ZPLOD649-20 | BOLD:AAI2353 | MIB:ZPL:08449 | MT298578 | Anisoptera | Libellulidae | *Orthetrum coerulescens* | M | Lombardy | Como | 45.7 | 9.2 | NW | A124 | this study |
| ZPLOD651-20 | BOLD:AAI2353 | MIB:ZPL:08451 | MT298579 | Anisoptera | Libellulidae | *Orthetrum coerulescens* | M | Lazio | Roma | 41.9 | 12.5 | C | A132 | this study |
| ZPLOD658-20 | BOLD:AAI2353 | MIB:ZPL:08458 | MT298580 | Anisoptera | Libellulidae | *Orthetrum coerulescens* | F | Sardinia | Sassari | 40.5 | 8.6 | SA | A128 | this study |
| ZPLOD659-20 | BOLD:AAI2353 | MIB:ZPL:08459 | MT298581 | Anisoptera | Libellulidae | *Orthetrum coerulescens* | M | Sardinia | Sud Sardegna | 39.4 | 8.7 | SA | A128 | this study |
| ZPLOD661-20 | BOLD:AEC4264 | MIB:ZPL:08461 | MT298584 | Anisoptera | Libellulidae | *Orthetrum nitidinerve* | M | Sardinia | Nuoro | 40.2 | 8.9 | SA | A133 | this study |
| ZPLOD663-20 | BOLD:AEC4264 | MIB:ZPL:08463 | MT298585 | Anisoptera | Libellulidae | *Orthetrum nitidinerve* | M | Sardinia | Sud Sardegna | 39.8 | 9 | SA | A133 | this study |
| ZPLOD665-20 | BOLD:ABA9397 | MIB:ZPL:08465 | MT298586 | Anisoptera | Libellulidae | *Orthetrum trinacria* | M | Sardinia | Sassari | 40.8 | 8.5 | SA | A134 | this study |
| ZPLOD667-20 | BOLD:ABA9397 | MIB:ZPL:08467 | MT298588 | Anisoptera | Libellulidae | *Orthetrum trinacria* | M | Sicily | Catania | 37.5 | 14.9 | SI | A135 | this study |
| ZPLOD668-20 | BOLD:ABA9397 | MIB:ZPL:08468 | MT298587 | Anisoptera | Libellulidae | *Orthetrum trinacria* | M | Sicily | Trapani | 37.7 | 12.8 | SI | A135 | this study |
| _ | _ | _ | KX241515 | Anisoptera | Anisoptera | *Oxygastra curtisii* | _ | Marches | Pesaro-Urbino | _ | _ | C | A136 | NCBI GenBank |
| ZPLOD671-20 | BOLD:ADC4889 | MIB:ZPL:08471 | MT298590 | Anisoptera | Anisoptera | *Oxygastra curtisii* | M | Lombardy | Bergamo | 45.7 | 9.4 | NW | A137 | this study |
| ZPLOD672-20 | BOLD:ADC4889 | MIB:ZPL:08472 | MT298589 | Anisoptera | Anisoptera | *Oxygastra curtisii* | M | Molise | Campobasso | 41.6 | 14.8 | S | A138 | this study |
| ZPLOD674-20 | BOLD:ABW0140 | MIB:ZPL:08474 | MT298593 | Anisoptera | Gomphidae | *Paragomphus genei* | M | Sardinia | Sassari | 40.7 | 9 | SA | A139 | this study |
| ZPLOD675-20 | BOLD:ABW0140 | MIB:ZPL:08475 | MT298592 | Anisoptera | Gomphidae | *Paragomphus genei* | M | Sardinia | Nuoro | 40.3 | 9.5 | SA | A139 | this study |
| ZPLOD676-20 | BOLD:ABW0140 | MIB:ZPL:08476 | MT298591 | Anisoptera | Gomphidae | *Paragomphus genei* | M | Sicily | Agrigento | 37.6 | 13.1 | SI | A141 | this study |
| ZPLOD679-20 | BOLD:ACG0515 | MIB:ZPL:08479 | MT298597 | Zygoptera | Platycnemididae | *Platycnemis pennipes* | M | Apulia | Taranto | 40.662 | 16.799 | S | Z124 | this study |
| ZPLOD680-20 | BOLD:ACG0515 | MIB:ZPL:08480 | MT298596 | Zygoptera | Platycnemididae | *Platycnemis pennipes* | M | Trentino-Alto Adige | Trento | 46 | 11.3 | NE | Z125 | this study |
| ZPLOD681-20 | BOLD:ACG0515 | MIB:ZPL:08481 | MT298595 | Zygoptera | Platycnemididae | *Platycnemis pennipes* | M | Piedmont | Torino | 45.1 | 7.5 | NW | Z125 | this study |
| ZPLOD683-20 | BOLD:ACG0515 | MIB:ZPL:08483 | MT298594 | Zygoptera | Platycnemididae | *Platycnemis pennipes* | _ | Piedmont | Vercelli | 45.2 | 8.1 | NW | Z125 | this study |
| ZPLOD685-20 | BOLD:ACG0515 | MIB:ZPL:08485 | MT298600 | Zygoptera | Platycnemididae | *Platycnemis pennipes* | M | Lombardy | Lecco | 45.8 | 9.3 | NW | Z128 | this study |
| ZPLOD687-20 | BOLD:ACG0515 | MIB:ZPL:08487 | MT298599 | Zygoptera | Platycnemididae | *Platycnemis pennipes* | M | Tuscany | Grosseto | 43 | 10.9 | C | Z125 | this study |
| ZPLOD690-20 | BOLD:ACG0515 | MIB:ZPL:08490 | MT298602 | Zygoptera | Platycnemididae | *Platycnemis pennipes* | M | Molise | Campobasso | 41.6 | 14.8 | S | Z129 | this study |
| ZPLOD691-20 | BOLD:ACG0515 | MIB:ZPL:08491 | MT298601 | Zygoptera | Platycnemididae | *Platycnemis pennipes* | M | Lazio | Viterbo | 42.2 | 12 | C | Z125 | this study |
| ZPLOD695-20 | BOLD:AAD5734 | MIB:ZPL:08495 | MT298604 | Zygoptera | Coenagrionidae | *Pyrrhosoma nymphula* | M | Trentino-Alto Adige | Trento | 46 | 11.6 | NE | Z130 | this study |
| ZPLOD696-20 | BOLD:AAD5734 | MIB:ZPL:08496 | MT298605 | Zygoptera | Coenagrionidae | *Pyrrhosoma nymphula* | M | Umbria | Perugia | 42.9 | 12.7 | C | Z131 | this study |
| ZPLOD698-20 | BOLD:AAD5734 | MIB:ZPL:08498 | MT298603 | Zygoptera | Coenagrionidae | *Pyrrhosoma nymphula* | M | Lombardy | Como | 45.8 | 9.2 | NW | Z130 | this study |
| ZPLOD700-20 | BOLD:AAE9139 | MIB:ZPL:08500 | MT298608 | Anisoptera | Libellulidae | *Selysiothemis nigra* | F | Sicily | Catania | 37.5 | 14.9 | SI | A142 | this study |
| ZPLOD701-20 | BOLD:AAE9139 | MIB:ZPL:08501 | MT298609 | Anisoptera | Libellulidae | *Selysiothemis nigra* | M | Emilia-Romagna | Ferrara | 44.7 | 12.2 | NE | A143 | this study |
| ZPLOD703-20 | BOLD:AAE9139 | MIB:ZPL:08503 | MT298607 | Anisoptera | Libellulidae | *Selysiothemis nigra* | M | Molise | Campobasso | 41.8 | 15 | S | A144 | this study |
| ZPLOD706-20 | BOLD:ACP5227 | MIB:ZPL:08506 | MT298613 | Anisoptera | Anisoptera | *Somatochlora alpestris* | F | Trentino-Alto Adige | Trento | 46.5 | 11.1 | NE | A145 | this study |
| ZPLOD707-20 | BOLD:ACP5227 | MIB:ZPL:08507 | MT298612 | Anisoptera | Anisoptera | *Somatochlora alpestris* | M | Friuli-Venezia Giulia | Udine | 46.5 | 12.7 | NE | A145 | this study |
| ZPLOD709-20 | BOLD:ACP5227 | MIB:ZPL:08509 | MT298611 | Anisoptera | Anisoptera | *Somatochlora alpestris* | M | Lombardy | Sondrio | 46.1 | 9.6 | NW | A147 | this study |
| ZPLOD710-20 | BOLD:ACP5227 | MIB:ZPL:08510 | MT298610 | Anisoptera | Anisoptera | *Somatochlora alpestris* | F | Lombardy | Sondrio | 46.4 | 9.4 | NW | A145 | this study |
| ZPLOD712-20 | BOLD:ACP7013 | MIB:ZPL:08512 | MT298615 | Anisoptera | Anisoptera | *Somatochlora arctica* | M | Trentino-Alto Adige | Trento | 46.1 | 11.2 | NE | A149 | this study |
| ZPLOD713-20 | BOLD:ACP7013 | MIB:ZPL:08513 | MT298614 | Anisoptera | Anisoptera | *Somatochlora arctica* | M | Lombardy | Bergamo | 46 | 9.8 | NW | A150 | this study |
| ZPLOD715-20 | BOLD:AEC6167 | MIB:ZPL:08515 | MT298616 | Anisoptera | Anisoptera | *Somatochlora flavomaculata* | M | Apulia | Lecce | 40.4 | 18.3 | S | A151 | this study |
| ZPLOD717-20 | BOLD:AEC6167 | MIB:ZPL:08517 | MT298618 | Anisoptera | Anisoptera | *Somatochlora flavomaculata* | M | Trentino-Alto Adige | Trento | 46.1 | 11.2 | NE | A152 | this study |
| ZPLOD718-20 | BOLD:AEC6167 | MIB:ZPL:08518 | MT298619 | Anisoptera | Anisoptera | *Somatochlora flavomaculata* | _ | Piedmont | Vercelli | 45.5 | 8.3 | NW | A153 | this study |
| ZPLOD720-20 | BOLD:ABW6681 | MIB:ZPL:08520 | MT298621 | Anisoptera | Anisoptera | *Somatochlora meridionalis* | M | Lazio | Frosinone | 41.5 | 13.3 | C | A154 | this study |
| ZPLOD721-20 | BOLD:ABW6681 | MIB:ZPL:08521 | MT298622 | Anisoptera | Anisoptera | *Somatochlora meridionalis* | M | Marches | Macerata | 43.3 | 13.5 | C | A155 | this study |
| ZPLOD722-20 | BOLD:ABW6681 | MIB:ZPL:08522 | MT298623 | Anisoptera | Anisoptera | *Somatochlora meridionalis* | M | Umbria | Perugia | 42.9 | 12.2 | C | A154 | this study |
| ZPLOD723-20 | BOLD:ABW6681 | MIB:ZPL:08523 | MT298624 | Anisoptera | Anisoptera | *Somatochlora meridionalis* | M | Friuli-Venezia Giulia | Gorizia | 46 | 13.6 | NE | A157 | this study |
| ZPLOD724-20 | BOLD:ABW6681 | MIB:ZPL:08524 | MT298625 | Anisoptera | Anisoptera | *Somatochlora meridionalis* | F | Friuli-Venezia Giulia | Udine | 46.2 | 13.3 | NE | A158 | this study |
| ZPLOD725-20 | BOLD:ABW6681 | MIB:ZPL:08525 | MT298626 | Anisoptera | Anisoptera | *Somatochlora meridionalis* | M | Piedmont | Cuneo | 44.4 | 8.1 | NW | A159 | this study |
| ZPLOD726-20 | BOLD:ABW6681 | MIB:ZPL:08526 | MT298627 | Anisoptera | Anisoptera | *Somatochlora meridionalis* | M | Piedmont | Cuneo | 44.4 | 8.1 | NW | A159 | this study |
| ZPLOD727-20 | BOLD:ABW6681 | MIB:ZPL:08527 | MT298628 | Anisoptera | Anisoptera | *Somatochlora meridionalis* | M | Piedmont | Cuneo | 44.4 | 8.1 | NW | A159 | this study |

MOLECULAR ECOLOGY RESOURCES

| BOLD PROCESS ID | BIN | VOUCHER ID | GENBANK a.n. | Suborder | Family | SPECIES | SEX | ADMINISTRATIVE REGION | ADMINISTRATIVE PROVINCE | Coordinate N, WGS84 | Coordinate E, WGS84 | MACRO-REGION | HAPLOTYPE | SOURCE |
|---|---|---|---|---|---|---|---|---|---|---|---|---|---|---|
| ZPLOD728-20 | BOLD:ABW6681 | MIB:ZPL:08528 | MT298630 | Anisoptera | Anisoptera | *Somatochlora metallica* | M | Trentino-Alto Adige | Trento | 46 | 11.6 | NE | A160 | this study |
| ZPLOD729-20 | BOLD:ABW6681 | MIB:ZPL:08529 | MT298631 | Anisoptera | Anisoptera | *Somatochlora metallica* | M | Lombardy | Pavia | 45.2 | 8.7 | NW | A160 | this study |
| ZPLOD730-20 | BOLD:ABW6681 | MIB:ZPL:08530 | MT298629 | Anisoptera | Anisoptera | *Somatochlora metallica* | _ | Piedmont | Vercelli | 45.5 | 8.3 | NW | A160 | this study |
| ZPLOD731-20 | BOLD:ABW6681 | MIB:ZPL:08531 | MT298632 | Anisoptera | Anisoptera | *Somatochlora metallica* | M | Trentino-Alto Adige | Trento | 45.8 | 10.5 | NE | A161 | this study |
| ZPLOD733-20 | BOLD:ADC2840 | MIB:ZPL:08533 | MT298633 | Anisoptera | Gomphidae | *Stylurus flavipes* | M | Lombardy | Pavia | 45.3 | 9 | NW | A162 | this study |
| ZPLOD735-20 | BOLD:AAK1032 | MIB:ZPL:08535 | MT298639 | Zygoptera | Lestidae | *Sympecma fusca* | F | Sardinia | Sud Sardegna | 39.8 | 8.9 | SA | Z132 | this study |
| ZPLOD737-20 | BOLD:AAK1032 | MIB:ZPL:08537 | MT298640 | Zygoptera | Lestidae | *Sympecma fusca* | M | Apulia | Bari | 40.9 | 16.6 | S | Z133 | this study |
| ZPLOD739-20 | BOLD:AAK1032 | MIB:ZPL:08539 | MT298635 | Zygoptera | Lestidae | *Sympecma fusca* | F | Sicily | Messina | 38 | 14.7 | SI | Z132 | this study |
| ZPLOD740-20 | BOLD:AAK1032 | MIB:ZPL:08540 | MT298636 | Zygoptera | Lestidae | *Sympecma fusca* | F | Lombardy | Varese | 45.6 | 9 | NW | Z132 | this study |
| ZPLOD741-20 | BOLD:AAK1032 | MIB:ZPL:08541 | MT298637 | Zygoptera | Lestidae | *Sympecma fusca* | F | Piedmont | Biella | 45.5 | 8.1 | NW | Z134 | this study |
| ZPLOD742-20 | BOLD:AAK1032 | MIB:ZPL:08542 | MT298638 | Zygoptera | Lestidae | *Sympecma fusca* | M | Emilia-Romagna | Modena | 44.8 | 10.9 | NW | Z132 | this study |
| ZPLOD744-20 | BOLD:AAK1032 | MIB:ZPL:08544 | MT298634 | Zygoptera | Lestidae | *Sympecma fusca* | M | Sicily | Trapani | 37.9 | 12.7 | SI | Z132 | this study |
| ZPLOD745-20 | BOLD:ACG0335 | MIB:ZPL:08545 | MT298641 | Zygoptera | Lestidae | *Sympecma paedisca* | M | Piedmont | Biella | 45.5 | 8.1 | NW | Z135 | this study |
| ZPLOD746-20 | BOLD:AAA3766 | MIB:ZPL:08546 | MT298642 | Anisoptera | Libellulidae | *Sympetrum danae* | F | Aosta Valley | Aosta | 45.9 | 7.6 | NW | A163 | this study |
| ZPLOD748-20 | BOLD:AAA3766 | MIB:ZPL:08548 | MT298643 | Anisoptera | Libellulidae | *Sympetrum danae* | F | Trentino-Alto Adige | Trento | 46.3 | 11.3 | NE | A164 | this study |
| ZPLOD749-20 | BOLD:ACQ1493 | MIB:ZPL:08549 | MT298645 | Anisoptera | Libellulidae | *Sympetrum depressiusculum* | M | Piedmont | Torino | 45.1 | 7.2 | NW | A165 | this study |
| ZPLOD752-20 | BOLD:ACQ1493 | MIB:ZPL:08552 | MT298644 | Anisoptera | Libellulidae | *Sympetrum depressiusculum* | F | Lombardy | Bergamo | 45.7 | 9.4 | NW | A165 | this study |
| ZPLOD753-20 | BOLD:ADI4820 | MIB:ZPL:08553 | MT298649 | Anisoptera | Libellulidae | *Sympetrum flaveolum* | M | Aosta Valley | Aosta | 45.8 | 7.6 | NW | A166 | this study |
| ZPLOD757-20 | BOLD:ADI4820 | MIB:ZPL:08557 | MT298646 | Anisoptera | Libellulidae | *Sympetrum flaveolum* | M | Emilia-Romagna | Piacenza | 44.6 | 9.5 | NW | A166 | this study |
| ZPLOD758-20 | BOLD:ADI4820 | MIB:ZPL:08558 | MT298647 | Anisoptera | Libellulidae | *Sympetrum flaveolum* | M | Abruzzi | L`Aquila | 42.4 | 13.7 | C | A168 | this study |
| ZPLOD760-20 | BOLD:AAI0218 | MIB:ZPL:08560 | MT298653 | Anisoptera | Libellulidae | *Sympetrum fonscolombii* | M | Sardinia | Sassari | 40.9 | 8.2 | SA | A169 | this study |
| ZPLOD762-20 | BOLD:AAI0218 | MIB:ZPL:08562 | MT298652 | Anisoptera | Libellulidae | *Sympetrum fonscolombii* | M | Apulia | Taranto | 40.4 | 16.9 | S | A170 | this study |
| ZPLOD763-20 | BOLD:AAI0218 | MIB:ZPL:08563 | MT298650 | Anisoptera | Libellulidae | *Sympetrum fonscolombii* | M | Trentino-Alto Adige | Trento | 46 | 11.3 | NE | A170 | this study |
| ZPLOD768-20 | BOLD:AAI0218 | MIB:ZPL:08568 | MT298651 | Anisoptera | Libellulidae | *Sympetrum fonscolombii* | F | Lombardy | Pavia | 45.3 | 9 | NW | A169 | this study |
| ZPLOD772-20 | BOLD:AAD8296 | MIB:ZPL:08572 | MT298655 | Anisoptera | Libellulidae | *Sympetrum meridionale* | M | Sardinia | Sud Sardegna | 39.2 | 8.2 | SA | A172 | this study |
| ZPLOD776-20 | BOLD:AAD8296 | MIB:ZPL:08576 | MT298654 | Anisoptera | Libellulidae | *Sympetrum meridionale* | F | Apulia | Bari | 40.9 | 17.1 | S | A173 | this study |
| ZPLOD777-20 | BOLD:AAD8296 | MIB:ZPL:08577 | MT298657 | Anisoptera | Libellulidae | *Sympetrum meridionale* | M | Emilia-Romagna | Modena | 44.5 | 10.9 | NW | A174 | this study |
| ZPLOD779-20 | BOLD:AAK1022 | MIB:ZPL:08579 | MT298659 | Anisoptera | Libellulidae | *Sympetrum pedemontanum* | _ | Piedmont | Alessandria | 45.2 | 8.3 | NW | A175 | this study |
| ZPLOD781-20 | BOLD:AAK1022 | MIB:ZPL:08581 | MT298658 | Anisoptera | Libellulidae | *Sympetrum pedemontanum* | M | Lombardy | Pavia | 45.3 | 9 | NW | A176 | this study |
| ZPLOD783-20 | BOLD:AAB2237 | MIB:ZPL:08583 | MT298663 | Anisoptera | Libellulidae | *Sympetrum sanguineum* | M | Sicily | Messina | 37.9 | 14.7 | SI | A177 | this study |
| ZPLOD786-20 | BOLD:AAB2237 | MIB:ZPL:08586 | MT298661 | Anisoptera | Libellulidae | *Sympetrum sanguineum* | M | Trentino-Alto Adige | Trento | 46.1 | 11.2 | NE | A178 | this study |
| ZPLOD790-20 | BOLD:AAB2237 | MIB:ZPL:08590 | MT298662 | Anisoptera | Libellulidae | *Sympetrum sanguineum* | F | Lombardy | Lecco | 45.8 | 9.3 | NW | A179 | this study |
| ZPLOD793-20 | BOLD:AAB2237 | MIB:ZPL:08593 | MT298660 | Anisoptera | Libellulidae | *Sympetrum sanguineum* | M | Molise | Campobasso | 41.8 | 15 | S | A178 | this study |
| ZPLOD794-20 | BOLD:AAB2237 | MIB:ZPL:08594 | MT298664 | Anisoptera | Libellulidae | *Sympetrum sanguineum* | M | Abruzzi | L`Aquila | 42.3 | 13.6 | C | A179 | this study |
| ZPLOD795-20 | BOLD:AAB2236 | MIB:ZPL:08595 | MT298667 | Anisoptera | Libellulidae | *Sympetrum striolatum* | M | Sardinia | Sassari | 40.8 | 8.5 | SA | A180 | this study |
| ZPLOD797-20 | BOLD:AAB2236 | MIB:ZPL:08597 | MT298666 | Anisoptera | Libellulidae | *Sympetrum striolatum* | M | Trentino-Alto Adige | Trento | 46 | 11 | NE | A180 | this study |
| ZPLOD799-20 | BOLD:AAB2236 | MIB:ZPL:08599 | MT298669 | Anisoptera | Libellulidae | *Sympetrum striolatum* | M | Apulia | Bari | 41.1 | 16.8 | S | A180 | this study |
| ZPLOD800-20 | BOLD:AAB2236 | MIB:ZPL:08600 | MT298668 | Anisoptera | Libellulidae | *Sympetrum striolatum* | F | Emilia-Romagna | Piacenza | 44.6 | 9.5 | NW | A182 | this study |
| ZPLOD802-20 | BOLD:AAE2658 | MIB:ZPL:08602 | MT298671 | Anisoptera | Libellulidae | *Sympetrum vulgatum* | M | Trentino-Alto Adige | Trento | 46.3 | 11.4 | NE | A183 | this study |
| ZPLOD803-20 | BOLD:AAE2658 | MIB:ZPL:08603 | MT298672 | Anisoptera | Libellulidae | *Sympetrum vulgatum* | M | Lombardy | Lecco | 45.8 | 9.4 | NW | A183 | this study |
| ZPLOD804-20 | BOLD:ABA9471 | MIB:ZPL:08604 | MT298676 | Anisoptera | Libellulidae | *Trithemis annulata* | M | Sardinia | Sassari | 40.7 | 8.2 | SA | A184 | this study |
| ZPLOD805-20 | BOLD:ABA9471 | MIB:ZPL:08605 | MT298675 | Anisoptera | Libellulidae | *Trithemis annulata* | M | Liguria | Spezia | 44.1 | 10 | NW | A185 | this study |
| ZPLOD809-20 | BOLD:ABA9471 | MIB:ZPL:08609 | MT298673 | Anisoptera | Libellulidae | *Trithemis annulata* | M | Lombardy | Milano | 45.4 | 9.1 | NW | A185 | this study |
| ZPLOD811-20 | BOLD:ABA9471 | MIB:ZPL:08611 | MT298674 | Anisoptera | Libellulidae | *Trithemis annulata* | M | Sicily | Trapani | 37.7 | 12.8 | SI | A186 | this study |
| ZPLOD812-20 | BOLD:AAZ4163 | MIB:ZPL:08612 | MT298679 | Anisoptera | Libellulidae | *Zygonyx torridus* | M | Sicily | Trapani | 37.7 | 12.9 | SI | A187 | this study |

MOLECULAR ECOLOGY
RESOURCES

**APPENDIX S2**: dataset DS2, composed by the sequences mined from BOLD and NCBI Genbank plus dataset DS1 and other COI barcode sequences obtained from additional samples retrieved by authors in other Western Palearctic countries. Species taxonomy, DS1 specimen IDs, depositories accessions, are reported for each COI haplotype.

| Suborder | Species name/Sample ID | Haplotype | DNA barcode source |
|---|---|---|---|
| Anisoptera | Aeshna_affinis_ASG673_SS | A1 | DS1 |
| Anisoptera | Aeshna_affinis_GA54_TO_Aeshna_affinis_HM422047 | A2 | DS1 / NCBI |
| Anisoptera | Aeshna_affinis_KJ873232 | A188 | NCBI |
| Anisoptera | Aeshna_affinis_MIBAG0111_TA | A3 | DS1 |
| Anisoptera | Aeshna_caerulea_GA76_TN | A4 | DS1 |
| Anisoptera | Aeshna_cyanea_GBMIX2219_15_Aeshna_cyanea_MIBAG0092_BG | A6 | DS1 / BOLD |
| Anisoptera | Aeshna_cyanea_KC912199_Aeshna_cyanea_KC912200_Aeshna_cyanea_KC912201_Aeshna_cyanea_KC912202_Aeshna_cyanea_KY847574_Aeshna_cyanea_KY847575_Aeshna_cyanea_KY847576_Aeshna_cyanea_KY847577 | A189 | NCBI |
| Anisoptera | Aeshna_cyanea_KU180304_Aeshna_cyanea_KU180306 | A190 | NCBI |
| Anisoptera | Aeshna_cyanea_KU180305_Sicily | A5 | DS1 / NCBI |
| Anisoptera | Aeshna_cyanea_KU180307 | A191 | NCBI |
| Anisoptera | Aeshna_cyanea_KU180308 | A192 | NCBI |
| Anisoptera | Aeshna_cyanea_KU180309 | A193 | NCBI |
| Anisoptera | Aeshna_cyanea_KU180310_Aeshna_cyanea_KU180311 | A194 | NCBI |
| Anisoptera | Aeshna_cyanea_KU180312_Aeshna_cyanea_KU180313_Aeshna_cyanea_KU180314_Aeshna_cyanea_KU180315_Aeshna_cyanea_KU180316_Aeshna_cyanea_KU180317_Aeshna_cyanea_KU180319_Aeshna_cyanea_KU180321 | A195 | NCBI |
| Anisoptera | Aeshna_cyanea_KU180318 | A196 | NCBI |
| Anisoptera | Aeshna_cyanea_KU180320 | A197 | NCBI |
| Anisoptera | Aeshna_grandis_GA79_TN_Aeshna_grandis_GA79_TN | A7 | DS1 |
| Anisoptera | Aeshna_grandis_KC912203_Aeshna_grandis_KY847578 | A198 | NCBI |
| Anisoptera | Aeshna_grandis_KJ873213 | A199 | NCBI |
| Anisoptera | Aeshna_grandis_KU180299 | A200 | NCBI |
| Anisoptera | Aeshna_grandis_KY926549_Aeshna_grandis_KY926550_Aeshna_grandis_KY926551_Aeshna_grandis_KY926552_Aeshna_grandis_KY926553_Aeshna_grandis_KY926554_Aeshna_grandis_KY926555_Aeshna_grandis_KY926556_Aeshna_grandis_KY926557_Aeshna_grandis_KY926558_Aeshna_grandis_KY926559_Aeshna_grandis_KY926560_Aeshna_grandis_KY926561_Aeshna_grandis_KY926562_Aeshna_grandis_KY926563_Aeshna_grandis_KY926564_Aeshna_grandis_KY926565_Aeshna_grandis_KY926582_Aeshna_grandis_KY926584_Aeshna_grandis_ZMBN005_15 | A202 | NCBI |
| Anisoptera | Aeshna_grandis_KY926566 | A203 | NCBI |
| Anisoptera | Aeshna_grandis_KY926567 | A204 | NCBI |
| Anisoptera | Aeshna_grandis_KY926579 | A205 | NCBI |
| Anisoptera | Aeshna_grandis_KY926585_Aeshna_grandis_KY926586_Aeshna_grandis_KY926587_Aeshna_grandis_KY926588_Aeshna_grandis_MIBAG0168_AO | A8 | DS1 / NCBI |
| Anisoptera | Aeshna_grandis_KY926589 | A206 | NCBI |
| Anisoptera | Aeshna_grandis_KY926591 | A207 | NCBI |
| Anisoptera | Aeshna_grandis_KY926592 | A208 | NCBI |
| Anisoptera | Aeshna_grandis_KY926593 | A209 | NCBI |
| Anisoptera | Aeshna_grandis_KY926594 | A210 | NCBI |
| Anisoptera | Aeshna_grandis_KY926595 | A211 | NCBI |
| Anisoptera | Aeshna_grandis_KY926597 | A212 | NCBI |
| Anisoptera | Aeshna_grandis_KY926598 | A213 | NCBI |
| Anisoptera | Aeshna_grandis_KY926599 | A214 | NCBI |
| Anisoptera | Aeshna_grandis_KY926600 | A215 | NCBI |
| Anisoptera | Aeshna_isoceles_ASG672_SS | A9 | DS1 |
| Anisoptera | Aeshna_isoceles_MG11_BR | A10 | DS1 |
| Anisoptera | Aeshna_isoceles_MIBAG0021_CO | A11 | DS1 |
| Anisoptera | Aeshna_juncea_DB14_AO | A12 | DS1 |
| Anisoptera | Aeshna_juncea_FBAQU1431_13 | A217 | BOLD |
| Anisoptera | Aeshna_juncea_GA72_TN | A13 | DS1 |
| Anisoptera | Aeshna_juncea_JF839254 | A219 | NCBI |
| Anisoptera | Aeshna_juncea_JF839255_Aeshna_juncea_JF839256_Aeshna_juncea_KR144886_Aeshna_juncea_KR148626 | A220 | NCBI |
| Anisoptera | Aeshna_juncea_JF839257_Aeshna_juncea_KR148296 | A221 | NCBI |
| Anisoptera | Aeshna_juncea_JF839297 | A222 | NCBI |
| Anisoptera | Aeshna_juncea_JN294385_Aeshna_juncea_JN294387 | A223 | NCBI |
| Anisoptera | Aeshna_juncea_JN294388_Aeshna_juncea_KR142592_Aeshna_juncea_KR143717_Aeshna_juncea_KR147641 | A224 | NCBI |
| Anisoptera | Aeshna_juncea_KF369278 | A225 | NCBI |
| Anisoptera | Aeshna_juncea_KR140582 | A226 | NCBI |
| Anisoptera | Aeshna_juncea_KR141108 | A227 | NCBI |
| Anisoptera | Aeshna_juncea_KR142266 | A228 | NCBI |
| Anisoptera | Aeshna_juncea_KR142375 | A229 | NCBI |
| Anisoptera | Aeshna_juncea_KR142385 | A230 | NCBI |
| Anisoptera | Aeshna_juncea_KR142479 | A231 | NCBI |
| Anisoptera | Aeshna_juncea_KR142506 | A232 | NCBI |
| Anisoptera | Aeshna_juncea_KR142917 | A233 | NCBI |
| Anisoptera | Aeshna_juncea_KR143341 | A234 | NCBI |
| Anisoptera | Aeshna_juncea_KR144353 | A235 | NCBI |
| Anisoptera | Aeshna_juncea_KR144661 | A236 | NCBI |
| Anisoptera | Aeshna_juncea_KR144938 | A237 | NCBI |
| Anisoptera | Aeshna_juncea_KR145533 | A238 | NCBI |
| Anisoptera | Aeshna_juncea_KR146790 | A239 | NCBI |
| Anisoptera | Aeshna_juncea_KR147910 | A240 | NCBI |
| Anisoptera | Aeshna_juncea_KU180297 | A241 | NCBI |

| Suborder | Species name/Sample ID | Haplotype | DNA barcode source |
|---|---|---|---|
| Anisoptera | Aeshna_juncea_KU873989_Aeshna_juncea_KU873990 | A242 | NCBI |
| Anisoptera | Aeshna_juncea_MF358804 | A243 | NCBI |
| Anisoptera | Aeshna_juncea_MF358805 | A244 | NCBI |
| Anisoptera | Aeshna_juncea_MIBAG0222_SO | A14 | DS1 |
| Anisoptera | Aeshna_juncea_ZPLOD813-20 | A218 | BOLD |
| Anisoptera | Aeshna_mixta_FBAQU480_10 | A245 | BOLD |
| Anisoptera | Aeshna_mixta_FL46_MC | A15 | DS1 |
| Anisoptera | Aeshna_mixta_HM901884 | A247 | BOLD |
| Anisoptera | Aeshna_mixta_KC912204_Aeshna_mixta_KY847569 | A248 | NCBI |
| Anisoptera | Aeshna_mixta_KC912205_Aeshna_mixta_KY847570 | A249 | NCBI |
| Anisoptera | Aeshna_mixta_MIBAG0170_CO | A16 | DS1 |
| Anisoptera | Aeshna_mixta_ZPLOD814-20 | A246 | BOLD |
| Anisoptera | Aeshna_subarctica_GA71_TN | A17 | DS1 |
| Anisoptera | Aeshna_subarctica_GA80_TN | A18 | DS1 |
| Anisoptera | Aeshna_subarctica_HM901883 | A250 | NCBI |
| Anisoptera | Aeshna_subarctica_JF839363 | A251 | NCBI |
| Anisoptera | Aeshna_subarctica_JN294386 | A252 | NCBI |
| Anisoptera | Aeshna_subarctica_KR143264 | A253 | NCBI |
| Anisoptera | Aeshna_subarctica_KR147009 | A254 | NCBI |
| Anisoptera | Aeshna_subarctica_KU180298 | A255 | NCBI |
| Anisoptera | Anax_ephippiger_DB09_GE | A19 | DS1 |
| Anisoptera | Anax_ephippiger_KC912208_Anax_ephippiger_KC912209 | A256 | NCBI |
| Anisoptera | Anax_ephippiger_KC912210_Anax_ephippiger_KC912213_Anax_ephippiger_KC912214 | A257 | NCBI |
| Anisoptera | Anax_ephippiger_KC912211 | A258 | NCBI |
| Anisoptera | Anax_ephippiger_KC912212 | A259 | NCBI |
| Anisoptera | Anax_ephippiger_KC912215 | A260 | NCBI |
| Anisoptera | Anax_ephippiger_KC912216 | A261 | NCBI |
| Anisoptera | Anax_ephippiger_KC912217 | A262 | NCBI |
| Anisoptera | Anax_ephippiger_MIBAG0096_PV | A20 | DS1 |
| Anisoptera | Anax_imperator_ASG682_SS_Anax_imperator_FL07_FM_Anax_imperator_GA67_TO_Anax_imperator_MIBAG0068_MI | A21 | DS1 |
| Anisoptera | Anax_imperator_GA26_TN | A22 | DS1 |
| Anisoptera | Anax_imperator_HM901859 | A263 | NCBI |
| Anisoptera | Anax_imperator_KC912218_Anax_imperator_KC912221_Anax_imperator_KC912222_Anax_imperator_KC912223_Anax_imperator_KC912224_Anax_imperator_KC912225_ Anax_imperator_KC912226_Anax_imperator_KY847559_Anax_imperator_KY847560_Anax_imperator_KY847561_Anax_imperator_KY847562_Anax_imperator_KY847563_ Anax_imperator_KY847566_Anax_imperator_KY847568 | A264 | NCBI |
| Anisoptera | Anax_imperator_KC912219_Anax_imperator_KC912220_Anax_imperator_KY847558_Anax_imperator_KY847564 | A265 | NCBI |
| Anisoptera | Anax_imperator_KC912227_Anax_imperator_KY847567 | A266 | NCBI |
| Anisoptera | Anax_imperator_KC912228_Anax_imperator_KY847565 | A267 | NCBI |
| Anisoptera | Anax_imperator_KF584974 | A268 | NCBI |
| Anisoptera | Anax_imperator_KU565916 | A269 | NCBI |
| Anisoptera | Anax_imperator_KX161841_Anax_imperator_NC_031821 | A270 | NCBI |
| Anisoptera | Anax_imperator_LI12_ME | A23 | DS1 |
| Anisoptera | Anax_imperator_MIBAG0266_CB | A24 | DS1 |
| Anisoptera | Anax_imperator_SS08_TP_Anax_parthenope_FL37_MC | A25 | DS1 |
| Anisoptera | Anax_parthenope_KC135891 | A271 | NCBI |
| Anisoptera | Anax_parthenope_KF257072 | A272 | NCBI |
| Anisoptera | Anax_parthenope_KR149805 | A273 | NCBI |
| Anisoptera | Anax_parthenope_MH669065 | A274 | NCBI |
| Anisoptera | Anax_parthenope_MIBAG0106_VA | A27 | DS1 |
| Anisoptera | Anax_parthenope_MIBAG0240_BR | A28 | DS1 |
| Anisoptera | Boyeria_irene_ASG732_SU | A29 | DS1 |
| Anisoptera | Boyeria_irene_GA60_TO_Boyeria_irene_LI01_PV | A30 | DS1 |
| Anisoptera | Brachytemis_impartita_LI27_CT_Brachythemis_impartita_ASG468_SS_Brachythemis_impartita_ASG725_OR | A31 | DS1 |
| Anisoptera | Brachythemis_impartita_ZPLOD842-20 | A275 | BOLD |
| Anisoptera | Brachytron_pratense_GLP06 | A33 | DS1 |
| Anisoptera | Brachytron_pratense_IC01_UD_Brachytron_pratense_MG16_LE | A34 | DS1 |
| Anisoptera | Brachytron_pratense_KC912235_Brachytron_pratense_KC912236_Brachytron_pratense_KY847550_Brachytron_pratense_KY847551 | A276 | NCBI |
| Anisoptera | Brachytron_pratense_MIBAG0184_NO | A35 | DS1 |
| Zygoptera | Calopteryx_haemorrhoidalis_ASG694_SS_Calopteryx_haemorrhoidalis_FM23_BT_Calopteryx_haemorrhoidalis_LI13_ME_Calopteryx_haemorrhoidalis_MIBAG0180_GR_Calopteryx_haemorrhoidalis_SR11_LT | Z1 | DS1 |
| Zygoptera | Calopteryx_haemorrhoidalis_DB04_GE | Z2 | DS1 |
| Zygoptera | Calopteryx_haemorrhoidalis_DB05_GE_Calopteryx_haemorrhoidalis_DB06_GE | Z3 | DS1 |
| Zygoptera | Calopteryx_splendens_FM16_BA | Z5 | DS1 |
| Zygoptera | Calopteryx_splendens_GA13_TN_Calopteryx_splendens_MIBAG0009_CO_Calopteryx_splendens_GAM13_MON_2_Calopteryx_xanthostoma_DB28_SV_Calopteryx_xanthostoma_DB29_SV | Z6 | DS1 |
| Zygoptera | Calopteryx_splendens_SR10_LT | Z8 | DS1 |
| Zygoptera | Calopteryx_splendens_ZPLOD816-20_Calopteryx_xanthostoma_DB29_SV | Z13 | DS1 / BOLD |
| Zygoptera | Calopteryx_virgo_DB10_CN_Calopteryx_virgo_GA28_TN_Calopteryx_virgo_GODO00118_Calopteryx_virgo_MIBAG0087_PV | Z9 | DS1 |
| Zygoptera | Calopteryx_virgo_FL19_MC_Calopteryx_virgo_GLP11_PG | Z10 | DS1 |
| Zygoptera | Calopteryx_virgo_HM901860 | Z137 | NCBI |
| Zygoptera | Calopteryx_virgo_HM901885 | Z138 | NCBI |
| Zygoptera | Calopteryx_virgo_ZPLOD817-20 | Z136 | BOLD |
| Zygoptera | Ceriagrion_tenellum_ASG702_SS_Ceriagrion_tenellum_MG15_LE_Ceriagrion_tenellum_SR15_LT_Ceriagrion_tenellum_LI15_SR | Z14 | DS1 |
| Zygoptera | Ceriagrion_tenellum_IC18_UD_Ceriagrion_tenellum_MIBAG0024_CO_Ceriagrion_tenellum_MIBAG0230_CN | Z15 | DS1 |
| Zygoptera | Ceriagrion_tenellum_KC912305_Ceriagrion_tenellum_KC912306_Ceriagrion_tenellum_KC912309 | Z139 | NCBI |
| Zygoptera | Ceriagrion_tenellum_KC912307_Ceriagrion_tenellum_KC912308 | Z140 | NCBI |
| Zygoptera | Chalcolestes_parvidens_GLP13_TR_Chalcolestes_parvidens_IC34_GO_Chalcolestes_parvidens_MIBAG0300_BO_Chalcolestes_parvidens_IC33_TS | Z17 | DS1 |
| Zygoptera | Chalcolestes_viridis_ASG735_SU_Chalcolestes_viridis_GLP14_PG_Chalcolestes_viridis_IC30_UD_Chalcolestes_viridis_LI21_CT_Chalcolestes_viridis_SS24_TP | Z19 | DS1 |

| Suborder | Species name/Sample ID | Haplotype | DNA barcode source |
|---|---|---|---|
| Zygoptera | Chalcolestes_viridis_ASG760_NU | Z20 | DS1 |
| Zygoptera | Chalcolestes_viridis_FM39_BA | Z21 | DS1 |
| Zygoptera | Chalcolestes_viridis_FM40_BA | Z22 | DS1 |
| Zygoptera | Chalcolestes_viridis_GA78_TN | Z23 | DS1 |
| Zygoptera | Chalcolestes_viridis_GU682188 | Z141 | NCBI |
| Zygoptera | Chalcolestes_viridis_GU682190 | Z142 | NCBI |
| Zygoptera | Chalcolestes_viridis_MIBAG0108_CO | Z24 | DS1 |
| Zygoptera | Coenagrion_caerulescens_AC08_AG | Z27 | DS1 |
| Zygoptera | Coenagrion_caerulescens_ASG700_SS | Z28 | DS1 |
| Zygoptera | Coenagrion_caerulescens_KP272398_Df1005_MAR | Z143 | NCBI |
| Zygoptera | Coenagrion_caerulescens_KP272422 | Z29 | NCBI |
| Zygoptera | Coenagrion_caerulescens_KP272528_Df1669_MAR | Z144 | NCBI |
| Zygoptera | Coenagrion_caerulescens_KP272542_Df1713_TUN | Z145 | NCBI |
| Zygoptera | Coenagrion_caerulescens_KP272593_Df972_MAR | Z146 | NCBI |
| Zygoptera | Coenagrion_caerulescens_MIBAG0231_CN | Z30 | DS1 |
| Zygoptera | Coenagrion_caerulescens_MIBAG0232_CN | Z31 | DS1 |
| Zygoptera | Coenagrion_caerulescens_MIBAG0233_CN_Coenagrion_caerulescens_MIBAG0234_CN | Z32 | DS1 |
| Zygoptera | Coenagrion_caerulescens_MIBAG0267_CB_Coenagrion_caerulescens_FM12_BA | Z26 | DS1 |
| Zygoptera | Coenagrion_hastulatum_GA18_TN_Coenagrion_hastulatum_FBAQU556_10_Coenagrion_hastulatum_GAM29_MON | Z34 | DS1 |
| Zygoptera | Coenagrion_hastulatum_IC14_UD | Z35 | DS1 |
| Zygoptera | Coenagrion_hastulatum_ZPLOD818-20 | Z148 | BOLD |
| Zygoptera | Coenagrion_mercuriale_FBAQU1429_13 | Z149 | BOLD |
| Zygoptera | Coenagrion_mercuriale_FL01_MC_Coenagrion_mercuriale_FM01_LE_Coenagrion_mercuriale_FM26_BT_Coenagrion_mercuriale_KX241512_Rome_Coenagrion_mercuriale_KX241513_Rome_Coenagrion_mercuriale_MIBAG0226_CN_Coenagrion_mercuriale_MIBAG0228_CN | Z36 | DS1 / NCBI |
| Zygoptera | Coenagrion_mercuriale_GLP15_PG_Coenagrion_mercuriale_SR02_LT | Z37 | DS1 |
| Zygoptera | Coenagrion_mercuriale_KP272399_Df1018_ESP | Z150 | NCBI |
| Zygoptera | Coenagrion_mercuriale_KP272400_Df1020_ESP_Coenagrion_mercuriale_KP272402_Df1022_ESP_Coenagrion_mercuriale_KP272428_Df1258_ESP_Coenagrion_mercuriale_KP272429_Df1259_ESP_Coenagrion_mercuriale_KP272430_Df1260_ESP_Coenagrion_mercuriale_KP272437_Df1268_ESP_Coenagrion_mercuriale_KP272443_Df1285_ESP_Coenagrion_mercuriale_KP272445_Df1289_ESP_Coenagrion_mercuriale_KP272446_Df1290_ESP_Coenagrion_mercuriale_KP272487_Df1524_ESP_Coenagrion_mercuriale_KP272500_Df1561_ESP_Coenagrion_mercuriale_KP272517_Df1591_PRT_Coenagrion_mercuriale_KP272520_Df1595_PRT_Coenagrion_mercuriale_KP272521_Df1616_PRT_Coenagrion_mercuriale_KP272522_Df1617_PRT_Coenagrion_mercuriale_KP272523_Df1650_PRT_Coenagrion_mercuriale_KP272524_Df1651_PRT_Coenagrion_mercuriale_KP272561_Df273_PRT_Coenagrion_mercuriale_KP272565_Df508_PRT_Coenagrion_mercuriale_KP272567_Df543_PRT_Coenagrion_mercuriale_KP272568_Df544_PRT_Coenagrion_mercuriale_KP272569_Df554_PRT_Coenagrion_mercuriale_KP272570_Df555_PRT_Coenagrion_mercuriale_KP272571_Df584_ESP_Coenagrion_mercuriale_KP272572_Df585_ESP_Coenagrion_mercuriale_KP272573_Df591_ESP_Coenagrion_mercuriale_KP272574_Df593_ESP | Z151 | NCBI |
| Zygoptera | Coenagrion_mercuriale_KP272401_Df1021_ESP_Coenagrion_mercuriale_KP272434_Df1265_ESP_Coenagrion_mercuriale_KP272435_Df1266_ESP_Coenagrion_mercuriale_KP272436_Df1267_ESP_Coenagrion_mercuriale_KP272497_Df1543_ESP_Coenagrion_mercuriale_KP272498_Df1544_ESP | Z152 | NCBI |
| Zygoptera | Coenagrion_mercuriale_KP272404_Df1155_DZA_Coenagrion_mercuriale_KP272405_Df1159_DZA_Coenagrion_mercuriale_KP272406_Df1161_DZA_Coenagrion_mercuriale_KP272407_Df1162_DZA_Coenagrion_mercuriale_KP272408_Df1163_DZA_Coenagrion_mercuriale_KP272409_Df1164_DZA_Coenagrion_mercuriale_KP272410_Df1165_DZA_Coenagrion_mercuriale_KP272411_Df1166_DZA_Coenagrion_mercuriale_KP272412_Df1170_DZA_Coenagrion_mercuriale_KP272413_Df1171_DZA_Coenagrion_mercuriale_KP272512_Df1581_TUN_Coenagrion_mercuriale_KP272515_Df1584_TUN_Coenagrion_mercuriale_KP272516_Df1585_TUN_Coenagrion_mercuriale_KP272525_Df1653_MAR_Coenagrion_mercuriale_KP272529_Df1673_MAR_Coenagrion_mercuriale_KP272530_Df1674_MAR_Coenagrion_mercuriale_KP272531_Df1685_MAR_Coenagrion_mercuriale_KP272532_Df1687_MAR_Coenagrion_mercuriale_KP272533_Df1688_MAR_Coenagrion_mercuriale_KP272562_Df283_MAR_Coenagrion_mercuriale_KP272563_Df284_MAR_Coenagrion_mercuriale_KP272564_Df288_MAR_Coenagrion_mercuriale_KP272585_Df836_MAR_Coenagrion_mercuriale_KP272587_Df840_MAR_Coenagrion_mercuriale_KP272590_Df951_MAR_Coenagrion_mercuriale_KP272591_Df954_MAR | Z153 | NCBI |
| Zygoptera | Coenagrion_mercuriale_KP272414_Coenagrion_mercuriale_KP272415_Coenagrion_mercuriale_KP272416_Coenagrion_mercuriale_KP272417_Coenagrion_mercuriale_KP272418_Coenagrion_mercuriale_KP272419_Coenagrion_mercuriale_KP272420_Coenagrion_mercuriale_KP272424_Coenagrion_mercuriale_KP272425_Coenagrion_mercuriale_KP272505_Coenagrion_mercuriale_KP272506_Coenagrion_mercuriale_KP272507_Coenagrion_mercuriale_KP272551_Coenagrion_mercuriale_KP272552_Coenagrion_mercuriale_KP272553_Coenagrion_mercuriale_KP272555_Coenagrion_mercuriale_KP272556_Coenagrion_mercuriale_KP272557_Coenagrion_mercuriale_KP272558_Coenagrion_mercuriale_KP272559_RA_Coenagrion_mercuriale_KP272560 | Z38 | NCBI |
| Zygoptera | Coenagrion_mercuriale_KP272426_Df1255_ESP | Z154 | NCBI |
| Zygoptera | Coenagrion_mercuriale_KP272427_Df1257_ESP_Coenagrion_mercuriale_KP272440_Df1272_ESP | Z155 | NCBI |
| Zygoptera | Coenagrion_mercuriale_KP272431_Df1261_ESP | Z156 | NCBI |
| Zygoptera | Coenagrion_mercuriale_KP272432_Df1262_ESP | Z157 | NCBI |
| Zygoptera | Coenagrion_mercuriale_KP272433_Df1263_ESP | Z158 | NCBI |
| Zygoptera | Coenagrion_mercuriale_KP272438_Df1269_ESP_Coenagrion_mercuriale_KP272444_Df1286_ESP | Z159 | NCBI |
| Zygoptera | Coenagrion_mercuriale_KP272439_Df1270_ESP | Z160 | NCBI |
| Zygoptera | Coenagrion_mercuriale_KP272441_Df1274_ESP | Z161 | NCBI |
| Zygoptera | Coenagrion_mercuriale_KP272442_Df1278_ESP | Z162 | NCBI |
| Zygoptera | Coenagrion_mercuriale_KP272447_Df1292_ESP | Z163 | NCBI |
| Zygoptera | Coenagrion_mercuriale_KP272448_Coenagrion_mercuriale_KP272449_Coenagrion_mercuriale_KP272456_Coenagrion_mercuriale_KP272457 | Z39 | NCBI |
| Zygoptera | Coenagrion_mercuriale_KP272450 | Z40 | NCBI |
| Zygoptera | Coenagrion_mercuriale_KP272451_Df1303_FRA_Coenagrion_mercuriale_KP272452_Df1306_FRA_Coenagrion_mercuriale_KP272453_Df1307_GBR_Coenagrion_mercuriale_KP272454_Df1308_GBR_Coenagrion_mercuriale_KP272455_Df1311_GBR_Coenagrion_mercuriale_KP272458_Df1336_DEU_Coenagrion_mercuriale_KP272459_Df1347_DEU_Coenagrion_mercuriale_KP272460_Df1348_DEU_Coenagrion_mercuriale_KP272461_Df1350_DEU_Coenagrion_mercuriale_KP272462_Df1351_DEU_Coenagrion_mercuriale_KP272465_Df1360_FRA_Coenagrion_mercuriale_KP272467_Df1364_FRA_Coenagrion_mercuriale_KP272468_Df1365_FRA_Coenagrion_mercuriale_KP272469_Df1367_FRA_Coenagrion_mercuriale_KP272475_Df1391_GBR_Coenagrion_mercuriale_KP272477_Df1393_GBR_Coenagrion_mercuriale_KP272478_Df1396_GBR_Coenagrion_mercuriale_KP272479_Df1397_GBR_Coenagrion_mercuriale_KP272482_Df1411_DEU_Coenagrion_mercuriale_KP272483_Df1412_DEU_Coenagrion_mercuriale_KP272484_Df1516_GBR_Coenagrion_mercuriale_KP272485_Df1517_GBR_Coenagrion_mercuriale_KP272486_Df1520_GBR_Coenagrion_mercuriale_KP272488_Df1530_GBR_Coenagrion_mercuriale_KP272489_Df1531_GBR_Coenagrion_mercuriale_KP272490_Df1532_GBR_Coenagrion_mercuriale_KP272492_Df1535_DEU_Coenagrion_mercuriale_KP272493_Df1536_DEU_Coenagrion_mercuriale_KP272494_Df1537_DEU | Z164 | NCBI |
| Zygoptera | Coenagrion_mercuriale_KP272463_Df1357_FRA_Coenagrion_mercuriale_KP272471_Df1372_FRA_Coenagrion_mercuriale_KP272480_Df1398_DEU_Coenagrion_mercuriale_KP272481_Df1400_DEU_Coenagrion_mercuriale_KP272491_Df1534_DEU_Coenagrion_mercuriale_KP272495_Df1538_DEU_Coenagrion_mercuriale_KP272496_Df1539_DEU_Coenagrion_mercuriale_KP272534_Df1697_CHE_Coenagrion_mercuriale_KP272535_Df1698_CHE_Coenagrion_mercuriale_KP272536_Df1699_CHE_Coenagrion_mercuriale_KP272537_Df1700_CHE | Z165 | NCBI |
| Zygoptera | Coenagrion_mercuriale_KP272464_Df1358_FRA_Coenagrion_mercuriale_KP272470_Df1369_FRA | Z166 | NCBI |
| Zygoptera | Coenagrion_mercuriale_KP272466_Df1362_FRA | Z167 | NCBI |
| Zygoptera | Coenagrion_mercuriale_KP272472_Df1381_FRA | Z168 | NCBI |
| Zygoptera | Coenagrion_mercuriale_KP272473_Df1382_FRA | Z169 | NCBI |
| Zygoptera | Coenagrion_mercuriale_KP272474_Df1389_GBR | Z170 | NCBI |
| Zygoptera | Coenagrion_mercuriale_KP272476_Df1392_DEU | Z171 | NCBI |

MOLECULAR ECOLOGY RESOURCES

| Suborder | Species name/Sample ID | Haplotype | DNA barcode source |
|---|---|---|---|
| Zygoptera | Coenagrion_mercuriale_KP272499_Df1560_ESP | Z172 | NCBI |
| Zygoptera | Coenagrion_mercuriale_KP272501_Df1563_ESP | Z173 | NCBI |
| Zygoptera | Coenagrion_mercuriale_KP272508_Df1577_TUN_Coenagrion_mercuriale_KP272509_Df1578_TUN_Coenagrion_mercuriale_KP272510_Df1579_TUN_Coenagrion_mercuriale_KP272511_Df1580_TUN_Coenagrion_mercuriale_KP272514_Df1583_TUN | Z174 | NCBI |
| Zygoptera | Coenagrion_mercuriale_KP272513_Df1582_TUN | Z175 | NCBI |
| Zygoptera | Coenagrion_mercuriale_KP272518_Df1593_PRT | Z176 | NCBI |
| Zygoptera | Coenagrion_mercuriale_KP272519_Df1594_PRT | Z177 | NCBI |
| Zygoptera | Coenagrion_mercuriale_KP272526_Df1654_MAR | Z178 | NCBI |
| Zygoptera | Coenagrion_mercuriale_KP272527_Df1667_MAR | Z179 | NCBI |
| Zygoptera | Coenagrion_mercuriale_KP272554 | Z41 | NCBI |
| Zygoptera | Coenagrion_mercuriale_KP272566_Df509_PRT | Z180 | NCBI |
| Zygoptera | Coenagrion_mercuriale_KP272575_Df823_MAR_Coenagrion_mercuriale_KP272576_Df825_MAR_Coenagrion_mercuriale_KP272577_Df826_MAR_Coenagrion_mercuriale_KP272579_Df828_MAR_Coenagrion_mercuriale_KP272580_Df829_MAR_Coenagrion_mercuriale_KP272581_Df830_MAR_Coenagrion_mercuriale_KP272582_Df831_MAR_Coenagrion_mercuriale_KP272583_Df832_MAR_Coenagrion_mercuriale_KP272584_Df833_MAR | Z181 | NCBI |
| Zygoptera | Coenagrion_mercuriale_KP272578_Df827_MAR | Z182 | NCBI |
| Zygoptera | Coenagrion_mercuriale_KP272586_Df837_MAR | Z183 | NCBI |
| Zygoptera | Coenagrion_mercuriale_KP272588_Df912_MAR | Z184 | NCBI |
| Zygoptera | Coenagrion_mercuriale_KP272589_Df913_MAR | Z185 | NCBI |
| Zygoptera | Coenagrion_mercuriale_KX241514_Rome | Z42 | NCBI |
| Zygoptera | Coenagrion_ornatum_FBAQU1446_13_Coenagrion_ornatum_FBAQU1447_13_Coenagrion_ornatum_FBAQU1448_13_Coenagrion_ornatum_FBAQU559_10 | Z186 | BOLD |
| Zygoptera | Coenagrion_ornatum_FM15_BA_Coenagrion_ornatum_FM17_PZ_Coenagrion_ornatum_FM21_BT | Z43 | DS1 |
| Zygoptera | Coenagrion_ornatum_KP272502_Df1568_SVN_Coenagrion_puella_KP272403_Df1143_RUS_Coenagrion_puella_KP272543_Df1716_RUS | Z187 | NCBI |
| Zygoptera | Coenagrion_ornatum_KP272503_Df1569_SVN_Coenagrion_ornatum_KP272504_Df1570_SVN | Z188 | NCBI |
| Zygoptera | Coenagrion_puella_ASG433_CO | Z44 | DS1 |
| Zygoptera | Coenagrion_puella_FM37_CS_Coenagrion_ornatum_GU682176_Coenagrion_puella_Coenagrion_puella_ZPLOD819-20_Coenagrion_puella_GA22_TN_Coenagrion_puella_GU682174_Coenagrion_puella_MG22_BA_Coenagrion_puella_MIBAG0036_LC_Coenagrion_puella_MIBAG0197_TS_Coenagrion_puella_MIBAG0208_VC_Coenagrion_puella_MIBAG0212_MO_Coenagrion_puella_MIBAG0261_CB_Coenagrion_puella_MIBAG0277_AQ_Coenagrion_puella_VF02_VT_Coenagrion_pulchellum_ASG432_CO_Coenagrion_pulchellum_GA01_TN_Coenagrion_pulchellum_KF369350_Coenagrion_puella_GLP16_PG_Coenagrion_puella_MIBAG0229_CN | Z45 | DS1 |
| Zygoptera | Coenagrion_puella_GA05_TN | Z46 | DS1 |
| Zygoptera | Coenagrion_puella_HM901887 | Z189 | NCBI |
| Zygoptera | Coenagrion_puella_KP272538_Df1705_MAR | Z190 | NCBI |
| Zygoptera | Coenagrion_puella_KP272539_Df1706_MAR_Coenagrion_puella_KP272540_Df1710_MAR_Coenagrion_puella_KP272541_Df1711_MAR | Z191 | NCBI |
| Zygoptera | Coenagrion_puella_KP272546_Df1809_FIN | Z192 | NCBI |
| Zygoptera | Coenagrion_puella_KP272547_Df1813_FIN | Z193 | NCBI |
| Zygoptera | Coenagrion_puella_KU695837_Df1709_MAR | Z194 | NCBI |
| Zygoptera | Coenagrion_puella_KU695838_Df1717_RUS_Coenagrion_puella_KU695842_Df1778_MKD_Coenagrion_puella_KU695843_Df1779_MKD_Coenagrion_puella_KU695844_Df1792_DEU_Coenagrion_puella_KU695845_Df1795_RUS | Z195 | NCBI |
| Zygoptera | Coenagrion_puella_KU695839_Df1718_NLD | Z196 | NCBI |
| Zygoptera | Coenagrion_puella_KU695840_Df1775_PRT_Coenagrion_puella_KU695841_Df1776_PRT | Z197 | NCBI |
| Zygoptera | Coenagrion_puella_LI33_ME_Coenagrion_puella_LI34_ME | Z49 | DS1 |
| Zygoptera | Coenagrion_puella_MH449990 | Z198 | NCBI |
| Zygoptera | Coenagrion_puella_MIBAG0095_CO_Coenagrion_puella_MIBAG0286_CO_Coenagrion_pulchellum_KF369349 | Z51 | DS1 |
| Zygoptera | Coenagrion_puella_MIBAG0131_PC | Z52 | DS1 |
| Zygoptera | Coenagrion_puella_MIBAG0181_PV_Coenagrion_puella_VF09_VT_Coenagrion_puella_MIBAG0294_MO | Z53 | DS1 |
| Zygoptera | Coenagrion_pulchellum_ASG431_CO_Coenagrion_pulchellum_GA08_TN_Coenagrion_pulchellum_MIBAG0011_CO | Z57 | DS1 |
| Zygoptera | Coenagrion_pulchellum_GLP17_TR_Coenagrion_pulchellum_MG04_FG_Coenagrion_pulchellum_MG13_LE_Coenagrion_pulchellum_SR06_LT_Coenagrion_pulchellum_SR07_LT_Coenagrion_pulchellum_SR18_LT_Coenagrion_pulchellum_SR20_LT | Z58 | DS1 |
| Zygoptera | Coenagrion_pulchellum_HM901861 | Z199 | NCBI |
| Zygoptera | Coenagrion_pulchellum_HM901888 | Z200 | NCBI |
| Zygoptera | Coenagrion_pulchellum_KP272544_Df1800_FIN_Coenagrion_pulchellum_KP272545_Df1801_FIN | Z201 | NCBI |
| Zygoptera | Coenagrion_pulchellum_SR08_LT | Z59 | DS1 |
| Zygoptera | Coenagrion_pulchellum_ZMBN958_17 | Z202 | BOLD |
| Zygoptera | Coenagrion_scitulum_FM10_BA | Z60 | DS1 |
| Zygoptera | Coenagrion_scitulum_KP272421_Coenagrion_scitulum_KP272423_Coenagrion_scitulum_KP272592 | Z61 | NCBI |
| Zygoptera | Coenagrion_scitulum_MIBAG0195_VA_Coenagrion_scitulum_MIBAG0196_TS | Z62 | DS1 |
| Zygoptera | Coenagrion_scitulum_MIBAG0252_CB | Z63 | DS1 |
| Anisoptera | Cordulegaster_bidentata_ASG450_TN_Cordulegaster_bidentata_GA38_LC | A36 | DS1 |
| Anisoptera | Cordulegaster_bidentata_DB08_CN | A37 | DS1 |
| Anisoptera | Cordulegaster_bidentata_KF584922 | A277 | NCBI |
| Anisoptera | Cordulegaster_bidentata_KF584923 | A278 | NCBI |
| Anisoptera | Cordulegaster_bidentata_KF584924 | A279 | NCBI |
| Anisoptera | Cordulegaster_bidentata_KF584925 | A280 | NCBI |
| Anisoptera | Cordulegaster_bidentata_KF584926 | A281 | NCBI |
| Anisoptera | Cordulegaster_bidentata_KF584927 | A282 | NCBI |
| Anisoptera | Cordulegaster_bidentata_KF584928 | A283 | NCBI |
| Anisoptera | Cordulegaster_bidentata_KF584929_PZ | A38 | NCBI |
| Anisoptera | Cordulegaster_bidentata_KF584930_PZ | A39 | NCBI |
| Anisoptera | Cordulegaster_bidentata_KF584931 | A284 | NCBI |
| Anisoptera | Cordulegaster_bidentata_KF584947_ME | A40 | NCBI |
| Anisoptera | Cordulegaster_bidentata_KF584958 | A285 | NCBI |
| Anisoptera | Cordulegaster_bidentata_KF584963 | A286 | NCBI |
| Anisoptera | Cordulegaster_bidentata_KF584971_ME | A41 | NCBI |
| Anisoptera | Cordulegaster_bidentata_KF584972_SR | A42 | NCBI |
| Anisoptera | Cordulegaster_bidentata_MH304666 | A287 | NCBI |
| Anisoptera | Cordulegaster_boltonii_FBAQU53110 | A288 | BOLD |
| Anisoptera | Cordulegaster_boltonii_GA31_TN_Cordulegaster_boltonii_IC25_UD | A43 | DS1 |
| Anisoptera | Cordulegaster_boltonii_GA48_TO | A44 | DS1 |

MOLECULAR ECOLOGY
RESOURCES

| Suborder | Species name/Sample ID | Haplotype | DNA barcode source |
|---|---|---|---|
| Anisoptera | Cordulegaster_boltonii_KF584933_FI_Cordulegaster_boltonii_MH304646_AL_Cordulegaster_boltonii_MH304646_AT_Cordulegaster_boltonii_MH304646_MC_Cordulegaster_boltonii_MH304646_RA | A45 | NCBI |
| Anisoptera | Cordulegaster_boltonii_KF584934_FI_Cordulegaster_boltonii_MH304650_FI_Cordulegaster_boltonii_MH304650_RA | A46 | NCBI |
| Anisoptera | Cordulegaster_boltonii_KF584953 | A290 | NCBI |
| Anisoptera | Cordulegaster_boltonii_KF584954 | A291 | NCBI |
| Anisoptera | Cordulegaster_boltonii_KF584955 | A292 | NCBI |
| Anisoptera | Cordulegaster_boltonii_KF584956 | A293 | NCBI |
| Anisoptera | Cordulegaster_boltonii_KF584957 | A294 | NCBI |
| Anisoptera | Cordulegaster_boltonii_KF584959 | A295 | NCBI |
| Anisoptera | Cordulegaster_boltonii_KF584961 | A297 | NCBI |
| Anisoptera | Cordulegaster_boltonii_KF584962 | A298 | NCBI |
| Anisoptera | Cordulegaster_boltonii_KF584964 | A299 | NCBI |
| Anisoptera | Cordulegaster_boltonii_KF584965 | A300 | NCBI |
| Anisoptera | Cordulegaster_boltonii_KF584967 | A301 | NCBI |
| Anisoptera | Cordulegaster_boltonii_KF584968 | A302 | NCBI |
| Anisoptera | Cordulegaster_boltonii_MH304647_AT_Cordulegaster_boltonii_MH304647_BS_Cordulegaster_boltonii_MH304647_TN_Cordulegaster_boltonii_KF584960 | A47 | NCBI |
| Anisoptera | Cordulegaster_boltonii_MH304648_AL_Cordulegaster_boltonii_KF584932 | A48 | NCBI |
| Anisoptera | Cordulegaster_boltonii_MH304649_AL | A49 | NCBI |
| Anisoptera | Cordulegaster_boltonii_ZMBN03715 | A303 | BOLD |
| Anisoptera | Cordulegaster_heros_IC21_GO | A50 | DS1 |
| Anisoptera | Cordulegaster_heros_IC22_TS_Cordulegaster_heros_IC23_UD | A51 | DS1 |
| Anisoptera | Cordulegaster_heros_KF584940 | A305 | NCBI |
| Anisoptera | Cordulegaster_heros_ZPLOD820-20 | A304 | BOLD |
| Anisoptera | Cordulegaster_trinacriae_FM46_BA | A53 | DS1 |
| Anisoptera | Cordulegaster_trinacriae_KF584945_Sicily | A54 | NCBI |
| Anisoptera | Cordulegaster_trinacriae_KF584946_Cordulegaster_trinacriae_MH304653_Sicily | A55 | NCBI |
| Anisoptera | Cordulegaster_trinacriae_MH304651_PZ | A56 | NCBI |
| Anisoptera | Cordulegaster_trinacriae_MH304654_ME | A57 | NCBI |
| Anisoptera | Cordulegaster_trinacriae_MH304656 | A58 | NCBI |
| Anisoptera | Cordulegaster_trinacriae_MH304658_SR | A59 | NCBI |
| Anisoptera | Cordulegaster_trinacriae_MH304661_VV | A60 | NCBI |
| Anisoptera | Cordulegaster_trinacriae_MH304662_VV | A61 | NCBI |
| Anisoptera | Cordulegaster_trinacriae_MH304663_PZ | A62 | NCBI |
| Anisoptera | Cordulegaster_trinacriae_MH304664_BT | A63 | NCBI |
| Anisoptera | Cordulegaster_trinacriae_MH304665_PZ | A64 | NCBI |
| Anisoptera | Cordulegaster_trinacriae_MIBAG0275_CB | A65 | DS1 |
| Anisoptera | Cordulia_aenea_FBAQU489_10_Cordulia_aenea_GBMHO494_14_Cordulia_aenea_MIBAG0185_VA | A68 | DS1 / BOLD |
| Anisoptera | Cordulia_aenea_GA3_TN_Cordulia_aenea_GLP19_TR | A66 | DS1 |
| Anisoptera | Cordulia_aenea_ZPLOD821-20 | A306 | BOLD |
| Anisoptera | Crocothemis_erythraea_ASG466_SS | A69 | DS1 |
| Anisoptera | Crocothemis_erythraea_GU682175 | A308 | NCBI |
| Anisoptera | Crocothemis_erythraea_KC912238_Crocothemis_erythraea_KY847584 | A309 | NCBI |
| Anisoptera | Crocothemis_erythraea_KC912239_Crocothemis_erythraea_KC912243_Crocothemis_erythraea_KY847579_Crocothemis_erythraea_KY847580 | A310 | NCBI |
| Anisoptera | Crocothemis_erythraea_KC912240_Crocothemis_erythraea_KC912242_Crocothemis_erythraea_KY847582_Crocothemis_erythraea_KY847583 | A311 | NCBI |
| Anisoptera | Crocothemis_erythraea_KC912241_Crocothemis_erythraea_KY847585 | A312 | NCBI |
| Anisoptera | Crocothemis_erythraea_KC912244_Crocothemis_erythraea_KY847581 | A313 | NCBI |
| Anisoptera | Crocothemis_erythraea_MIBAG0064_MB | A70 | DS1 |
| Anisoptera | Crocothemis_erythraea_MIBAG0243_CB | A71 | DS1 |
| Anisoptera | Crocothemis_erythraea_SISAF083_12 | A314 | BOLD |
| Anisoptera | Crocothemis_erythraea_SR22_LT | A72 | DS1 |
| Anisoptera | Crocothemis_erythraea_SS11_TP | A73 | DS1 |
| Anisoptera | Crocothemis_erythraea_ZPLOD822-20 | A307 | BOLD |
| Anisoptera | Diplacodes_lefebvrii_ASG742_SU | A74 | DS1 |
| Anisoptera | Diplacodes_lefebvrii_FL23_CA | A75 | DS1 |
| Zygoptera | Enallagma_cyathigerum_DB20_AO_Enallagma_cyathigerum_MIBAG0132_PC_Enallagma_cyathigerum_ZMBN960_17 | Z65 | DS1 |
| Zygoptera | Enallagma_cyathigerum_FBAQU1504_13_Enallagma_cyathigerum_GU682173_Enallagma_cyathigerum_MIBAG0057_LC | Z67 | DS1 / NCBI / BOLD |
| Zygoptera | Enallagma_cyathigerum_GU682183 | Z203 | NCBI |
| Zygoptera | Enallagma_cyathigerum_HETFI049_11_Enallagma_cyathigerum_HETFI052_11 | Z204 | BOLD |
| Zygoptera | Enallagma_cyathigerum_HQ563103 | Z205 | NCBI |
| Zygoptera | Enallagma_cyathigerum_HQ563105 | Z206 | NCBI |
| Zygoptera | Enallagma_cyathigerum_KC912310_Enallagma_cyathigerum_KC912311_Enallagma_cyathigerum_KC912313_Enallagma_cyathigerum_KC912314 | Z208 | NCBI |
| Zygoptera | Enallagma_cyathigerum_KC912312 | Z209 | NCBI |
| Zygoptera | Enallagma_cyathigerum_KX263691 | Z210 | NCBI |
| Zygoptera | Enallagma_cyathigerum_KX263692 | Z211 | NCBI |
| Zygoptera | Enallagma_cyathigerum_LI22_ME_Enellagma_cyathigerum_MIBAG0280_AQ | Z66 | DS1 |
| Zygoptera | Enallagma_cyathigerum_MIBAG0216_MO | Z69 | DS1 |
| Zygoptera | Enallagma_cyathigerum_XJDQD086_18 | Z212 | BOLD |
| Zygoptera | Erythromma_lindenii_AC09_AG_Erythromma_lindenii_AC11_TP_Erythromma_lindenii_AC12_SR_Erythromma_lindenii_FM14_BA_Erythromma_lindenii_LI29_CT_Erythromma_lindenii_MIBAG0056_LC_Erythromma_lindenii_MIBAG0206_VA_Erythromma_lindenii_MIBAG0248_CB_Erythromma_lindenii_MIBAG0276_AQ_Erythromma_lindenii_MIBAG0149_MO | Z71 | DS1 |
| Zygoptera | Erythromma_lindenii_ASG706_SS_Erythromma_lindenii_ASG768_NU_Erythromma_lindenii_FL27_VS | Z72 | DS1 |
| Zygoptera | Erythromma_lindenii_ASG772_TN_Erythromma_lindenii_GA36_TN_Erythromma_lindenii_LI04_BS_Erythromma_lindenii_LI06_BS_Erythromma_lindenii_ASG795_TN | Z73 | DS1 |
| Zygoptera | Erythromma_lindenii_ASG773_TN_Erythromma_lindenii_ASG794_TN_Erythromma_lindenii_ASG796_TN_Erythromma_lindenii_GS08_VC_Erythromma_lindenii_IC29_UD | Z74 | DS1 |
| Zygoptera | Erythromma_lindenii_ASG774_TN | Z75 | DS1 |
| Zygoptera | Erythromma_lindenii_FBAQU491_10 | Z213 | BOLD |
| Zygoptera | Erythromma_lindenii_FBAQU562_10 | Z214 | BOLD |
| Zygoptera | Erythromma_lindenii_FM22_BT | Z77 | DS1 |

MOLECULAR ECOLOGY RESOURCES

| Suborder | Species name/Sample ID | Haplotype | DNA barcode source |
|---|---|---|---|
| Zygoptera | Erythromma_lindenii_FM30_LE | Z78 | DS1 |
| Zygoptera | Erythromma_lindenii_FM33_LE | Z79 | DS1 |
| Zygoptera | Erythromma_lindenii_GLP21_PG | Z80 | DS1 |
| Zygoptera | Erythromma_lindenii_IC03_UD_Erythromma_lindenii_IC35_PN | Z81 | DS1 |
| Zygoptera | Erythromma_lindenii_IC32_TS | Z83 | DS1 |
| Zygoptera | Erythromma_lindenii_MF458702 | Z216 | NCBI |
| Zygoptera | Erythromma_lindenii_MIBAG0053_LC | Z85 | DS1 |
| Zygoptera | Erythromma_lindenii_MIBAG0084_MI | Z86 | DS1 |
| Zygoptera | Erythromma_lindenii_MIBAG0268_CB | Z88 | DS1 |
| Zygoptera | Erythromma_lindenii_ZPLOD823-20 | Z215 | BOLD |
| Zygoptera | Erythromma_lindenii_ZPLOD843-20 | Z217 | BOLD |
| Zygoptera | Erythromma_najas_EF176683_Erythromma_najas_EF176688_Erythromma_najas_EF176689_Erythromma_najas_EF176690_Erythromma_najas_EF176692_Erythromma_najas_EF176697_Erythromma_najas_EF176701_Erythromma_najas_EF176702_Erythromma_najas_EF176704_Erythromma_najas_EF176706_Erythromma_najas_EF176708_Erythromma_najas_EF176709_Erythromma_najas_EF176710_Erythromma_najas_EF176713_Erythromma_najas_EF176714_Erythromma_najas_EF176717_Erythromma_najas_EF176719_Erythromma_najas_EF176726_Erythromma_najas_EF176727_Erythromma_najas_EF176732_Erythromma_najas_EF176733_Erythromma_najas_EF176734_Erythromma_najas_EF176735_Erythromma_najas_EF176736_Erythromma_najas_EF176738_Erythromma_najas_EF176740_Erythromma_najas_EF176742_Erythromma_najas_EF176745_Erythromma_najas_EF176753_Erythromma_najas_EF176754_Erythromma_najas_EF176755_Erythromma_najas_EF176759_Erythromma_najas_EF176762 | Z218 | NCBI |
| Zygoptera | Erythromma_najas_EF176684_Erythromma_najas_EF176686_Erythromma_najas_EF176687_Erythromma_najas_EF176691_Erythromma_najas_EF176694_Erythromma_najas_EF176715_Erythromma_najas_EF176716_Erythromma_najas_EF176718_Erythromma_najas_EF176720_Erythromma_najas_EF176722_Erythromma_najas_EF176724_Erythromma_najas_EF176725_Erythromma_najas_EF176729_Erythromma_najas_EF176730_Erythromma_najas_EF176737_Erythromma_najas_EF176739_Erythromma_najas_EF176741_Erythromma_najas_EF176747_Erythromma_najas_EF176763_Erythromma_najas_EF176764_Erythromma_najas_EF176765_Erythromma_najas_EF176767_Erythromma_najas_EF176768_Erythromma_najas_EF176769 | Z219 | NCBI |
| Zygoptera | Erythromma_najas_EF176685 | Z220 | NCBI |
| Zygoptera | Erythromma_najas_EF176693_Erythromma_najas_EF176696_Erythromma_najas_EF176699_Erythromma_najas_EF176711_Erythromma_najas_EF176760 | Z221 | NCBI |
| Zygoptera | Erythromma_najas_EF176695_Erythromma_najas_EF176700_Erythromma_najas_EF176750_Erythromma_najas_EF176751_Erythromma_najas_EF176752_Erythromma_najas_EF176757 | Z222 | NCBI |
| Zygoptera | Erythromma_najas_EF176698 | Z223 | NCBI |
| Zygoptera | Erythromma_najas_EF176703_Erythromma_najas_EF176712_Erythromma_najas_EF176731_Erythromma_najas_EF176746_Erythromma_najas_EF176748_Erythromma_najas_EF176749_Erythromma_najas_EF176756 | Z224 | NCBI |
| Zygoptera | Erythromma_najas_EF176705 | Z225 | NCBI |
| Zygoptera | Erythromma_najas_EF176707 | Z226 | NCBI |
| Zygoptera | Erythromma_najas_EF176721 | Z227 | NCBI |
| Zygoptera | Erythromma_najas_EF176723 | Z228 | NCBI |
| Zygoptera | Erythromma_najas_EF176728 | Z229 | NCBI |
| Zygoptera | Erythromma_najas_EF176743 | Z230 | NCBI |
| Zygoptera | Erythromma_najas_EF176744 | Z231 | NCBI |
| Zygoptera | Erythromma_najas_EF176758 | Z232 | NCBI |
| Zygoptera | Erythromma_najas_EF176761 | Z233 | NCBI |
| Zygoptera | Erythromma_najas_EF176766 | Z234 | NCBI |
| Zygoptera | Erythromma_najas_FBAQU563_10 | Z235 | BOLD |
| Zygoptera | Erythromma_najas_GA21_TN | Z89 | DS1 |
| Zygoptera | Erythromma_najas_MIBAG0193_VA | Z90 | DS1 |
| Zygoptera | Erythromma_najas_MN345616 | Z236 | NCBI |
| Zygoptera | Erythromma_viridulum_ASG683_SS | Z91 | DS1 |
| Zygoptera | Erythromma_viridulum_FBAQU314_09_Erythromma_viridulum_LI17_SR | Z93 | DS1 / BOLD |
| Zygoptera | Erythromma_viridulum_FBAQU565_10 | Z237 | BOLD |
| Zygoptera | Erythromma_viridulum_FM31_LE_Erythromma_viridulum_GA37_TN_Erythromma_viridulum_MIBAG0281_AQ | Z92 | DS1 |
| Zygoptera | Erythromma_viridulum_MIBAG0052_LC | Z94 | DS1 |
| Zygoptera | Erythromma_viridulum_SS12_TP | Z95 | DS1 |
| Anisoptera | Gomphus_flavipes_KX891019 | A315 | NCBI |
| Anisoptera | Gomphus_flavipes_LI2_PV | A162 | DS1 |
| Anisoptera | Gomphus_flavipes_XJDQD087_18_Gomphus_flavipes_XJDQD088_18_Gomphus_flavipes_XJDQD089_18 | A316 | BOLD |
| Anisoptera | Gomphus_vulgatissimus_FBAQU1445_13 | A317 | BOLD |
| Anisoptera | Gomphus_vulgatissimus_FBAQU534_10 | A318 | BOLD |
| Anisoptera | Gomphus_vulgatissimus_FL08_AN_Gomphus_vulgatissimus_MIBAG0017_CO_Gomphus_vulgatissimus_IC02_UD | A76 | DS1 |
| Anisoptera | Gomphus_vulgatissimus_GBMIN88707_17 | A319 | BOLD |
| Zygoptera | Ischnura_elegans_FBAQU495_10 | Z238 | BOLD |
| Zygoptera | Ischnura_elegans_FBAQU535_10 | Z239 | BOLD |
| Zygoptera | Ischnura_elegans_FBAQU536_10 | Z240 | BOLD |
| Zygoptera | Ischnura_elegans_GA09_TN_Ischnura_elegans_KF369415 | Z96 | DS1 |
| Zygoptera | Ischnura_elegans_GA50_TO_Ischnura_elegans_MIBAG0178_GR_Ischnura_elegans_VF14_VT | Z97 | DS1 |
| Zygoptera | Ischnura_elegans_GQ256030_Ischnura_elegans_GQ256031_Ischnura_elegans_GQ256032 | Z242 | NCBI |
| Zygoptera | Ischnura_elegans_HM376192 | Z243 | NCBI |
| Zygoptera | Ischnura_elegans_HQ834803 | Z244 | NCBI |
| Zygoptera | Ischnura_elegans_KF257118 | Z245 | NCBI |
| Zygoptera | Ischnura_elegans_KY127432 | Z247 | NCBI |
| Zygoptera | Ischnura_elegans_KY127433 | Z248 | NCBI |
| Zygoptera | Ischnura_elegans_KY127434 | Z249 | NCBI |
| Zygoptera | Ischnura_elegans_KY127437 | Z250 | NCBI |
| Zygoptera | Ischnura_elegans_KY127438 | Z251 | NCBI |
| Zygoptera | Ischnura_elegans_KY127439 | Z252 | NCBI |
| Zygoptera | Ischnura_elegans_KY127440 | Z253 | NCBI |
| Zygoptera | Ischnura_elegans_MF458738 | Z254 | NCBI |
| Zygoptera | Ischnura_elegans_MG14_LE_Ischnura_elegans_GAM11_MON_5 | Z98 | DS1 |
| Zygoptera | Ischnura_elegans_XJDQD082_18_Ischnura_elegans_XJDQD084_18_Ischnura_elegans_XJDQD090_18_Ischnura_elegans_XJDQD091_18 | Z255 | BOLD |
| Zygoptera | Ischnura_elegans_XJDQD083_18 | Z256 | BOLD |
| Zygoptera | Ischnura_elegans_ZPLOD826-20_Ischnura_elegans_HQ563104_Ischnura_elegans_KU958378_Ischnura_elegans_MH449982_Ischnura_elegans_MH449993 | Z241 | NCBI / BOLD |
| Zygoptera | Ischnura_genei_ASG463_SS | Z99 | DS1 |

| Suborder | Species name/Sample ID | Haplotype | DNA barcode source |
|---|---|---|---|
| Zygoptera | Ischnura_genei_LI16_SR | Z100 | DS1 |
| Zygoptera | Ischnura_genei_SS05_TP | Z101 | DS1 |
| Zygoptera | Ischnura_pumilio_FBAQU496_10_Ischnura_pumilio_FBAQU538_10 | Z257 | BOLD |
| Zygoptera | Ischnura_pumilio_FBAQU537_10 | Z258 | BOLD |
| Zygoptera | Ischnura_pumilio_FM08_BA_Ischnura_pumilio_MIBAG0213_MO_Ischnura_pumilio_GA29_TN_Ischnura_pumilio_GA53_TO_Ischnura_pumilio_HM422049_Ischnura_pumilio_KC878732_Ischnura_pumilio_MH449985 | Z102 | DS1 |
| Zygoptera | Lestes_barbarus_ASG676_SS_Lestes_barbarus_MIBAG0260_CB | Z104 | DS1 |
| Zygoptera | Lestes_barbarus_MIBAG0112_TA | Z105 | DS1 |
| Zygoptera | Lestes_barbarus_MIBAG0157_FE | Z106 | DS1 |
| Zygoptera | Lestes_dryas_BBODA194_10_Lestes_dryas_KM528476_Lestes_dryas_KM528857_Lestes_dryas_KM530028_Lestes_dryas_KM531276_Lestes_dryas_KM531529_Lestes_dryas_KM532506_Lestes_dryas_KM534872_Lestes_dryas_KM534980_Lestes_dryas_KM535507_Lestes_dryas_KM535977_Lestes_dryas_KM536823_Lestes_dryas_KM537318_Lestes_dryas_KU875364 | Z259 | NCBI |
| Zygoptera | Lestes_dryas_DB26_SV | Z107 | DS1 |
| Zygoptera | Lestes_dryas_FM13_BA | Z108 | DS1 |
| Zygoptera | Lestes_dryas_GA64_TO_Lestes_dryas_Lestes_dryas_ZPLOD827-20 | Z109 | DS1 / BOLD |
| Zygoptera | Lestes_dryas_KM530438 | Z260 | NCBI |
| Zygoptera | Lestes_dryas_KM531733 | Z261 | NCBI |
| Zygoptera | Lestes_dryas_KM531862 | Z262 | NCBI |
| Zygoptera | Lestes_dryas_KM532810_Lestes_dryas_ODRMA014_10 | Z263 | NCBI / BOLD |
| Zygoptera | Lestes_dryas_KM534143 | Z264 | NCBI |
| Zygoptera | Lestes_dryas_KM535968 | Z265 | NCBI |
| Zygoptera | Lestes_dryas_KM537254 | Z266 | NCBI |
| Zygoptera | Lestes_dryas_KM537776 | Z267 | NCBI |
| Zygoptera | Lestes_dryas_LI24_ME | Z110 | DS1 |
| Zygoptera | Lestes_dryas_MIBAG0128_PC | Z111 | DS1 |
| Zygoptera | Lestes_dryas_MIBAG0282_AQ | Z112 | DS1 |
| Zygoptera | Lestes_macrostigma_FM27_LE | Z113 | DS1 |
| Zygoptera | Lestes_sponsa_DB01_AO | Z114 | DS1 |
| Zygoptera | Lestes_sponsa_GA34_TN | Z115 | DS1 |
| Zygoptera | Lestes_sponsa_MIBAG0054_LC | Z116 | DS1 |
| Zygoptera | Lestes_virens_ASG750_SU_Lestes_virens_MIBAG0122_PC_Lestes_virens_MIBAG0263_CB | Z117 | DS1 |
| Zygoptera | Lestes_virens_FBAQU1427_13 | Z268 | BOLD |
| Zygoptera | Lestes_virens_GA51_TO | Z118 | DS1 |
| Zygoptera | Lestes_virens_KF369424 | Z269 | NCBI |
| Zygoptera | Lestes_virens_LI19_CT_Lestes_virens_LI36_ME_Lestes_virens_SS13_TP | Z119 | DS1 |
| Zygoptera | Lestes_virens_MIBAG0104_CO | Z121 | DS1 |
| Anisoptera | Leucorrhinia_dubia_DB15_AO | A78 | DS1 |
| Anisoptera | Leucorrhinia_dubia_FBAQU500_10 | A320 | BOLD |
| Anisoptera | Leucorrhinia_dubia_FBAQU501_10 | A321 | BOLD |
| Anisoptera | Leucorrhinia_dubia_GA20_TN_Leucorrhinia_dubia_MIBAG0093_VB_Leucorrhinia_dubia_MIBAG0219_SO | A79 | DS1 |
| Anisoptera | Leucorrhinia_dubia_ZPLOD828-20 | A322 | BOLD |
| Anisoptera | Leucorrhinia_pectoralis_GA25_TN | A80 | DS1 |
| Anisoptera | Libellula_depressa_FBAQU502_10_Libellula_depressa_GA06_TN_Libellula_depressa_GA58_TO_Libellula_depressa_MIBAG0115_PC | A82 | DS1 / BOLD |
| Anisoptera | Libellula_depressa_FM05_BT | A81 | DS1 |
| Anisoptera | Libellula_depressa_MIBAG0258_CB | A84 | DS1 |
| Anisoptera | Libellula_fulva_FM34_LE | A85 | DS1 |
| Anisoptera | Libellula_fulva_GA10_TN | A86 | DS1 |
| Anisoptera | Libellula_fulva_MIBAG0045_LC | A87 | DS1 |
| Anisoptera | Libellula_fulva_SR05_LT | A88 | DS1 |
| Anisoptera | Libellula_quadrimaculata_DB03_AO | A89 | DS1 |
| Anisoptera | Libellula_quadrimaculata_FBAQU503_10 | A323 | BOLD |
| Anisoptera | Libellula_quadrimaculata_GA04_TN_Libellula_quadrimaculata_GLP29_TR | A90 | DS1 |
| Anisoptera | Libellula_quadrimaculata_HM399588 | A325 | NCBI |
| Anisoptera | Libellula_quadrimaculata_HM399589 | A326 | NCBI |
| Anisoptera | Libellula_quadrimaculata_HM399590 | A327 | NCBI |
| Anisoptera | Libellula_quadrimaculata_HM413521 | A328 | NCBI |
| Anisoptera | Libellula_quadrimaculata_HM413529_Libellula_quadrimaculata_HM413566 | A329 | NCBI |
| Anisoptera | Libellula_quadrimaculata_JF839248 | A330 | NCBI |
| Anisoptera | Libellula_quadrimaculata_JF839249 | A331 | NCBI |
| Anisoptera | Libellula_quadrimaculata_JF839307 | A332 | NCBI |
| Anisoptera | Libellula_quadrimaculata_JF839308_Libellula_quadrimaculata_JN294404_Libellula_quadrimaculata_JN294440_Libellula_quadrimaculata_KM533579_Libellula_quadrimaculata_KM536673 | A333 | NCBI |
| Anisoptera | Libellula_quadrimaculata_JN294338 | A334 | NCBI |
| Anisoptera | Libellula_quadrimaculata_JN294400 | A335 | NCBI |
| Anisoptera | Libellula_quadrimaculata_JN294401 | A336 | NCBI |
| Anisoptera | Libellula_quadrimaculata_JN294403 | A337 | NCBI |
| Anisoptera | Libellula_quadrimaculata_JN294437 | A338 | NCBI |
| Anisoptera | Libellula_quadrimaculata_JN294439 | A339 | NCBI |
| Anisoptera | Libellula_quadrimaculata_JN294441 | A340 | NCBI |
| Anisoptera | Libellula_quadrimaculata_JN419954 | A341 | NCBI |
| Anisoptera | Libellula_quadrimaculata_KF257060 | A342 | NCBI |
| Anisoptera | Libellula_quadrimaculata_KM528898 | A343 | NCBI |
| Anisoptera | Libellula_quadrimaculata_KM530724 | A344 | NCBI |
| Anisoptera | Libellula_quadrimaculata_KM531593 | A345 | NCBI |
| Anisoptera | Libellula_quadrimaculata_KM531766 | A346 | NCBI |
| Anisoptera | Libellula_quadrimaculata_KM537510 | A347 | NCBI |
| Anisoptera | Libellula_quadrimaculata_KR144312 | A348 | NCBI |
| Anisoptera | Libellula_quadrimaculata_KR919033_Libellula_quadrimaculata_MG463189 | A349 | NCBI |

MOLECULAR ECOLOGY RESOURCES

| Suborder | Species name/Sample ID | Haplotype | DNA barcode source |
|---|---|---|---|
| Anisoptera | Libellula_quadrimaculata_KR919420 | A350 | NCBI |
| Anisoptera | Libellula_quadrimaculata_KU875373 | A351 | NCBI |
| Anisoptera | Libellula_quadrimaculata_KU875374 | A352 | NCBI |
| Anisoptera | Libellula_quadrimaculata_MG376152 | A353 | NCBI |
| Anisoptera | Libellula_quadrimaculata_MG378286 | A354 | NCBI |
| Anisoptera | Libellula_quadrimaculata_MG381893 | A355 | NCBI |
| Anisoptera | Libellula_quadrimaculata_MIBAG0060_MB | A91 | DS1 |
| Anisoptera | Libellula_quadrimaculata_TZBCA545_07 | A356 | BOLD |
| Anisoptera | Libellula_quadrimaculata_TZBCA546_07 | A357 | BOLD |
| Anisoptera | Libellula_quadrimaculata_ZPLOD829-20 | A324 | BOLD |
| Anisoptera | Lindenia_tetraphylla_ASG722_OR_Lindenia_tetraphylla_SS18_TP | A92 | DS1 |
| Anisoptera | Lindenia_tetraphylla_KJ873214 | A358 | NCBI |
| Anisoptera | Lindenia_tetraphylla_KX241516_SI | A94 | NCBI |
| Anisoptera | Lindenia_tetraphylla_ZPLOD830-20_Lindenia_tetraphylla_GLP30_PG_Lindenia_tetraphylla_MIBAG0239_CB_Lindenia_tetraphylla_MIBAG0272_CB | A93 | DS1 / BOLD |
| Zygoptera | Nehalennia_speciosa_AM696290 | Z270 | NCBI |
| Zygoptera | Nehalennia_speciosa_FBAQU1428_13 | Z271 | BOLD |
| Zygoptera | Nehalennia_speciosa_FBAQU541_10 | Z272 | BOLD |
| Zygoptera | Nehalennia_speciosa_FN252223 | Z273 | NCBI |
| Zygoptera | Nehalennia_speciosa_FN252224 | Z274 | NCBI |
| Zygoptera | Nehalennia_speciosa_FN252226 | Z275 | NCBI |
| Zygoptera | Nehalennia_speciosa_FN252227 | Z276 | NCBI |
| Zygoptera | Nehalennia_speciosa_FN252228 | Z277 | NCBI |
| Zygoptera | Nehalennia_speciosa_FN252229 | Z278 | NCBI |
| Zygoptera | Nehalennia_speciosa_FN252231 | Z279 | NCBI |
| Zygoptera | Nehalennia_speciosa_ICA_UD | Z122 | DS1 |
| Zygoptera | Nehalennia_speciosa_VO1_VA | Z123 | DS1 |
| Anisoptera | Onychogomphus_forcipatus_ASG451_TN_Onychogomphus_forcipatus_MIBAG0043_LC_Onychogomphus_forcipatus_MIBAG0136_MO_Onychogomphus_forcipatus_ASG771_TN_Onychogomphus_forcipatus_ASG822_TN_Onychogomphus_forcipatus_MIBAG0110_TA_Onychogomphus_forcipatus_MIBAG0245_CB | A96 | DS1 |
| Anisoptera | Onychogomphus_forcipatus_ASG456_TN_Onychogomphus_forcipatus_ASG781_TN_Onychogomphus_forcipatus_ASG823_TN | A97 | DS1 |
| Anisoptera | Onychogomphus_forcipatus_FBAQU504_10_Onychogomphus_forcipatus_FBAQU544_10 | A359 | BOLD |
| Anisoptera | Onychogomphus_forcipatus_FBAQU542_10 | A360 | BOLD |
| Anisoptera | Onychogomphus_forcipatus_FBAQU543_10 | A361 | BOLD |
| Anisoptera | Onychogomphus_forcipatus_GODO002_18 | A363 | BOLD |
| Anisoptera | Onychogomphus_forcipatus_GODO003_18_Onychogomphus_forcipatus_GODO004_18 | A364 | BOLD |
| Anisoptera | Onychogomphus_forcipatus_IC17_PN | A102 | DS1 |
| Anisoptera | Onychogomphus_forcipatus_IC19_GO | A103 | DS1 |
| Anisoptera | Onychogomphus_forcipatus_KF584975 | A365 | NCBI |
| Anisoptera | Onychogomphus_forcipatus_KJ873220 | A366 | NCBI |
| Anisoptera | Onychogomphus_forcipatus_LI08_ME_Onychogomphus_forcipatus_LI09_ME | A104 | DS1 |
| Anisoptera | Onychogomphus_forcipatus_ZPLOD831-20 | A362 | BOLD |
| Anisoptera | Onychogomphus_uncatus_AC02_SR | A107 | DS1 |
| Anisoptera | Onychogomphus_uncatus_DB07_GE_Onychogomphus_uncatus_MIBAG0069_MI_Onychogomphus_uncatus_MIBAG0259_CB | A108 | DS1 |
| Anisoptera | Onychogomphus_uncatus_KX891032 | A367 | NCBI |
| Anisoptera | Ophiogomphus_cecilia_KX891027_Ophiogomphus_cecilia_MIBAG0066_PV | A109 | DS1 / NCBI |
| Anisoptera | Orthetrum_albistylum_GA46_TO_Orthetrum_albistylum_MIBAG0073_MI | A110 | DS1 |
| Anisoptera | Orthetrum_albistylum_KF257070 | A368 | NCBI |
| Anisoptera | Orthetrum_albistylum_KF966558 | A369 | NCBI |
| Anisoptera | Orthetrum_albistylum_MF358739 | A370 | NCBI |
| Anisoptera | Orthetrum_albistylum_MF358740 | A371 | NCBI |
| Anisoptera | Orthetrum_albistylum_MF358741 | A372 | NCBI |
| Anisoptera | Orthetrum_brunneum_ASG469_SS | A111 | DS1 |
| Anisoptera | Orthetrum_brunneum_ASG711_SS_Orthetrum_brunneum_ASG737_SU_Orthetrum_brunneum_DB11_SP | A112 | DS1 |
| Anisoptera | Orthetrum_brunneum_FBAQU319_09_Orthetrum_brunneum_LI07_ME | A115 | DS1 / BOLD |
| Anisoptera | Orthetrum_brunneum_GA16_TN_Orthetrum_brunneum_GLP32_PG | A114 | DS1 |
| Anisoptera | Orthetrum_brunneum_SS01_TP | A116 | DS1 |
| Anisoptera | Orthetrum_cancellatum_ASG677_SS | A117 | DS1 |
| Anisoptera | Orthetrum_cancellatum_FBAQU505_10 | A373 | BOLD |
| Anisoptera | Orthetrum_cancellatum_FL18_MC_Orthetrum_cancellatum_SS07_TP | A118 | DS1 |
| Anisoptera | Orthetrum_cancellatum_GA17_TN | A119 | DS1 |
| Anisoptera | Orthetrum_cancellatum_GLP33_PG | A120 | DS1 |
| Anisoptera | Orthetrum_cancellatum_LI20_CT | A121 | DS1 |
| Anisoptera | Orthetrum_cancellatum_MG20_BR | A122 | DS1 |
| Anisoptera | Orthetrum_cancellatum_MIBAG0070_MI | A123 | DS1 |
| Anisoptera | Orthetrum_cancellatum_ZPLOD832-20 | A374 | BOLD |
| Anisoptera | Orthetrum_coerulescens_AC16_ME_Orthetrum_coerulescens_AC17_CT_Orthetrum_coerulescens_AC18_AG_Orthetrum_coerulescens_GLP34_PG_Orthetrum_coerulescens_LI14_SR_Orthetrum_coerulescens_MIBAG0237_CB_Orthetrum_coerulescens_MIBAG0287_CO_Orthetrum_coerulescens_AC22_TP_Orthetrum_coerulescens_GA57_TO | A124 | DS1 |
| Anisoptera | Orthetrum_coerulescens_ASG690_SS_Orthetrum_coerulescens_ASG731_SU_Orthetrum_coerulescens_FBAQU1439_13_Orthetrum_coerulescens_FBAQU1440_13_Orthetrum_coerulescens_FBAQU1442_13_Orthetrum_coerulescens_HM422051 | A128 | DS1 / NCBI / BOLD |
| Anisoptera | Orthetrum_coerulescens_FBAQU506_10 | A375 | BOLD |
| Anisoptera | Orthetrum_coerulescens_GA12_TN | A129 | DS1 |
| Anisoptera | Orthetrum_coerulescens_KC912263_FR_Orthetrum_coerulescens_KC912264_FR_Orthetrum_coerulescens_KC912265_FR | A131 | NCBI |
| Anisoptera | Orthetrum_coerulescens_KC912266_Orthetrum_coerulescens_KC912267_Orthetrum_coerulescens_KC912268_Orthetrum_coerulescens_KC912269_Orthetrum_coerulescens_KC912270_Orthetrum_coerulescens_KC912271 | A377 | NCBI |
| Anisoptera | Orthetrum_coerulescens_SR32_RM | A132 | DS1 |
| Anisoptera | Orthetrum_coerulescens_ZMBN329_16 | A378 | BOLD |
| Anisoptera | Orthetrum_coerulescens_ZPLOD833-20 | A376 | BOLD |
| Anisoptera | Orthetrum_nitidinerve_ASG716_NU_Orthetrum_nitidinerve_ASG758_SU | A133 | DS1 |
| Anisoptera | Orthetrum_trinacria_ASG458_SS | A134 | DS1 |

| Suborder | Species name/Sample ID | Haplotype | DNA barcode source |
|---|---|---|---|
| Anisoptera | Orthetrum_trinacria_KC912282_Orthetrum_trinacria_KC912283_Orthetrum_trinacria_KC912284_Orthetrum_trinacria_KC912285_Orthetrum_trinacria_KC912286_Orthetrum_trinacria_KY847595_Orthetrum_trinacria_KY847596_Orthetrum_trinacria_KY847597_Orthetrum_trinacria_KY847598_Orthetrum_trinacria_KY847599 | A379 | NCBI |
| Anisoptera | Orthetrum_trinacria_LI30_CT_Orthetrum_trinacria_SS17_TP | A135 | DS1 |
| Anisoptera | Oxygastra_curtisii_KX241515_PU | A136 | NCBI |
| Anisoptera | Oxygastra_curtisii_MIBAG0041_BG | A137 | DS1 |
| Anisoptera | Oxygastra_curtisii_MIBAG0270_CB | A138 | DS1 |
| Anisoptera | Paragomphus_genei_ASG464_SS_Paragomphus_genei_ASG761_NU | A139 | DS1 |
| Anisoptera | Paragomphus_genei_KU566310 | A380 | NCBI |
| Anisoptera | Paragomphus_genei_KX891031 | A381 | NCBI |
| Anisoptera | Paragomphus_genei_MN345045 | A382 | NCBI |
| Anisoptera | Paragomphus_genei_SS25_AG | A141 | DS1 |
| Zygoptera | Platycnemis_pennipes_FM06_TA | Z124 | DS1 |
| Zygoptera | Platycnemis_pennipes_GA14_TN_Platycnemis_pennipes_GA61_TO_Platycnemis_pennipes_GS04_VC_Platycnemis_pennipes_MIBAG0179_GR_Platycnemis_pennipes_VF11_VT | Z125 | DS1 |
| Zygoptera | Platycnemis_pennipes_KF369498 | Z281 | NCBI |
| Zygoptera | Platycnemis_pennipes_MIBAG0038_LC | Z128 | DS1 |
| Zygoptera | Platycnemis_pennipes_MIBAG0271_CB | Z129 | DS1 |
| Zygoptera | Platycnemis_pennipes_ZPLOD834-20 | Z280 | BOLD |
| Zygoptera | Pyrrhosoma_nymphula_FBAQU509_10_Pyrrhosoma_nymphula_FBAQU511_10_Pyrrhosoma_nymphula_FBAQU547_10 | Z281 | BOLD |
| Zygoptera | Pyrrhosoma_nymphula_FBAQU510_10 | Z282 | BOLD |
| Zygoptera | Pyrrhosoma_nymphula_GA41_TN_Pyrrhosoma_nymphula_ZPLOD835-20_Pyrrhosoma_nymphula_MIBAG0016_CO | Z130 | DS1 / BOLD |
| Zygoptera | Pyrrhosoma_nymphula_GLP36_PG | Z131 | DS1 |
| Zygoptera | Pyrrhosoma_nymphula_GU682172 | Z283 | NCBI |
| Zygoptera | Pyrrhosoma_nymphula_GU682191 | Z284 | NCBI |
| Zygoptera | Pyrrhosoma_nymphula_KU220874 | Z285 | NCBI |
| Zygoptera | Pyrrhosoma_nymphula_KU220878 | Z286 | NCBI |
| Zygoptera | Pyrrhosoma_nymphula_KU220879 | Z287 | NCBI |
| Zygoptera | Pyrrhosoma_nymphula_KU220880 | Z288 | NCBI |
| Zygoptera | Pyrrhosoma_nymphula_KU220881 | Z289 | NCBI |
| Zygoptera | Pyrrhosoma_nymphula_KU220882_Pyrrhosoma_nymphula_KU220883 | Z290 | NCBI |
| Zygoptera | Pyrrhosoma_nymphula_KU220884 | Z292 | NCBI |
| Zygoptera | Pyrrhosoma_nymphula_KU220885 | Z293 | NCBI |
| Anisoptera | Selysiothemis_nigra_LI26_CT | A142 | DS1 |
| Anisoptera | Selysiothemis_nigra_MIBAG0156_FE | A143 | DS1 |
| Anisoptera | Selysiothemis_nigra_MIBAG0257_CB | A144 | DS1 |
| Anisoptera | Somatochlora_alpestris_GA19_TN_Somatochlora_alpestris_IC15_UD_Somatochlora_alpestris_MIBAG0220_SO_Somatochlora_alpestris_HO01_NOR_3 | A145 | DS1 |
| Anisoptera | Somatochlora_alpestris_MIBAG0027_SO | A147 | DS1 |
| Anisoptera | Somatochlora_arctica_GA24_TN | A149 | DS1 |
| Anisoptera | Somatochlora_arctica_MIBAG0105_BG | A150 | DS1 |
| Anisoptera | Somatochlora_flavomaculata_FBAQU548_10 | A383 | BOLD |
| Anisoptera | Somatochlora_flavomaculata_FM42_LE_Somatochlora_flavomaculata_ZPLOD836-20 | A151 | DS1 / BOLD |
| Anisoptera | Somatochlora_flavomaculata_GA23_TN | A152 | DS1 |
| Anisoptera | Somatochlora_flavomaculata_GS17_VC | A153 | DS1 |
| Anisoptera | Somatochlora_meridionalis_AC35_FR_Somatochlora_meridionalis_GLP37_PG | A154 | DS1 |
| Anisoptera | Somatochlora_meridionalis_FL36_MC | A155 | DS1 |
| Anisoptera | Somatochlora_meridionalis_IC20_GO | A157 | DS1 |
| Anisoptera | Somatochlora_meridionalis_IC24_UD | A158 | DS1 |
| Anisoptera | Somatochlora_meridionalis_MIBAG0223_CN_Somatochlora_meridionalis_MIBAG0224_CN_Somatochlora_meridionalis_MIBAG0225_CN | A159 | DS1 |
| Anisoptera | Somatochlora_meridionalis_ZPLOD837-20 | A384 | BOLD |
| Anisoptera | Somatochlora_metallica_GA40_TN_Somatochlora_metallica_GA45_PV_Somatochlora_metallica_GS11_VC | A160 | DS1 |
| Anisoptera | Somatochlora_metallica_HETFI053_11_Somatochlora_metallica_HETFI055_11_Somatochlora_metallica_HETFI056_11 | A385 | BOLD |
| Anisoptera | Somatochlora_metallica_LI03_TN | A161 | DS1 |
| Zygoptera | Sympecma_fusca_ASG753_SU_Sympecma_fusca_FBAQU1426_13_Sympecma_fusca_HM901877_Sympecma_fusca_LI32_ME_Sympecma_fusca_MIBAG0163_VA_Sympecma_fusca_MIBAG0214_MO_Sympecma_fusca_SS15_TP | Z132 | DS1 / NCBI / BOLD |
| Zygoptera | Sympecma_fusca_FM02_BA_Sympecma_fusca_KF369553 | Z133 | DS1 / NCBI |
| Zygoptera | Sympecma_fusca_HM422052 | Z294 | NCBI |
| Zygoptera | Sympecma_fusca_MIBAG0177_BI | Z134 | DS1 |
| Zygoptera | Sympecma_paedisca_FBAQU1444_13 | Z295 | BOLD |
| Zygoptera | Sympecma_paedisca_KF257126 | Z296 | NCBI |
| Zygoptera | Sympecma_paedisca_MIBAG0176_BI | Z135 | DS1 |
| Zygoptera | Sympecma_paedisca_XJDQD098_18 | Z297 | BOLD |
| Zygoptera | Sympecma_paedisca_XJDQD099_18_Sympecma_paedisca_XJDQD100_18_Sympecma_paedisca_XJDQD102_18 | Z298 | BOLD |
| Zygoptera | Sympecma_paedisca_XJDQD101_18 | Z299 | BOLD |
| Zygoptera | Sympecma_paedisca_XJDQD103_18 | Z300 | BOLD |
| Zygoptera | Sympecma_paedisca_XJDQD104_18 | Z301 | BOLD |
| Anisoptera | Sympetrum_danae_DB02_AO | A163 | DS1 |
| Anisoptera | Sympetrum_danae_FBAQU516_10 | A386 | BOLD |
| Anisoptera | Sympetrum_danae_GA74_TN | A164 | DS1 |
| Anisoptera | Sympetrum_danae_JF839312_Sympetrum_danae_JN294417_Sympetrum_danae_JN294419_Sympetrum_danae_JN294422_Sympetrum_danae_JN294513 | A387 | NCBI |
| Anisoptera | Sympetrum_danae_JF839317_Sympetrum_danae_JF839318 | A388 | NCBI |
| Anisoptera | Sympetrum_danae_JN294355_Sympetrum_danae_JN294378_Sympetrum_danae_JN294409_Sympetrum_danae_JN294412_Sympetrum_danae_JN294415_Sympetrum_danae_JN294418_Sympetrum_danae_JN294425_Sympetrum_danae_KM532923 | A389 | NCBI |
| Anisoptera | Sympetrum_danae_JN294359 | A390 | NCBI |
| Anisoptera | Sympetrum_danae_JN294420 | A391 | NCBI |
| Anisoptera | Sympetrum_danae_JN294426 | A392 | NCBI |
| Anisoptera | Sympetrum_danae_JN294429 | A393 | NCBI |
| Anisoptera | Sympetrum_danae_JN294430 | A394 | NCBI |
| Anisoptera | Sympetrum_danae_JN294475 | A395 | NCBI |

| Suborder | Species name/Sample ID | Haplotype | DNA barcode source |
|---|---|---|---|
| Anisoptera | Sympetrum_danae_JN294508 | A396 | NCBI |
| Anisoptera | Sympetrum_danae_KM528837 | A397 | NCBI |
| Anisoptera | Sympetrum_depressiusculum_GA69_TO_Sympetrum_depressiusculum_MIBAG0152_BG | A165 | DS1 |
| Anisoptera | Sympetrum_flaveolum_DB23_AO_Sympetrum_flaveolum_MIBAG0119_PC | A166 | DS1 |
| Anisoptera | Sympetrum_flaveolum_MIBAG0283_AQ | A168 | DS1 |
| Anisoptera | Sympetrum_flaveolum_ZPLOD838-20 | A398 | BOLD |
| Anisoptera | Sympetrum_fonscolombii_ASG674_SS_Sympetrum_fonscolombii_MIBAG0086_PV | A169 | DS1 |
| Anisoptera | Sympetrum_fonscolombii_FBAQU552_10 | A399 | BOLD |
| Anisoptera | Sympetrum_fonscolombii_FM04_TA_Sympetrum_fonscolombii_GA15_TN | A170 | DS1 |
| Anisoptera | Sympetrum_fonscolombii_KF257098 | A400 | NCBI |
| Anisoptera | Sympetrum_meridionale_ASG745_SU | A172 | DS1 |
| Anisoptera | Sympetrum_meridionale_MG10_BA | A173 | DS1 |
| Anisoptera | Sympetrum_meridionale_MIBAG0141_MO | A174 | DS1 |
| Anisoptera | Sympetrum_meridionale_ZPLOD839-20 | A401 | BOLD |
| Anisoptera | Sympetrum_pedemontanum_FBAQU553_10 | A402 | BOLD |
| Anisoptera | Sympetrum_pedemontanum_GBMHO562_14_Sympetrum_pedemontanum_KF257095 | A403 | NCBI / BOLD |
| Anisoptera | Sympetrum_pedemontanum_GS13_AL | A175 | DS1 |
| Anisoptera | Sympetrum_pedemontanum_MIBAG0089_PV | A176 | DS1 |
| Anisoptera | Sympetrum_sanguineum_AC04_ME | A177 | DS1 |
| Anisoptera | Sympetrum_sanguineum_FBAQU518_10 | A404 | BOLD |
| Anisoptera | Sympetrum_sanguineum_FBAQU554_10 | A405 | BOLD |
| Anisoptera | Sympetrum_sanguineum_GA75_TN_Sympetrum_sanguineum_MIBAG0253_CB | A178 | DS1 |
| Anisoptera | Sympetrum_sanguineum_MIBAG0035_LC_Sympetrum_sanguineum_MIBAG0284_AQ | A179 | DS1 |
| Anisoptera | Sympetrum_sanguineum_ZPLOD840-20 | A406 | BOLD |
| Anisoptera | Sympetrum_striolatum_ASG460_SS_Sympetrum_striolatum_GA82_TN_Sympetrum_striolatum_ZPLOD841-20_Sympetrum_striolatum_GU682171_Sympetrum_striolatum_MG19_BA | A180 | DS1 |
| Anisoptera | Sympetrum_striolatum_FBAQU519_10 | A407 | BOLD |
| Anisoptera | Sympetrum_striolatum_KF257086 | A408 | NCBI |
| Anisoptera | Sympetrum_striolatum_MIBAG0116_PC | A182 | DS1 |
| Anisoptera | Sympetrum_striolatum_ZMBN328_16 | A409 | BOLD |
| Anisoptera | Sympetrum_vulgatum_FBAQU520_10_Sympetrum_vulgatum_GA33_TN_Sympetrum_vulgatum_GU682170_Sympetrum_vulgatum_LT634113_Sympetrum_vulgatum_LT634115_Sympetrum_vulgatum_LT898338_Sympetrum_vulgatum_LT898339_Sympetrum_vulgatum_MIBAG0160_LC | A183 | DS1 / NCBI / BOLD |
| Anisoptera | Sympetrum_vulgatum_FBAQU555_10 | A410 | BOLD |
| Anisoptera | Sympetrum_vulgatum_LT634114 | A411 | NCBI |
| Anisoptera | Sympetrum_vulgatum_LT634116 | A412 | NCBI |
| Anisoptera | Sympetrum_vulgatum_LT634117 | A413 | NCBI |
| Anisoptera | Sympetrum_vulgatum_LT634118 | A414 | NCBI |
| Anisoptera | Sympetrum_vulgatum_LT634119 | A415 | NCBI |
| Anisoptera | Sympetrum_vulgatum_LT634120_Sympetrum_vulgatum_LT634122_Sympetrum_vulgatum_LT634124_Sympetrum_vulgatum_LT898334_Sympetrum_vulgatum_LT898335_Sympetrum_vulgatum_LT898336 | A416 | NCBI |
| Anisoptera | Sympetrum_vulgatum_LT634121 | A417 | NCBI |
| Anisoptera | Sympetrum_vulgatum_LT634123 | A418 | NCBI |
| Anisoptera | Sympetrum_vulgatum_LT634125_Sympetrum_vulgatum_LT634126 | A419 | NCBI |
| Anisoptera | Sympetrum_vulgatum_LT898333 | A420 | NCBI |
| Anisoptera | Sympetrum_vulgatum_LT898337 | A421 | NCBI |
| Anisoptera | Trhitemis_annulata_ASG688_SS | A184 | DS1 |
| Anisoptera | Trithemis_annulata_DB12_SP_Trithemis_annulata_MIBAG0172_MI | A185 | DS1 |
| Anisoptera | Trithemis_annulata_FJ358479 | A422 | NCBI |
| Anisoptera | Trithemis_annulata_FJ358480 | A423 | NCBI |
| Anisoptera | Trithemis_annulata_KU566417_Trithemis_annulata_NSPAF295_15 | A424 | NCBI / BOLD |
| Anisoptera | Trithemis_annulata_SS20_TP | A186 | DS1 |
| Anisoptera | Zygonyx_torridus_KJ994781_Zygonyx_torridus_KU566506_Zygonyx_torridus_NSPAF216_15 | A425 | NCBI / BOLD |
| Anisoptera | Zygonyx_torridus_LC198675 | A426 | NCBI |
| Anisoptera | Zygonyx_torridus_SS31_TP | A187 | DS1 |

MOLECULAR ECOLOGY RESOURCES

**APPENDIX S3:** Support values of the species delimitation hypotheses obtained with the Bayesian implementation of the PTP and GMYC methods, for both DS1 and DS2 (n.d.: not detected in DS1).

| | Dataset 1 | | Dataset 2 | |
|---|---|---|---|---|
| | bPTP | bGMYC | bPTP | bGMYC |
| **Zygoptera** | | | | |
| *Calopteryx haemorrhoidalis* | 0.967 | 0.9-1 | 0.995 | 0.9-1 |
| *Calopteryx splendens* + *Calopteryx xanthostoma* | 0.944 | 0.9-1 | 0.996 | 0.9-1 |
| *Calopteryx virgo* | 0.899 | 0.9-1 | 0.972 | 0.9-1 |
| *Ceriagrion tenellum* | 0.929 | 0.9-1 | 0.994 | 0.9-1 |
| *Chalcolestes parvidens* + *Chalcolestes viridis* | 0.980 | 0.9-1 | 0.979 | 0.9-1 |
| *Chalcolestes viridis* | 0.780 | 0.9-1 | 0.891 | 0.9-1 |
| *Coenagrion caerulescens* | 0.975; 0.950; 0.914 | 0.7-0.8; 0.7-0.8 | 0.897 | 0.9-1; 0.9-1; 0.9-1 |
| *Coenagrion hastulatum* | 0.982 | 0.9-1 | 0.761 | 0.9-1 |
| *Coenagrion mercuriale* I | n.d. | n.d. | 0.643 | 0.9-1; 0.9-1 |
| *Coenagrion mercuriale* II | n.d. | n.d. | 0.660 | 0.9-1 |
| *Coenagrion mercuriale* III | 0.824 | 0.9-1 | 0.921 | 0.9-1 |
| *Coenagrion ornatum* + *Coenagrion puella* + *Coenagrion pulchellum* | 0.812; 0.756 | 0.7-0.8 | 0.705 | 0.8-0.9; 0.8-0.9 |
| *Coenagrion puella* | n.d. | n.d. | 0.990 | 0.9-1 |
| *Coenagrion scitulum* | 0.918 | 0.9-1 | 0.953 | 0.9-1 |
| *Enallagma cyathigerum* | 0.729 | 0.9-1 | 0.976 | 0.7-0.8 |
| *Erythromma lindenii* I | 0.825; 0.791 | 0.7-0.8 | 0.998 | 0.9-1 |
| *Erythromma lindenii* II | 0.853 | 0.9-1 | 0.956 | 0.9-1 |
| *Erythromma najas* | 0.691 | 0.8-0.9 | 0.953 | 0.8-0.9 |
| *Erythromma viridulum* | 0.890; 0.892 | 0.7-0.8 | 0.987 | 0.8-0.9 |
| *Ischnura elegans* + *Ischnura genei* | 0.762 | 0.9-1 | 0.712 | 0.9-1; 0.9-1 |
| *Ischnura pumilio* | 0.995 | 0.9-1 | 0.944 | 0.9-1 |
| *Lestes barbarus* | 0.980 | 0.9-1 | 0.994 | 0.9-1 |
| *Lestes dryas* I | n.d. | n.d. | 0.762 | 0.9-1 |
| *Lestes dryas* II | 0.724 | 0.9-1 | 0.685 | 0.9-1 |
| *Lestes macrostigma* | 1 | 0.9-1 | 1 | 0.9-1 |
| *Lestes sponsa* | 0.894 | 0.9-1 | 0.736 | 0.9-1 |
| *Lestes virens* | 0.964; 0.982 | 0.9-1; 0.9-1 | 0.764 | 0.9-1; 0.9-1 |
| *Nehalennia speciosa* | 0.984 | 0.9-1 | 0.948 | 0.9-1; 0.9-1 |
| *Platycnemis pennipes* | 0.672 | 0.9-1 | 0.904; 0.858 | 0.7-0.8 |
| *Pyrrhosoma nymphula* | 0.884 | 0.8-0.9 | 0.964 | 0.9-1; 0.9-1; 0.9-1 |
| *Sympecma fusca* | 0.714 | 0.8-0.9 | 0.993 | 0.8-0.9 |
| *Sympecma paedisca* | 1 | 0.9-1 | 0.982 | 0.8-0.9 |
| **Anisoptera** | | | | |
| *Aeshna affinis* | 0.973 | 0.9-1 | 0.993 | 0.9-1 |
| *Aeshna caerulea* | 1 | 0.9-1 | 1 | 0.9-1 |
| *Aeshna cyanea* | 0.990 | 0.9-1 | 0.771 | 0.9-1; 0.9-1 |
| *Aeshna grandis* | 0.976 | 0.9-1 | 0.747 | 0.9-1 |
| *Aeshna isoceles* | 0.886 | 0.9-1 | 0.849 | 0.9-1 |
| *Aeshna juncea* I | n.d. | n.d. | 0.541 | 0.7-0.8 |
| *Aeshna juncea* II | 0.935 | 0.9-1 | 0.522 | 0.9-1 |
| *Aeshna mixta* | 0.902 | 0.9-1 | 0.843 | 0.9-1 |
| *Aeshna subarctica* | 0.980 | 0.9-1 | 0.722 | 0.9-1 |
| *Anax ephippiger* | 0.722 | 0.9-1 | 0.906 | 0.9-1 |

| | Dataset 1 | | Dataset 2 | |
|---|---|---|---|---|
| | bPTP | bGMYC | bPTP | bGMYC |
| *Anax imperator + Anax parthenope* | 0.99 | 0.7-0.8 | 0.914 | 0.7-0.8 |
| *Boyeria irene* | 0.987 | 0.9-1 | 0.985 | 0.9-1 |
| *Brachythemis impartita* | 0.995 | 0.9-1 | 0.952 | 0.9-1 |
| *Brachytron pratense* | 0.915 | 0.9-1 | 0.960 | 0.9-1 |
| *Cordulegaster bidentata* | 0.581; 0.577 | 0.7-0.8 | 0.676 | 0.8-0.9 |
| *Cordulegaster boltonii* | 0.975 | 0.7-0.8 | 0.986 | 0.8-0.9 |
| *Cordulegaster heros* | 0.976 | 0.9-1 | 0.876 | 0.9-1 |
| *Cordulegaster trinacriae* | 0.823 | 0.9-1 | 0.815 | 0.9-1 |
| *Cordulia aenea* | 0.981 | 0.9-1 | 0.994 | 0.9-1 |
| *Crocothemis erythraea* | 0.932 | 0.9-1 | 0.780 | 0.8-0.9 |
| *Diplacodes lefebvrii* | 0.964 | 0.9-1 | 0.995 | 0.9-1 |
| *Gomphus vulgatissimus* | 0.996 | 0.9-1 | 0.998 | 0.9-1 |
| *Leucorrhinia dubia* | 0.951 | 0.9-1 | 0.989 | 0.9-1 |
| *Leucorrhinia pectoralis* | 0.961 | 0.9-1 | 1 | 0.9-1 |
| *Libellula depressa* | 0.973 | 0.9-1 | 0.994 | 0.9-1 |
| *Libellula fulva* | 0.846 | 0.9-1 | 0.990 | 0.9-1 |
| *Libellula quadrimaculata* | 0.886 | 0.7-0.8 | 0.810 | 0.8-0.9; 0.8-0.9 |
| *Lindenia tetraphylla* | 0.984 | 0.9-1 | 0.997 | 0.9-1 |
| *Onychogomphus forcipatus* I | 0.573 | 0.8-0.9 | 0.755 | 0.9-1 |
| *Onychogomphus forcipatus* II | 0.967 | 0.9-1 | 0.793 | 0.9-1 |
| *Onychogomphus uncatus* | 0.990 | 0.9-1 | 0.855 | 0.9-1 |
| *Ophiogomphus cecilia* | 1 | 0.9-1 | 1 | 0.9-1 |
| *Orthetrum albistylum* I | 1 | 0.9-1 | 1 | 0.9-1 |
| *Orthetrum albistylum* II | n.d. | n.d. | 0.999 | 0.8-0.9 |
| *Orthetrum albistylum* III | n.d. | n.d. | 0.998 | 0.8-0.9 |
| *Orthetrum brunneum* | 0.874 | 0.9-1 | 0.943 | 0.9-1 |
| *Orthetrum cancellatum* | 0.982; 0.712 | 0.9-1; 0.9-1 | 0.916 | 0.7-0.8 |
| *Orthetrum coerulescens* | 0.601; 0.591 | 0.9-1; 0.9-1 | 0.994; 0.573 | 0.7-0.8 |
| *Orthetrum nitidinerve* | 1 | 0.9-1 | 1 | 0.9-1 |
| *Orthetrum trinacria* | 0.599 | 0.7-0.8 | 0.930 | 0.9-1 |
| *Oxygastra curtisii* | 0.978 | 0.9-1 | 0.981 | 0.9-1 |
| *Paragomphus genei* | 0.948 | 0.9-1 | 0.650 | 0.8-0.9 |
| *Selysiothemis nigra* | 0.871 | 0.9-1 | 0.980 | 0.9-1 |
| *Somatochlora alpestris* | 0.984 | 0.9-1 | 0.997 | 0.9-1 |
| *Somatochlora arctica* | 0.964 | 0.9-1 | 0.992 | 0.9-1 |
| *Somatochlora flavomaculata* | 0.827 | 0.9-1 | 0.993 | 0.9-1 |
| *Somatochlora meridionalis + Somatochlora metallica* | 0.797 | 0.9-1 | 0.981 | 0.9-1 |
| *Stylurus flavipes* I | n.d. | n.d. | 0.962 | 0.9-1 |
| *Stylurus flavipes* II | 1 | 0.9-1 | 1 | 0.9-1 |
| *Sympetrum danae* I | n.d. | n.d. | 0.896 | 0.9-1 |
| *Sympetrum danae* II | 0.994 | 0.9-1 | 0.989 | 0.9-1 |
| *Sympetrum depressiusculum* | 1 | 0.9-1 | 1 | 0.9-1 |
| *Sympetrum flaveolum* | 0.848 | 0.9-1 | 0.962 | 0.9-1 |
| *Sympetrum fonscolombii* | 0.986 | 0.9-1 | 0.985 | 0.9-1 |
| *Sympetrum meridionale* | 0.960 | 0.9-1 | 0.804 | 0.9-1 |
| *Sympetrum pedemontanum* | 0.962 | 0.9-1 | 0.991; 0.955 | 0.7-0.8 |
| *Sympetrum sanguineum* | 0.931 | 0.9-1 | 0.997 | 0.9-1 |
| *Sympetrum striolatum* | 0.993 | 0.9-1 | 0.992 | 0.9-1 |
| *Sympetrum vulgatum* | 1 | 0.9-1 | 0.872 | 0.9-1 |

| | Dataset 1 | | Dataset 2 | |
|---|---|---|---|---|
| | **bPTP** | **bGMYC** | **bPTP** | **bGMYC** |
| *Trithemis annulata* | 0.937 | 0.9-1 | 0.658 | 0.9-1 |
| *Zygonyx torridus* | 1 | 0.9-1 | 0.826 | 0.9-1 |

MOLECULAR ECOLOGY
RESOURCES

**APPENDIX S4:** Distributions of pairwise genetic distances (K2P) obtained with ABGD for a) Zygoptera DS1, b) Anisoptera DS1, c) Zygoptera DS2, and d) Anisoptera DS2.

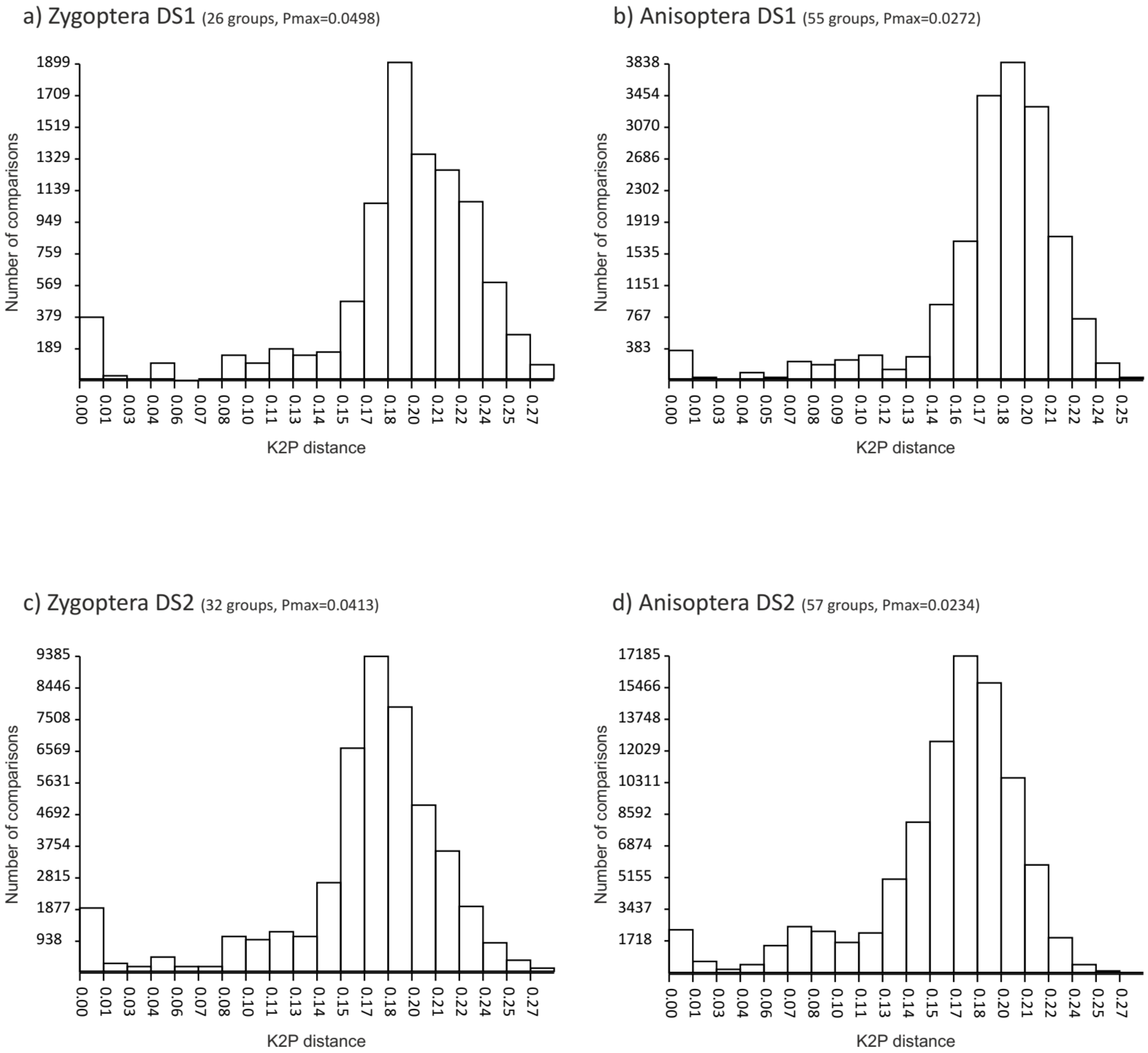

MOLECULAR ECOLOGY RESOURCES

**APPENDIX S5:** Barcode Gap Analysis of DS1 for Anisoptera and Zygoptera respectively, generated by BOLD. Three scatterplots are provided to confirm the existence and magnitude of the Barcode Gap. For each suborder, the first two scatterplots show the overlap of the max and mean intra-specific distances vs the inter-specific (nearest neighbour) distances. The third scatterplot plots the number of individuals in each species against their max intra-specific distances, as a test for sampling bias. Distance Model: Kimura 2 Parameter, Deletion Method: Pairwise Deletion, Alignment: BOLD Aligner (Amino Acid based HMM), Filters Applied: ≥ 500 bp only.

MOLECULAR ECOLOGY RESOURCES

### Barcode Gap Analysis Result BOLD SYSTEMS (PROJECT ZPLOD)

#### ANISOPTERA

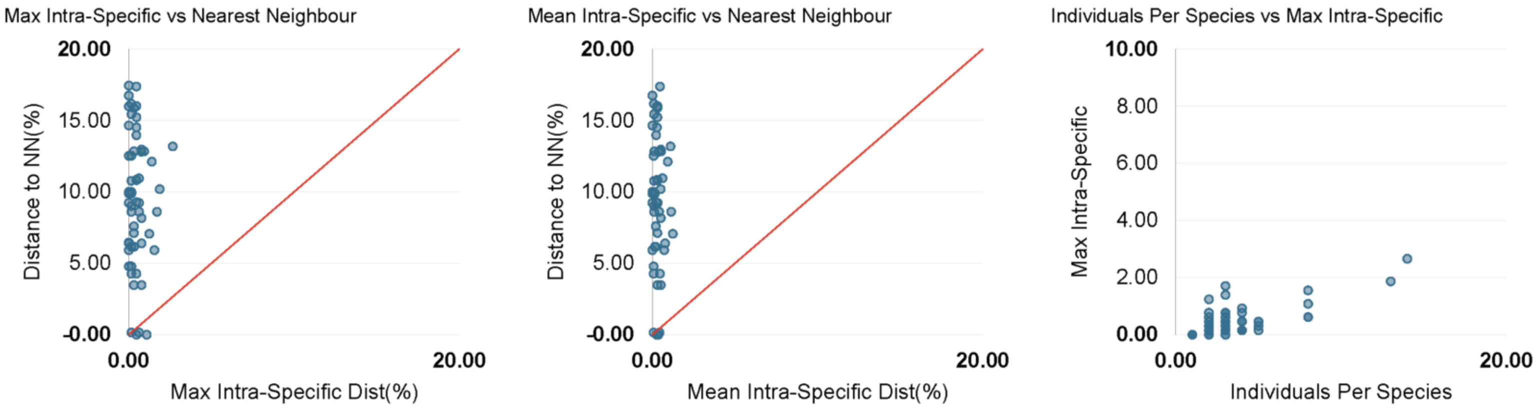


#### ZYGOPTERA

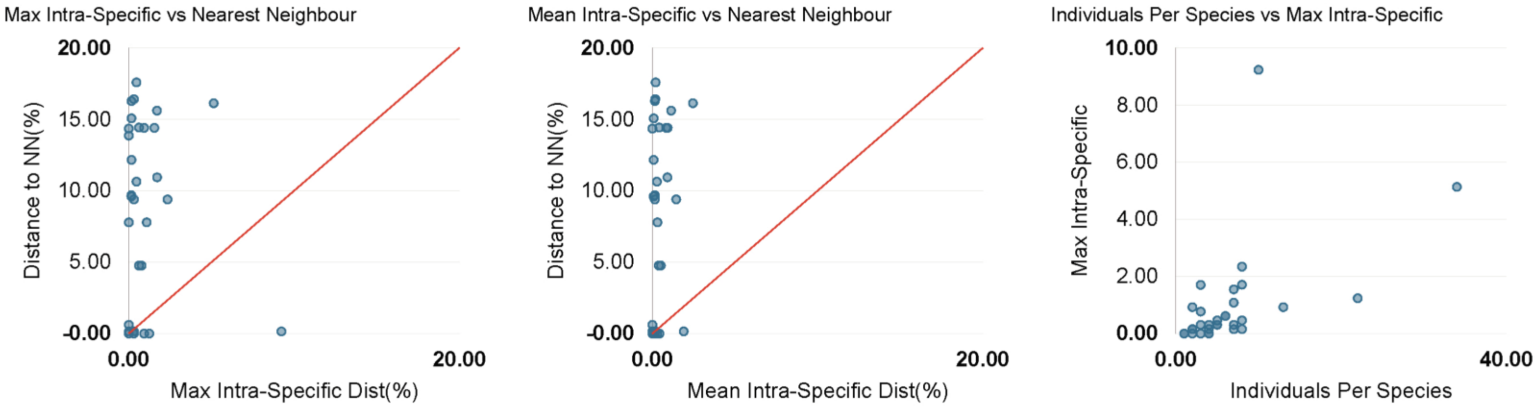

MOLECULAR ECOLOGY RESOURCES

**APPENDIX S7:** Average genetic p-distance divergence and Standard Deviation values among the geographic population of *Coenagrion mercuriale* (ITA: Italy, EUR: Europe excluding Italy, NAF: North Africa) and among three species belonging to *Coenagrion* (ORN: *ornatum*, PUL: *pulchellum*, PUE: *puella*) calculated at two mitochondrial (16s rDNA and COI) and three nuclear (AgT, PRMT and MLC) markers. Sequence data were retrieved from GenBank. Concerning COI, also the sequences produced in this study were used to calculate the genetic distances. The exclusiveness of haplotypes belonging to the Italian populations of *C. mercuriale* and to the other three *Coenagrion* species is indicated for each genetic marker. Genetic distance values have been calculated using MEGA X (Kumar et al., 2018).

**Geographic genetic divergence of *Coenagrion mercuriale***

| locus | ITA-EUR | ITA-NAF | EUR-NAF | Italian Exclusive haplotypes? |
|---|---|---|---|---|
| **16s rDNA** | 0.019 (0.006) | 0.016 (0.005) | 0.008 (0.004) | yes |
| **COI** | 0.059 (0.009) | 0.057 (0.009) | 0.043 (0.008) | yes |
| **AgT** | 0.030 (0.007) | 0.023 (0.007) | 0.0036 (0.005) | yes |
| **PRMT** | 0.012 (0.004) | 0.016 (0.004) | 0.016 (0.004) | yes |
| **MLC** | 0.035 (0.012) | 0.029 (0.009) | 0.031 (0.009) | yes |

**Interspecific genetic divergence of *Coenagrion puella, C. pulchellum* and *C. ornatum***

| locus | ORN-PUE | ORN-PUL | PUE-PUL | Species exclusive haplotypes? |
|---|---|---|---|---|
| **16s rDNA** | 0 (0) | 0 (0) | 0 (0) | no |
| **COI** | 0.0039 (0.0014) | 0.0062 (0.0019) | 0.0048 (0.0016) | no |
| **AgT** | 0.0423 (0.0096) | 0.0664 (0.0129) | 0.0563 (0.0105) | yes |
| **PRMT** | 0.0163 (0.0046) | 0.0148 (0.0042) | 0.0129 (0.039) | yes |
| **MLC** | 0.0325 (0.0091) | 0.0344 (0.0085) | 0.0276 (0.0072) | yes |
