## Appendix S6 for "Italian Odonates in the Pandora’s Box: A Comprehensive DNA Barcoding Inventory Shows Taxonomic Warnings at the Holarctic Scale"

MOLECULAR ECOLOGY RESOURCES

**APPENDIX S6:** Multi-approach species delimitation of the 31 Zygoptera and 57 Anisoptera species investigated in this study based on Holarctic COI DNA barcode sequences from DS2. A Bayesian tree is used as a base to summarize the two threshold-based (OT and ABGD) and four character explicit-based (PTP, MPTP, GMYC, bGMYC) approaches. Specimen haplotype identifiers are reported on tips (see Appendix S1 and S2 for further details) and the number of specimens sharing each haplotype is reported within brackets. Numbers above nodes represent BPP and BS, respectively, whereas asterisks correspond to maximal node support (BPP $\geq$ 0.99 and BS $\geq$ 95). Vertical colored solid boxes delimit putative species identified by the different approaches. Black: delimitation congruent with the identified morphospecies; Red: intraspecific delimitation; Light Blue: no interspecific delimitation; Yellow: Mixed species delimitation. In case of intraspecific delimitation, each lineage has been numbered with Roman numerals. This tree was created with BEAST 1.8.2.

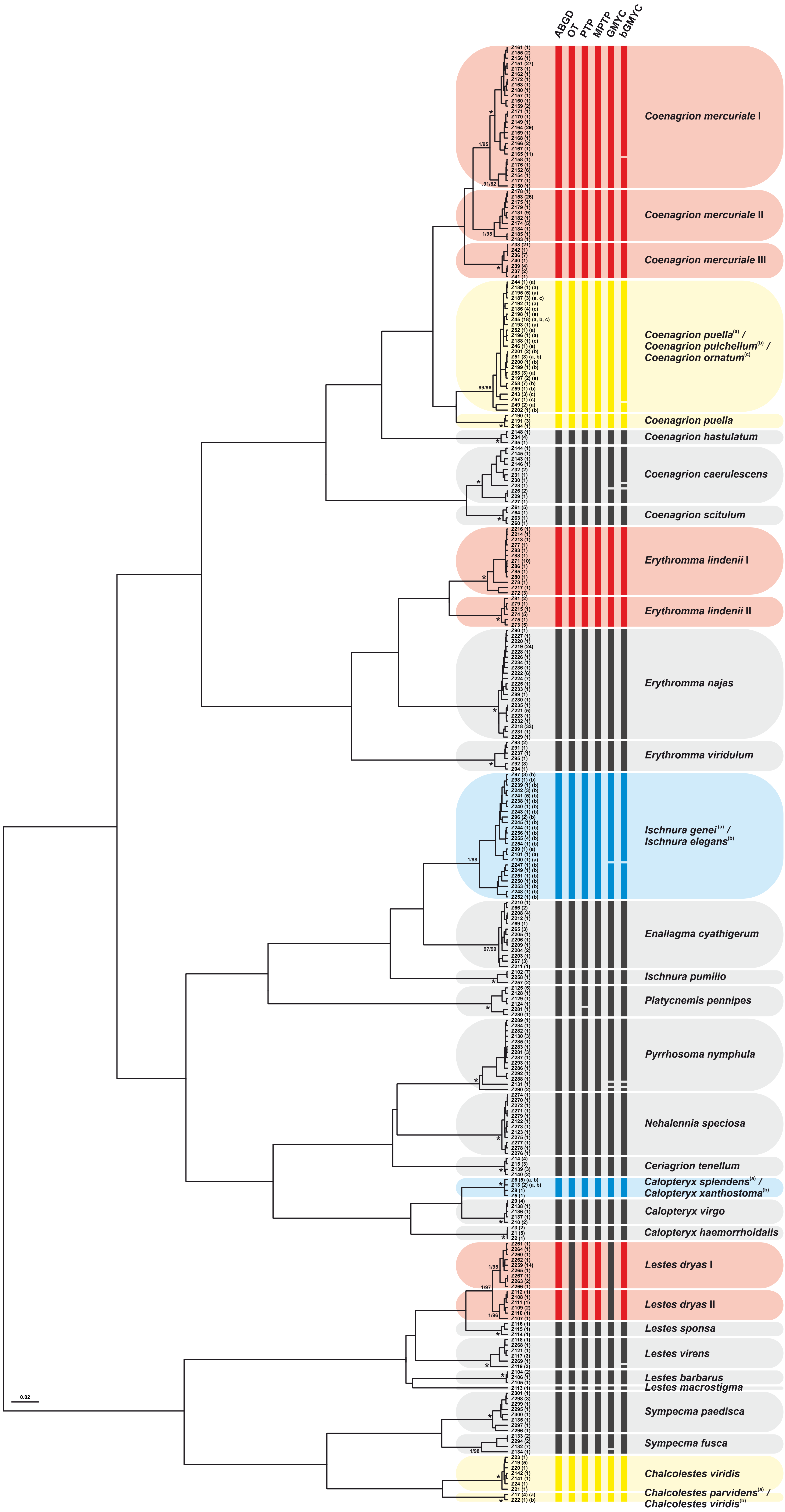

ABGD
OT
PTP
MPTP
GMYC
bGMYC
Coenagrion mercuriale I
Coenagrion mercuriale II
Coenagrion mercuriale III
Coenagrion puella(a) / Coenagrion pulchellum(b) / Coenagrion ornatum(c)
Coenagrion puella
Coenagrion hastulatum
Coenagrion caerulescens
Coenagrion scitulum
Erythromma lindenii I
Erythromma lindenii II
Erythromma najas
Erythromma viridulum
Ischnura genei(a) / Ischnura elegans(b)
Enallagma cyathigerum
Ischnura pumilio
Platycnemis pennipes
Pyrrhosoma nymphula
Nehalennia speciosa
Ceriagrion tenellum
Calopteryx splendens(a) / Calopteryx xanthostoma(b)
Calopteryx virgo
Calopteryx haemorrhoidalis
Lestes dryas I
Lestes dryas II
Lestes sponsa
Lestes virens
Lestes barbarus
Lestes macrostigma
Sympecma paedisca
Sympecma fusca
Chalcolestes viridis
Chalcolestes parvidens(a) / Chalcolestes viridis(b)
0.02

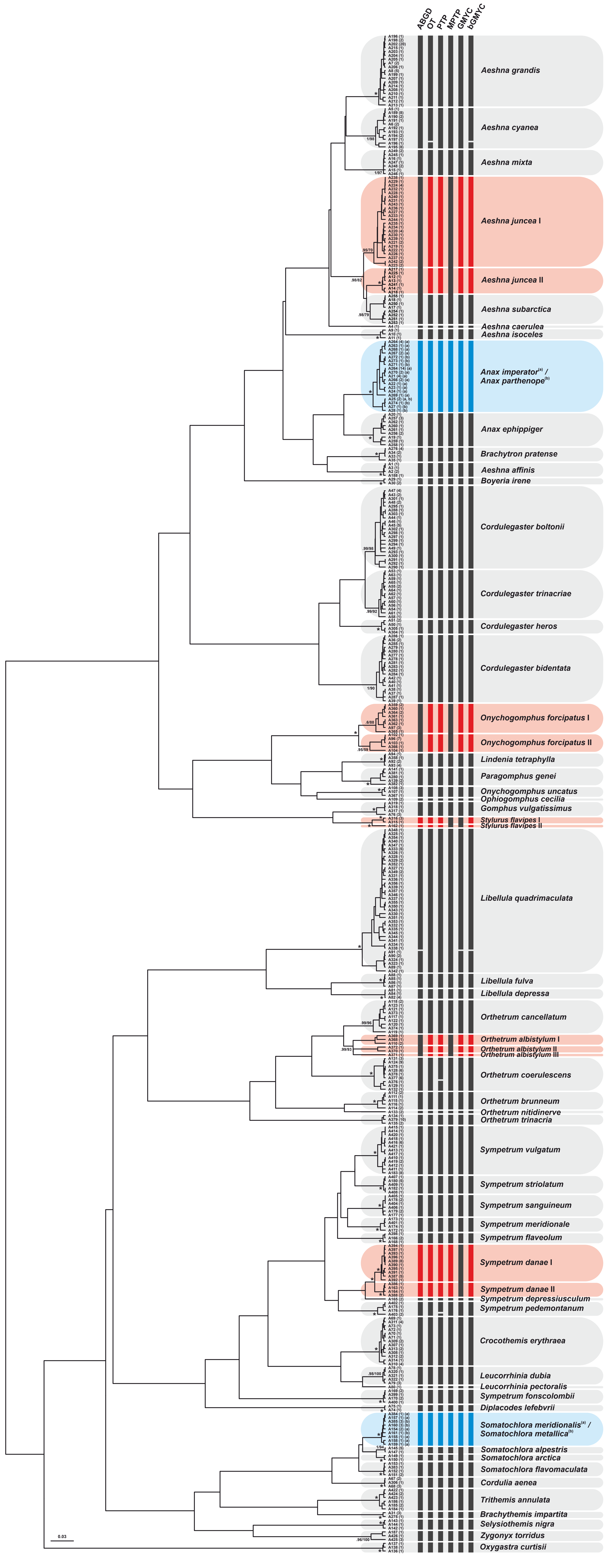
ABGD
OT
PTP
MPTP
GMYC
bGMYC
Aeshna grandis
Aeshna cyanea
Aeshna mixta
Aeshna juncea I
Aeshna juncea II
Aeshna subarctica
Aeshna caerulea
Aeshna isoceles
Anax imperator[(a)] /
Anax parthenope[(b)]
Anax ephippiger
Brachytron pratense
Aeshna affinis
Boyeria irene
Cordulegaster boltonii
Cordulegaster trinacriae
Cordulegaster heros
Cordulegaster bidentata
Onychogomphus forcipatus I
Onychogomphus forcipatus II
Lindenia tetraphylla
Paragomphus genei
Onychogomphus uncatus
Ophiogomphus cecilia
Gomphus vulgatissimus
Stylurus flavipes I
Stylurus flavipes II
Libellula quadrimaculata
Libellula fulva
Libellula depressa
Orthetrum cancellatum
Orthetrum albistylum I
Orthetrum albistylum II
Orthetrum albistylum III
Orthetrum coerulescens
Orthetrum brunneum
Orthetrum nitidinerve
Orthetrum trinacria
Sympetrum vulgatum
Sympetrum striolatum
Sympetrum sanguineum
Sympetrum meridionale
Sympetrum flaveolum
Sympetrum danae I
Sympetrum danae II
Sympetrum depressiusculum
Sympetrum pedemontanum
Crocothemis erythraea
Leucorrhinia dubia
Leucorrhinia pectoralis
Sympetrum fonscolombii
Diplacodes lefebvrii
Somatochlora meridionalis[(a)] /
Somatochlora metallica[(b)]
Somatochlora alpestris
Somatochlora arctica
Somatochlora flavomaculata
Cordulia aenea
Trithemis annulata
Brachythemis impartita
Selysiothemis nigra
Zygonyx torridus
Oxygastra curtisii
0.03
